# Broad Brain Networks Support Curiosity-Motivated Incidental Learning Of Naturalistic Dynamic Stimuli With And Without Monetary Incentives

**DOI:** 10.1101/2022.10.04.510790

**Authors:** Stefanie Meliss, Carien van Reekum, Kou Murayama

## Abstract

Curiosity – the intrinsic desire to know – is a concept central to the human mind and knowledge acquisition. Experimental studies on information-seeking have found that curiosity facilitates memory encoding and exhibits similar reward,ng properties as extrinsic rewards/incentives by eliciting a dopaminergic response. However, it is not clear whether these findings hold with more naturalistic dynamic stimuli and how the joint effect of curiosity and extrinsic incentive manifests in learning and neural activation patterns. Herein, we presented participants with videos of magic tricks across two behavioural (N_1_ = 77, N_2_ = 78) and one fMRI study (N = 50) and asked them to rate subjective feelings of curiosity, while also performing a judgement task that was incentivised for the half of participants. Incidental memory for the magic trick was tested a week later. The integrated results showed that both curiosity and availability of extrinsic incentives enhanced encoding but did not interact with each other. However, exploratory analyses showed that curiosity and monetary incentives were associated with recollection and familiarity differently, suggesting the involvement of different encoding mechanisms. Analysis of the fMRI data using the intersubject synchronisation framework showed that, while the effects of curiosity on memory were located in the hippocampus and dopaminergic brain areas, neither the effects of curiosity nor incentives themselves were found in the often-implicated reward network, but instead were associated with cortical areas involved in processing uncertainly and attention. These results suggest that curiosity recruits broader brain networks than what was implicated in the previous literature when investigated with dynamic stimuli.

## Highlights

- New paradigm developed to investigate incentive- and curiosity-motivated learning
- Dynamic stimuli - videos of magic tricks - used to elicit curiosity
- Curiosity-motivated learning linked to broadly distributed brain regions

## 1 Introduction

> ‘We find ourselves in a bewildering world. We want to make sense of what we see around us and to ask: what is the nature of the universe? What is our place in it and where did it and we come from? Why is it the way it is?’ (Hawking, 2016, p. 205)

With the words above, Stephen Hawking introduced the concluding chapter of his famous book ‘A Brief History Of Time’ where he aimed to explain our universe to a non-scientific audience. The wonder the words capture, the intrinsic desire to know, has not only motivated scientists to dedicate their careers to trying to find answers to the big questions of the universe, but also the readers of the more than 10 million copies sold to spend their time and monetary resources to acquire knowledge about the Big Bang. This is intriguing because, arguably for most of them, being able to understand how to combine weak and strong nuclear forces with those of gravity and electromagnetism into a single unified theory will have no instrumental value to maximise their rewards in their everyday lives.

In line with this anecdotal evidence, research has found that humans actively engage in non-instrumental information-seeking (Kobayashi et al., 2019; van Lieshout et al., 2021), even if it requires a small cost (Bennett et al., 2016; Brydevall et al., 2018; Kang et al., 2009; Kobayashi & Hsu, 2019; Marvin & Shohamy, 2016; van Lieshout et al., 2018), involves taking the risk of receiving an electric shock (Lau et al., 2020), or leads to experiencing negative emotions like regret (FitzGibbon et al., 2021). These observations have led researchers to propose that information is a reward (FitzGibbon et al., 2020; Marvin & Shohamy, 2016), functioning like extrinsic rewards (e.g., food or money) to govern our behaviour. In fact, in monkeys the same dopaminergic neurons in the midbrain that signal the expected amount of primary extrinsic rewards also signal the expectation of information (Bromberg-Martin & Hikosaka, 2009). Likewise, in humans, the subjective value of information and basic extrinsic rewards share a common neural code expressed in the striatum and other reward-related areas such as the ventromedial prefrontal cortex (Kang et al., 2009; Kobayashi & Hsu, 2019; Lau et al., 2020).

### 1.1 Curiosity-Motivated Learning

The subjective feelings underlying our desire to know - which we will refer to as the subjective feeling of *curiosity* - have been shown to facilitate memory encoding (for recent reviews, see Gruber et al., 2019; Gruber & Ranganath, 2019). More specifically, the subjective feeling of curiosity elicited by a curiosity-triggering cue (i.e., a trivia question; cf. Jepma et al., 2012) facilitates the intentional encoding (Duan et al., 2020; Halamish et al., 2019) of the target item (i.e., the answer to trivia question; cf. Jepma et al., 2012). The same curiosity effects have also been found in incidental encoding paradigms after short (Brod & Breitwieser, 2019; Galli et al., 2018; Gruber et al., 2014; Jepma et al., 2012; Ligneul et al., 2018; Mullaney et al., 2014; Murphy, Dehmelt, et al., 2021; Poh et al., 2021; Stare et al., 2018) and long (Fastrich et al., 2018; Gruber et al., 2014; Kang et al., 2009; Marvin & Shohamy, 2016; Murayama & Kuhbandner, 2011; Stare et al., 2018; Swirsky et al., 2021) delays between encoding and retrieval. Interestingly, incidental information that is semantically unrelated to the cue eliciting the feeling of curiosity but presented in close temporal proximity (i.e., during a state of high compared to low curiosity) is also preferably encoded (Galli et al., 2018; Gruber et al., 2014; Murphy, Dehmelt, et al., 2021; Stare et al., 2018).

Neuroimaging research has suggested that such curiosity-motivated learning is related to the activity and interaction between three brain areas: the nucleus accumbens (NAcc), the dopaminergic midbrain (VTA/SN), and the hippocampus (HPC). Specifically, Gruber and colleagues (2014) investigated whether the brain activity during curiosity elicitation at cue presentation (i.e., the trivia question) predicts later memory for the upcoming target information (i.e., the answer to the trivia question) and found that while the dopaminergic midbrain was more activated during the anticipation of later remembered compared to later forgotten targets irrespective of the degree of curiosity elicitation, the right HPC and the bilateral NAcc showed increased activation for remembered compared to forgotten targets only for high-, but not low-curiosity cues. They also found a strong correlation between the curiosity-driven memory benefit for incidental information and the curiosity-related subsequent memory effects in the VTA/SN and the HPC as well as the functional connectivity between them for high, but not low curiosity trials. Taken together, the results suggest that anticipatory activity in the mesolimbic dopaminergic circuit and the HPC supports the learning benefits associated with high compared to low states of curiosity (Gruber & Ranganath, 2019).

However, despite the increasing number of work on curiosity-motivated learning, the vast majority of studies have relied only on a single type of material (for exceptions, see e.g., Cen et al., 2021; Jepma et al., 2012) — trivia questions (e.g., Fastrich et al., 2018; Gruber et al., 2014; Kang et al., 2009; Marvin & Shohamy, 2016; Murayama & Kuhbandner, 2011; Wade & Kidd, 2019). Although the trivia question paradigm has obvious benefits as the paradigm allows researchers to examine memory in a similar manner to traditional memory experiments (e.g., questions are ‘cues’ and answers are ‘targets’), we identify two issues.

First, the paradigm examines memory encoding by relying on discrete, static elements lacking the complex, contextual and narrative nature of everyday events (Shamay-Tsoory & Mendelsohn, 2019). In fact, the actual process of curiosity elicitation is more dynamic. In classrooms, for example, students’ curiosity ebbs and flows over time while watching the lecture, and various factors contribute to the temporal dynamics (Hidi & Renninger, 2006). In other words, the state of curiosity should not be seen as a snapshot phenomenon, but as embedded within the sequence of events and psychological processes (Murayama, 2022). Thus, it is important to induce curiosity using more complex, dynamic stimuli to ensure ecological validity.

Second, while trivia questions trigger curiosity by promoting the detection of a gap in people’s knowledge (i.e., information-based prediction errors; Gruber & Ranganath, 2019), there has been increasing consensus that curiosity is elicited by multiple different factors, which may be governed by different psychological and neural mechanisms (Gruber & Ranganath, 2019; Jach et al., 2022; Kobayashi et al., 2019; Sharot & Sunstein, 2020). The heavy reliance on trivia questions may have researchers overlook some important neural mechanisms underlying curiosity and our information-seeking behaviour. For example, the subjective feeling of curiosity can also be elicited in novel environments or when events violate expectations and create a sense of surprise (i.e., context-based prediction errors; Gruber & Ranganath, 2019). In fact, research has shown that violation of expectations stimulates surprise, curiosity, and learning (Brod et al., 2018; Brod & Breitwieser, 2019); and that surprise is a reliable predictor of curiosity (Vogl et al., 2019). More so, Ligneul and colleagues (2018) showed that surprise mediated the effects of curiosity on memory. The surprise signal itself originated in the ventromedial prefrontal cortex (vmPFC) that was also more strongly activated during subsequently remembered items. Despite such intriguing preliminary findings, the role of this surprise-based curiosity on memory encoding has been under-examined, and little research has addressed the neural correlates underlying it.

### 1.2 Role Of Extrinsic Incentives

Another critical issue is the role of extrinsic incentives and rewards^1^ in curiosity-motivated learning. Overall, the facilitating effects of curiosity on memory encoding bear a striking resemblance to the effects of extrinsic rewards on memory in the literature (for a review, see Miendlarzewska et al., 2016): it has been shown that providing monetary incentives and rewards not only increases intentional encoding of incentivized items (Adcock et al., 2006; Gruber et al., 2013; Gruber & Otten, 2010; Wolosin et al., 2012), but also the incidental encoding of information presented in the context of rewarded task (Bunzeck et al., 2010, 2012; Gruber et al., 2016; Murayama & Kitagami, 2014; Murty & Adcock, 2014; Patil et al., 2017; Stanek et al., 2019; Wittmann et al., 2005, 2008, 2011). Neuroimaging studies have linked this behavioural incentive effect on intentional encoding to activity in NAcc, HPC and VTA showing an enhanced activity during cue presentation for later remembered compared to forgotten targets only in the context of high, but not low rewards (Adcock et al., 2006) and further showed that functional connectivity between HPC and VTA/SN supports the behavioural reward effect (Adcock et al., 2006; Wolosin et al., 2012). This involvement of VTA/SN and HPC is consistent with the hypothesis that reward promotes memory formation via dopamine release modulating hippocampal synaptic encoding processes during long-term potentiation (Lisman et al., 2011; Lisman & Grace, 2005; Shohamy & Adcock, 2010).

While the effects of monetary incentives/rewards and, more recently, curiosity have been studied in isolation leading to valuable insights, only a small portion of studies have actually looked at their conjunction. Studying both effects in the same study is necessary to closely understand the similarities and differences of neural mechanisms in how they benefit learning. Murayama and Kuhbander (2011) found that both monetary reward and the interestingness of trivia questions (as rated by a separate sample) had an enhancing effect on encoding, but the main effects were further qualified by an interaction, where monetary rewards only enhanced encoding of trivia questions rated as not interesting. The findings were replicated in younger and older adults (Swirsky et al., 2021) although some other studies failed to find the interaction effects (Duan et al., 2020; Halamish et al., 2019). Thus, the literature suggests the possibility that there may be unique non-additive neural patterns when both curiosity and monetary incentives are present.

### 1.3 Current Research

The current study aims to expand our understanding of curiosity-motivated learning in two substantive manners. First, to capture the dynamic nature of curiosity, we used novel naturalistic stimuli that strongly trigger curiosity — videos of magic tricks. Magic tricks induce curiosity independent of language and prior knowledge by showing implausible if not impossible events (Kuhn et al., 2008; Rensink & Kuhn, 2014). Because the viewer generates predictions as the magic trick unfolds, any violation of causal relationships would also violate the viewer’s expectations, triggering a relatively strong surprise-based curiosity (i.e., context-based or perceptual prediction error; Zacks et al., 2007). Indeed, previous research has shown that magic tricks are rated as surprising, violate cause and effect relations, and lead to unexpected outcomes (Danek et al., 2015; Parris et al., 2009), trigger epistemic emotions (surprise in response to the trick, interest in the trick, and curiosity in the solution; Ozono et al., 2021), and elicit curiosity to motivate risky decision-making in a similar way as do trivia questions, supported by the ventral striatum (Lau et al., 2020). Intriguingly, the ventral striatum and the caudate nucleus (CN) have also been linked to the violation of expectations seen in magic tricks (Danek et al., 2015), the elicitation (Gruber et al., 2014; Kang et al., 2009) and relief of curiosity (Ligneul et al., 2018), as well as the effects of curiosity on memory within the trivia question paradigm (Duan et al., 2020).

Second, we manipulated the availability of extrinsic incentives such that we can examine the potential interactive effects of curiosity and extrinsic incentives on learning. As indicated earlier, despite the strong suggestion that information-seeking is driven by reward learning, neuroimaging studies on motivated learning examined curiosity and extrinsic incentives rather separately, making it difficult to understand how these two types of motivating factors enhance (or do not enhance) memory altogether. The current study provides a first attempt to examine the interactive effect using fMRI.

We conducted three studies (two behavioural, one fMRI) which have a similar structure. In all experiments, participants viewed a series of magic trick videos and performed a judgement task including curiosity ratings. To examine the effects of extrinsic incentives, half of the participants were promised additional monetary bonus payments for the judgement task whereas the other half of the participants did not receive such instructions. A week later, the memory for the magic tricks was assessed using surprise recognition and recall tests. Based on the previous literature, we hypothesised that both curiosity and monetary incentives would facilitate memory encoding, both of which may be supported by similar neural processes located in the hippocampal-VTA loop (Lisman & Grace, 2005), hence we also examined the potential interaction between curiosity and monetary incentives.

## 2 Methods

### 2.1 Study 1: Behavioural Study

#### 2.1.1 Participants & Design

The *a priori* defined intended sample size was in total 80 participants. This was mainly limited by the budget, but our sensitivity analysis showed that this sample size is sufficient to detect medium-sized effects at 80% of power for the between-subjects effect of monetary incentives (*d* = 0.63). Given that the reward effects on memory have been established in the literature (Adcock et al., 2006; Gruber et al., 2016; Miendlarzewska et al., 2016; Wittmann et al., 2005), we decided to go with this sample size. Participants were recruited using Prolific (https://prolific.co) for an online study consisting of two parts spaced one week apart. Both parts took approximately 45min each and participants were reimbursed with a total of £7.50 for their time. For inclusion, the following criteria were defined: age between 18 and 37, fluency in English, a minimum approval rate of 95%, and at least ten previous submissions.

Unbeknownst to the participants, the study included a between-group incentive manipulation where the experimental group was instructed that they could earn additional monetary bonus payments for their performance in the judgement task whereas the control group did not receive such instructions. The amount of the bonus was defined as £0.10 per correct answer in the judgement task. By incentivizing performance in the judgement task rather than in the memory assessment, our task examines the effects of monetary incentives on incidental encoding.

Considering potential attrition, we oversampled participants against the predefined sample size. In total, we received data from 47 and 44 participants in the control and incentives condition, respectively, out of which five and three participants were excluded due to incomplete data. All 83 participants who had submitted complete datasets were invited to participate in the second part of the study and 42 and 39 participants in the control and incentive group responded. In total, four datasets were excluded from the second part (3 due to incomplete data and 1 due to a self-reported age below 18, all from the control condition). The final sample size included in the analysis included N = 77 participants (n_control_ = 38, n_incentive_ = 39). The participant characteristics are described in Table 1. The study was reviewed and approved by the University of Reading’s School Research Ethics Committee (SREC; 2016-109-KM).

**Table 1.**
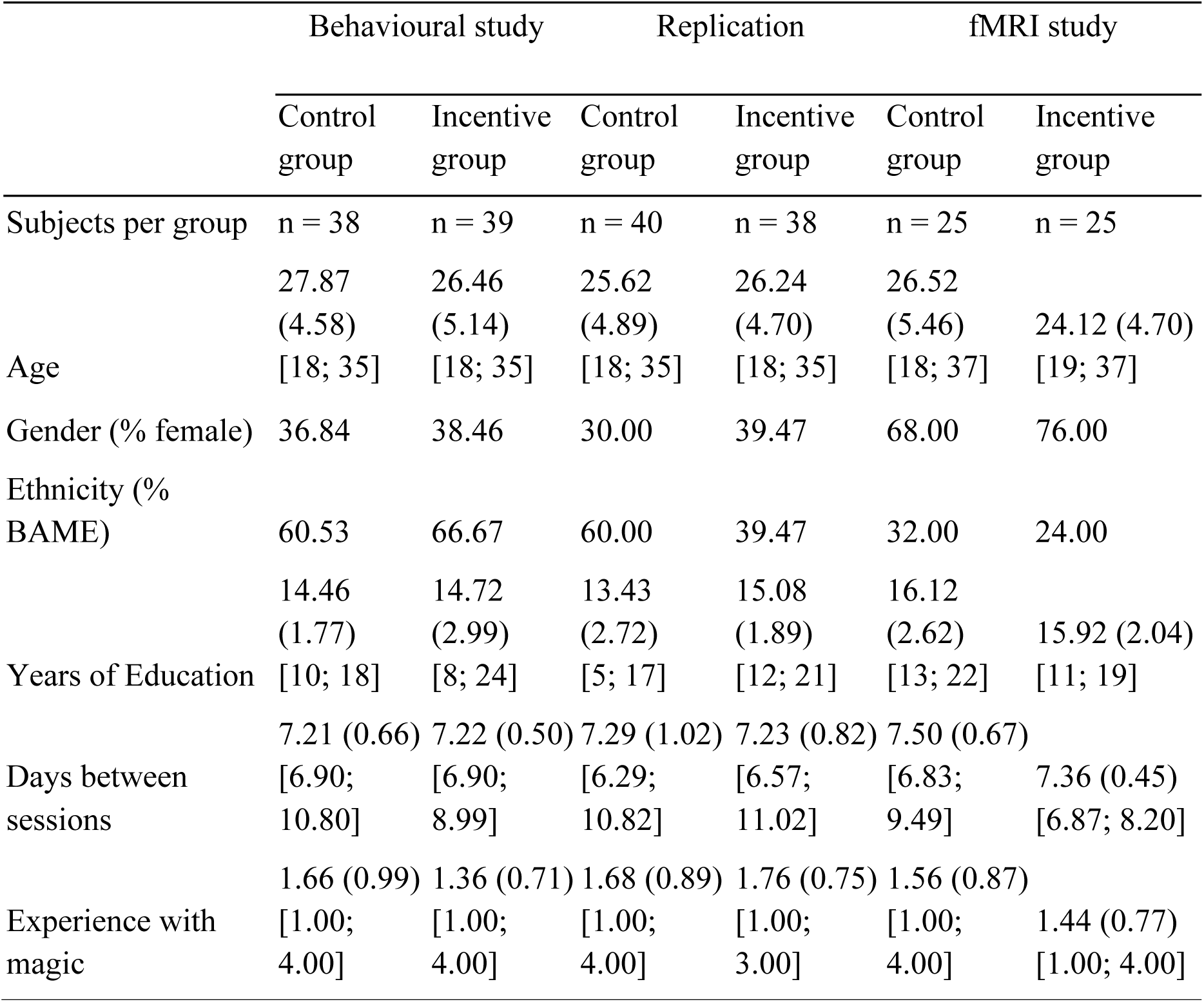
Participant Characteristics. *Note*. For interval-scaled variables, the table shows the mean (standard deviation) [minimum; maximum] separately for each group and data collection. Experience with magic tricks relates to the participant’s rating of their experience in producing magic tricks on a scale from 1 = ‘never’ to 6 = ‘very frequently’.

|  | Behavioural study |  | Replication |  | fMRI study |  |
| --- | --- | --- | --- | --- | --- | --- |
|  | Control group | Incentive group | Control group | Incentive group | Control group | Incentive group |
| Subjects per group | n = 38 | n = 39 | n = 40 | n = 38 | n = 25 | n = 25 |
| Age | 27.87<br>(4.58)<br>[18; 35] | 26.46<br>(5.14)<br>[18; 35] | 25.62<br>(4.89)<br>[18; 35] | 26.24<br>(4.70)<br>[18; 35] | 26.52<br>(5.46)<br>[18; 37] | 24.12 (4.70)<br>[19; 37] |
| Gender (% female) | 36.84 | 38.46 | 30.00 | 39.47 | 68.00 | 76.00 |
| Ethnicity (% BAME) | 60.53 | 66.67 | 60.00 | 39.47 | 32.00 | 24.00 |
| Years of Education | 14.46<br>(1.77)<br>[10; 18] | 14.72<br>(2.99)<br>[8; 24] | 13.43<br>(2.72)<br>[5; 17] | 15.08<br>(1.89)<br>[12; 21] | 16.12<br>(2.62)<br>[13; 22] | 15.92 (2.04)<br>[11; 19] |
| Days between sessions | 7.21 (0.66)<br>[6.90; 10.80] | 7.22 (0.50)<br>[6.90; 8.99] | 7.29 (1.02)<br>[6.29; 10.82] | 7.23 (0.82)<br>[6.57; 11.02] | 7.50 (0.67)<br>[6.83; 9.49] | 7.36 (0.45)<br>[6.87; 8.20] |
| Experience with magic | 1.66 (0.99)<br>[1.00; 4.00] | 1.36 (0.71)<br>[1.00; 4.00] | 1.68 (0.89)<br>[1.00; 4.00] | 1.76 (0.75)<br>[1.00; 3.00] | 1.56 (0.87)<br>[1.00; 4.00] | 1.44 (0.77)<br>[1.00; 4.00] |

#### 2.1.2 Material

We displayed short magic trick videos to participants. The magic trick videos were selected from the Magic Curiosity Arousing Tricks (MagicCATs) stimulus collection (Ozono et al., 2021). This collection was developed specifically for fMRI experiments, containing 166 magic tricks that were edited to achieve a similar background and viewing focus, and muted purposefully to minimise the effects of verbal interference. To select magic tricks used here, the following criteria were applied: (1) duration between 20 and 60s, (2) broad range of different materials and features so that magic tricks are distinguishable in a cued recall paradigm, (3) varying degrees of curiosity ratings as reported in the database, and (4) understandable without the use of subtitles. Additional editing was performed using Adobe^Ⓡ^ Premiere Pro CC^Ⓡ^ (2015) software where needed, for instance, to remove subtitles. Magic tricks were exported in a slightly larger size than available in the database (1280×720 pixels). In total, 36 magic tricks were displayed in the experiment and an additional two were used for practice trials. This number is equivalent to what has been used previously when studying decision-making using magic tricks (Lau et al., 2020). Please see Meliss et al. (2022) for more information about the magic tricks used.

A frame of each magic trick video was extracted as a cue image (1920×1080 pixels) for the memory test. For this, a frame was selected from before the moment(s) of surprise (i.e., moments violating one’s expectations) that was distinctive enough to cue the magic trick without revealing it entirely.

#### 2.1.3 Tasks & Measurements

##### Magic Trick Watching Task

During each trial of the magic trick watching task (see Figure 1, upper half), participants watched a magic trick video and were then asked to estimate how many people (out of 100) are able to correctly figure out the solution. For this, participants could choose out of the following answer options: ‘0 - 10%’, ‘11 - 20%’, ‘21 - 30 %’, and ‘31 % and more’. Afterwards, participants were asked to rate how curious they were while watching the magic trick on a 7-point Likert scale (1 = ‘not curious at all’, 7 = ‘very curious’). Importantly, the estimate rating was combined with the between-subject incentive manipulation. The incentive manipulation was part of the task instructions, which is described below.

**Figure 1.**
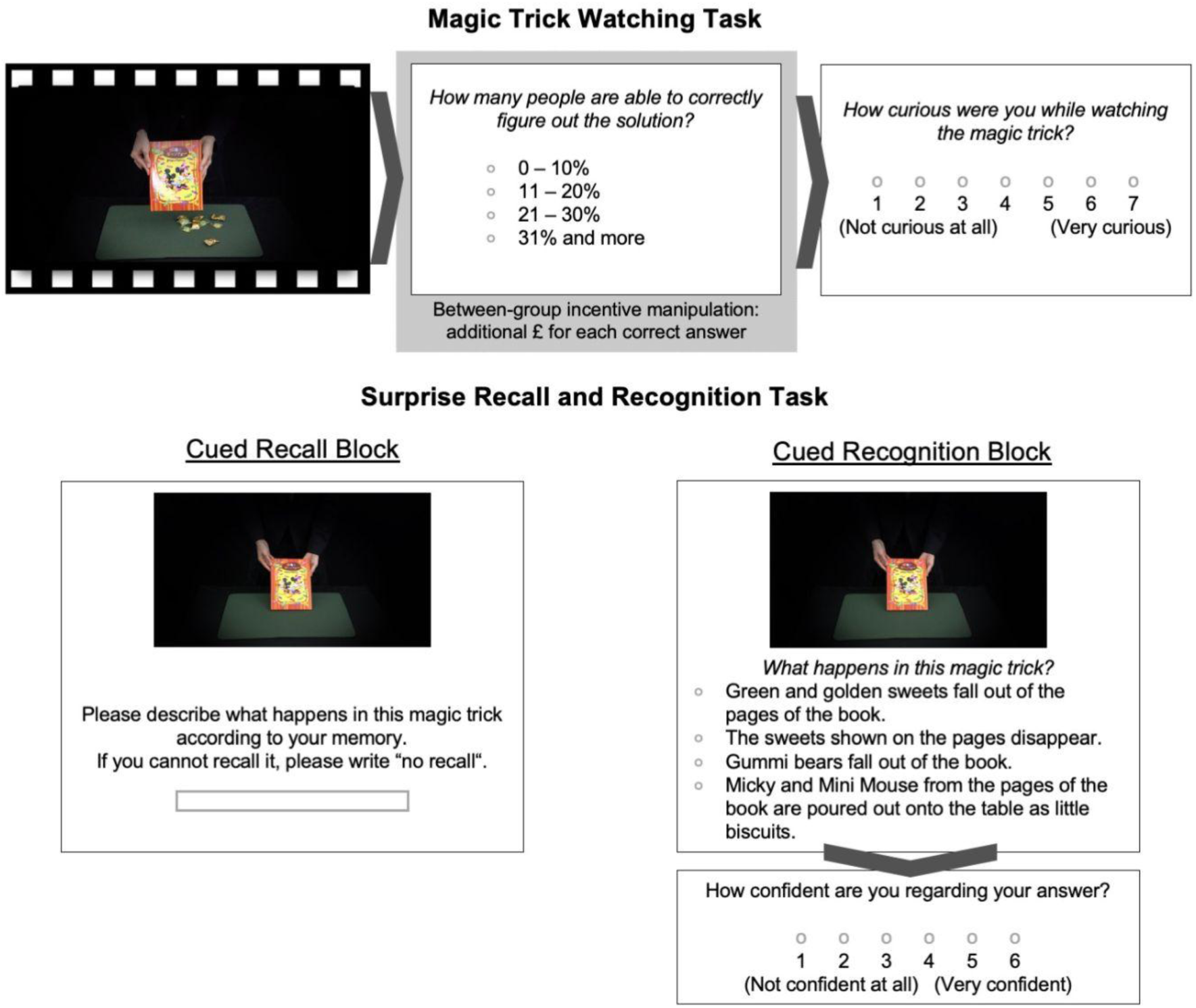
Overview Of The Task Trials. *Note*. The figure illustrates the incidental incentives-motivated learning task as well as the surprise memory test. Task flow is indicated using dark grey arrows. The upper half of the picture shows the magic trick watching task trial as used in online studies. After a magic trick was displayed, participants were asked to give an estimate of how many people (out of 100) could find the solution to the magic trick. In a between-subject design, participants were instructed that they could earn additional monetary rewards for each correct estimate or did not receive such an instruction. Participants were further asked to rate their curiosity regarding the magic trick. The same task was used in the fMRI experiment, but stimuli were edited and jittered fixations in between the magic trick video and ratings were introduced. For more details, please refer to the task description below or see Meliss et al. (2022). The lower half shows the memory task consisting of a cued recall and cued recognition block. Cue images were taken from the magic tricks and the same images were used during both blocks.

In total, the magic trick watching task consisted of 36 trials randomised across three blocks (12 trials each). There were no time-fixed response windows. Participants were able to take breaks in between blocks (self-paced).

##### Surprise Recall And Recognition Task

Approximately one week later, participants’ memory for the magic tricks was tested using a surprise cued recall and a four-alternative forced-choice recognition block (see Figure 1, lower half). During each trial in the cued recall block, the cue image was presented, and participants were asked to describe what has happened in the cued magic trick according to their memory using a free answer format text input. They were instructed to be as descriptive and detailed as possible because their answers would be used to categorise whether they remembered a magic trick. Additionally, they were asked to write ‘no recall’ if they were unable to recall what happened.

During the cued recognition task trials, the same cue image was presented, but this time paired with four choices to answer the question of what happened in this magic trick. The answer options were presented in random order. Behavioural piloting was conducted to achieve wordings of distractor items that do not lead to floor or ceiling effects. After participants selected an answer (self-paced), they were asked to rate their confidence on a scale from 1 (‘not confident at all’) to 6 (‘very confident’). All 36 magic tricks were cued in the recall and recognition task in independent, random order. A break was offered in between both blocks.

##### Task Motivation Inventory (TMI)

To measure task-dependent motivational constructs after the magic trick watching task, the Task Motivation Inventory (TMI) was used. More specifically, the subscales intrinsic motivation (3 items; Elliot & Harackiewicz, 1996), task engagement (3 items; Elliot & Harackiewicz, 1996), interest (3 items; Wigfield & Eccles, 2000), boredom (3 items; Pekrun et al., 2002), effort (5 items; Ryan, 1982), and pressure (5 items; Ryan, 1982)^2^ were used. Participants answered on a 7-point Likert scale from 1 (‘definitely disagree’) to 7 (‘definitely agree’). The item order was randomised, but the same order was used across all participants.

#### 2.1.4 Experimental Procedure

Participants were informed prior to starting the first part that they will be invited to a second part. They were asked to only proceed with the first part if they could participate in the second part one week later. After providing informed consent, participants filled in a quick demographics questionnaire. Afterwards, participants read through the task instructions containing the between-subject incentive manipulation. Half of the participants (incentives condition) were instructed that they could earn an additional £0.10 for each time they estimated correctly how many people would be able to figure out the solution to the magic trick (pseudo-task). The other half of the participants, however, did not receive such an instruction (control condition). Participants were additionally informed that another study was run simultaneously on Prolific, indicating that there was a correct estimate, but that the data collection was still running so there was no feedback. Afterwards, participants completed two practice trials followed by 36 trials of the magic trick watching task distributed across three blocks. At the end, participants completed the TMI. A week later, participants were invited to participate in the second part of the study consisting of the surprise recall and recognition task. Both experiments were executed using a developmental version of Collector (Haffey et al., 2020)

### 2.2 Study 2: Replication Behavioural Study

#### 2.2.1 Participants & Design

To ensure the robustness of effects, we ran a replication of the initial behavioural study with small adjustments. The study was again conducted using Prolific aiming for the predetermined sample size of 40 participants per group applying the same inclusion criteria. As the initial behavioural study, the replication study was set up as a two-part study spaced one week apart. The incentive manipulation was operationalised using a between-subject design setting up two different studies on Prolific. The wording of the incentive manipulation was adopted so that it could be translated to other study settings. More specifically, participants in the incentives condition were instructed that it is possible to earn an additional 50% bonus payment on top of the payment for both tasks if they estimated correctly how many people would be able to figure out the solution on all 36 trials and that this translates into additional £0.10 per correct estimate. Participants were reimbursed £7.50 for their time and received a bonus payment of £0.90 upon completing both parts mirroring chance level performance in the pseudo-task.

Complete data from the first session was received from 40 participants in each group. Because 2 participants in the incentive group did not complete the second session, the final sample size included in the analysis was N = 78 participants (n_control_ = 40, n_incentive_ = 38). The sample description can be found in Table 1. The study was conducted as part of the same ethics approval mentioned above (2016-109-KM).

#### 2.2.2 Material

The same magic trick movie stimuli and cue images were used as described above.

#### 2.2.3 Tasks & Measurements

The same tasks as described above were used. Small adjustments were made in the wordings in the recognition task items to enhance readability (e.g., by adding articles). Additionally, the TMI included all five items for the pressure scale.

#### 2.2.4 Experimental Procedure

Procedures were not modified in between data collections other than the above-mentioned change in the wording of the incentive manipulation. Data was collected using a later developmental version of Collector.

### 2.3 Study 3: fMRI Study

In addition to behavioural effects, we were also interested in the neural mechanisms underlying curiosity-motivated learning of dynamic stimuli, so we adapted the magic trick watching task for use in the fMRI scanner and also added a 10 min rest pre- and post-learning (data not included here). The whole MRI dataset has been made publicly available as the Magic, Memory, and Curiosity (MMC) Dataset (https://doi.org/10.18112/openneuro.ds004182.v1.0.0) and the task data were analysed for this report. We here briefly summarise the methods, while a more detailed description can be found elsewhere (Meliss et al., 2022).

#### 2.3.1 Participants & Design

Participants (see Table 1 for demographic information) were recruited using leaflets that were distributed around the campus to achieve a final sample size of N = 50 (i.e., 25 participants per group). Participants were required to be right-handed. The *a priori* sample size considerations were based on sample sizes used in previous behavioural studies (Murayama & Kuhbandner, 2011) as well as on sample size recommendations for between-subject effects in naturalistic imaging (Pajula & Tohka, 2016; Yeshurun et al., 2017). Similar to the behavioural studies, the fMRI study consisted of multiple sessions: a pre-scanning online assessment, the fMRI lab experiment where the magic trick task was performed inside the MRI scanner and the surprise memory session performed online a week later. In total, participants were reimbursed £30 for their time plus £7.20 additional bonus payment (i.e., chance level performance in the judgement task, see below).

The fMRI also included a between-subject incentive manipulation and participants were assigned to the experimental conditions in an interleaved manner. Using the same wording framework as in the behavioural replication study, participants in the incentive group were instructed that they could receive an additional 50% on top of their payment for the whole data collection if they estimated all 36 trials correctly and that this translates into additional £0.80 per correct estimate^3^. The study protocol was approved by the University of Reading Research Ethics Committee (UREC; 18/18).

#### 2.3.2 Material

In the fMRI study, the same magic tricks were presented as before, but the video files themselves underwent further editing to optimise them for usage within the MRI scanner. Luminance, for instance, was adapted where necessary. Furthermore, a mock video was created and added individually to the beginning of each magic trick. Over a period of 6s, the first frame of each magic trick was displayed overlayed with a black video including a viewing focus that gradually opened up to match the viewing focus of the magic trick file. The resulting magic trick files were on average 38.5s long (SD = 8.63, min = 26.6s, max = 58.64). The same frames as described above were used to create cue images.

#### 2.3.3 Tasks & Measurements

Overall, the tasks were not substantially changed and only adapted for the fMRI environment. The study protocol included more tasks (see Meliss et al., 2022), however, here only the tasks used for the analyses are described.

##### Magic Trick Watching Task

Participants were asked to perform the magic trick watching task inside the MRI scanner (see Figure S1 illustrating the trial structure used in the fMRI experiment). The experiment was displayed on a black background and all text was presented in white unless indicated differently. The beginning of the display of each magic trick video was synced with the scanner TTL (transistor-transistor logic) pulse at the beginning of each repetition time (TR). A jittered fixation (4-10s, TTL aligned, only even integers) was displayed in between the end of the magic trick and the estimate rating. Different from the behavioural studies, the percentage sign was omitted in the answer options and the answer options were displayed in colours matching the button colours on the four-button MRI-compatible response device (https://www.curdes.com/mainforp/responsedevices/buttonboxes/hhsc-1x4-cr.html). Estimate ratings were recorded by pressing the button in the colour of the corresponding estimate. There was a fixed response window of 6 s. If participants chose an estimate sooner, the answer options would turn white. After a brief fixation (0.05 s), the curiosity rating was displayed and a random number was highlighted in red. Participants were instructed to move the highlighted number to the left or right (using index and middle finger, respectively) before confirming their selection using the red button. The fixed response window was 5.95s.

Participants completed two practice trials outside the MRI scanner. Inside the MRI scanner, participants completed 36 trials of the magic trick watching task distributed over three blocks. The order in which magic tricks were displayed was pseudo-randomised to control for trial order effects. Trial orders were simulated so that high and low curiosity magic tricks were equally distributed across blocks (low and high curiosity magic tricks were defined based on data by Ozono and colleagues (2021)) while no more than four magic tricks of each category could follow consecutively. Furthermore, trial orders were restricted so that the maximum range of Spearman-rank correlations between any two trial orders did not exceed a threshold of 0.7. In total, 25 trial orders were simulated and used once in each group.

Self-paced breaks were offered in between blocks. Participants were exposed to the incentive manipulation in written form before the start of the first task block and had to confirm it by pressing a button on the button box. The incentive manipulation was also repeated verbally by the experimenters. Before the start of the second and third block, the incentive manipulation was repeated.

##### Surprise Recall And Recognition Task

No changes were made with respect to the memory task.

##### Task Motivation Inventory (TMI)

The TMI was completed inside the MRI scanner at the end of the experiment. Items were displayed in random order and participants’ responses were collected akin to the curiosity ratings.

#### 2.3.4 Experimental Procedure

After screening procedures and pre-scanning assessments (described elsewhere, Meliss et al., 2022), participants were invited to an fMRI scanning session at the Centre for Integrative Neuroscience and Neurodynamics (CINN) at the University of Reading for a two-hour session. Practice and experiment were presented using PsychophysicsToolbox (PTB) 3 (Brainard, 1997) with GStreamer media framework run on Matlab on a 13-inch Apple MacBook. Practice trials were completed outside the MRI scanner looking directly at the screen, whereas back projection was used during the experiment. Before and after the magic trick watching task, resting-state data (10 min, eyes open) was acquired. At the end of the experiment, the TMI was presented during which the anatomical sequence was run. The follow-up memory test was conducted online: One week later, participants received the link to the surprise memory assessment executed using Collector.

### 2.4 Data Pre-processing And Analysis

#### 2.4.1 Behavioural Data

Behavioural data from each data collection were processed and analysed in the same way. All behavioural pre-processing and analysis were carried out in R 3.6.3 (R Core Team, 2020).

To test for between-group differences in motivation (TMI scores as well as ratings of curiosity obtained in the magic trick watching task), data from the TMI were analysed using Welch’s Two-Sample t-tests. Curiosity ratings for the magic trick movies were analysed using Linear Mixed Effects (LME) models with the *lme4* package (Bates et al., 2015) specifying a fixed effect for incentives (effect-coded; -1 = control group, 1 = incentive group) and random effects for intercepts of participants and stimuli.

Data from the recognition block was dummy-coded by comparing the chosen response to the correct answer. Additionally, recognition performance was combined with confidence ratings. Specifically, a correct answer chosen with a confidence larger than three was coded as correct for ‘high confidence recognition’, a recollection-based recognition memory measurement (Yonelinas, 2001, 2002). For the recall performance of the answers collected in the cued recall paradigm, a script was used to assign 0 to all answers matching ‘no recall’ (or variants thereof). All remaining answers were coded by the same rater across all three data collections. A trick was rated as recalled if the change that occurred was remembered.

Our main analyses focused on the effects of curiosity, monetary incentives, and their interaction on memory encoding. Encoding data were analysed using a meta-analytic approach. For each data collection, Generalised LME (gLME) models were applied specifying fixed effects for curiosity, incentives, and their interaction as well as random effects for the participant and stimulus intercept and random slopes for the curiosity effect. Curiosity ratings were mean-centred within each participant and incentive manipulation was again effect-coded. The same model was run on three different memory thresholds: correct recognition (regardless of confidence), high confidence recognition, and cued recall. To further investigate whether incentives and curiosity influence the quality of memory in an exploratory analysis, we systematically varied the confidence cut-off, creating additional dependent variables (recognition with confidence > 0 through to recognition with confidence > 5) and applied the same gLME model as described above. The unstandardised parameter estimates from the gLME models (i.e., beta estimates and standard errors) from each data collection were extracted and submitted to a fixed-effect meta-analysis (weighted least squares) using the *metafor* package (Viechtbauer, 2010) to integrate individual coefficients from the three data collections.

#### 2.4.2 fMRI Data

##### fMRI Acquisition And Pre-processing

fMRI data were obtained in a 3.0 T Siemens Magnetom Prisma scanner with a 32-channel head coil. Whole brain images were acquired (37 axial slices, 3 x 3 x 3 mm, interslice gap of 0.75 mm) using an echo-planar T2*-weighted sequence (TR = 2000 ms, echo time = 30 ms, field of view: 1,344 x 1,344 mm^2^, flip angle: 90°).

Pre-processing steps included B_0_ distortion correction, despiking, slice-timing and head motion correction, and normalisation to MNI space using the ICBM 2009c Nonlinear Asymmetric Template. Additionally, data were smoothed to achieve an approximate smoothness of full width half maximum kernel of 8mm and time series were scaled to a mean of 100. Local white matter time series, the first three principal components of the lateral ventricles, as well as motion estimates, were included as regressors of no interest to denoise the data. During linear regression, time courses were also band-pass filtered for frequencies between 0.01 and 0.1 Hz. Time points were censored (i.e., set to zero) if the Euclidean norm of per-slice motion exceeded 0.3 mm or if more than 10% of brain voxels were outliers.

##### Intersubject Correlation (ISC) Analysis

Due to increased stimulus complexity in naturalistic paradigms, the applicability of traditional analysis methods developed for task-based fMRI relying on specifying onset and duration of stimuli (e.g., general linear models; GLMs) is limited and model-free approaches are used frequently (Sonkusare et al., 2019). One of these data-driven methods is intersubject correlation (ISC; Hasson et al., 2004). Here, the assumption is that the brain response when perceiving and processing naturalistic stimuli is composed of a stimulus-driven signal as well as spontaneous activity unrelated to the stimulus (Nummenmaa et al., 2018). The stimulus-driven signal is time-locked to the stimuli and shared across subjects whereas the intrinsic fluctuations are cancelled out as noise. To determine brain areas that encode information about the presented stimuli consistently across subjects, the time course of a given voxel in subject A is correlated with the time course of the same voxel in subject B. This is repeated for each voxel in the brain for each pair of participants in the sample creating pairwise ISC maps.

During the magic trick watching task, the beginning of each magic trick video was aligned with the beginning of a TR. Likewise, the jittered fixation after the magic trick presentation was aligned with the beginning of a TR and presentation times and response windows were multiple of the TR. These steps were undertaken to allow that the time series could be concatenated (see Figure 2A) to (a) remove volumes of no interest, (b) reorder the volumes so that the concatenated order would remain invariant across subjects irrespective of the pseudo-randomised order in which the magic tricks were presented (see Thomas et al., 2018), and (c) account for the delay in the hemodynamic response function (HRF) by shifting the time course. Volumes acquired during the mock video presentation, fixation and estimate/curiosity ratings were considered as volumes of no interest because ISC critically relies on subjects receiving the same time-locked stimuli and transient, non-specific activity can be found at stimulus onset (Nastase et al., 2019).

**Figure 2.**
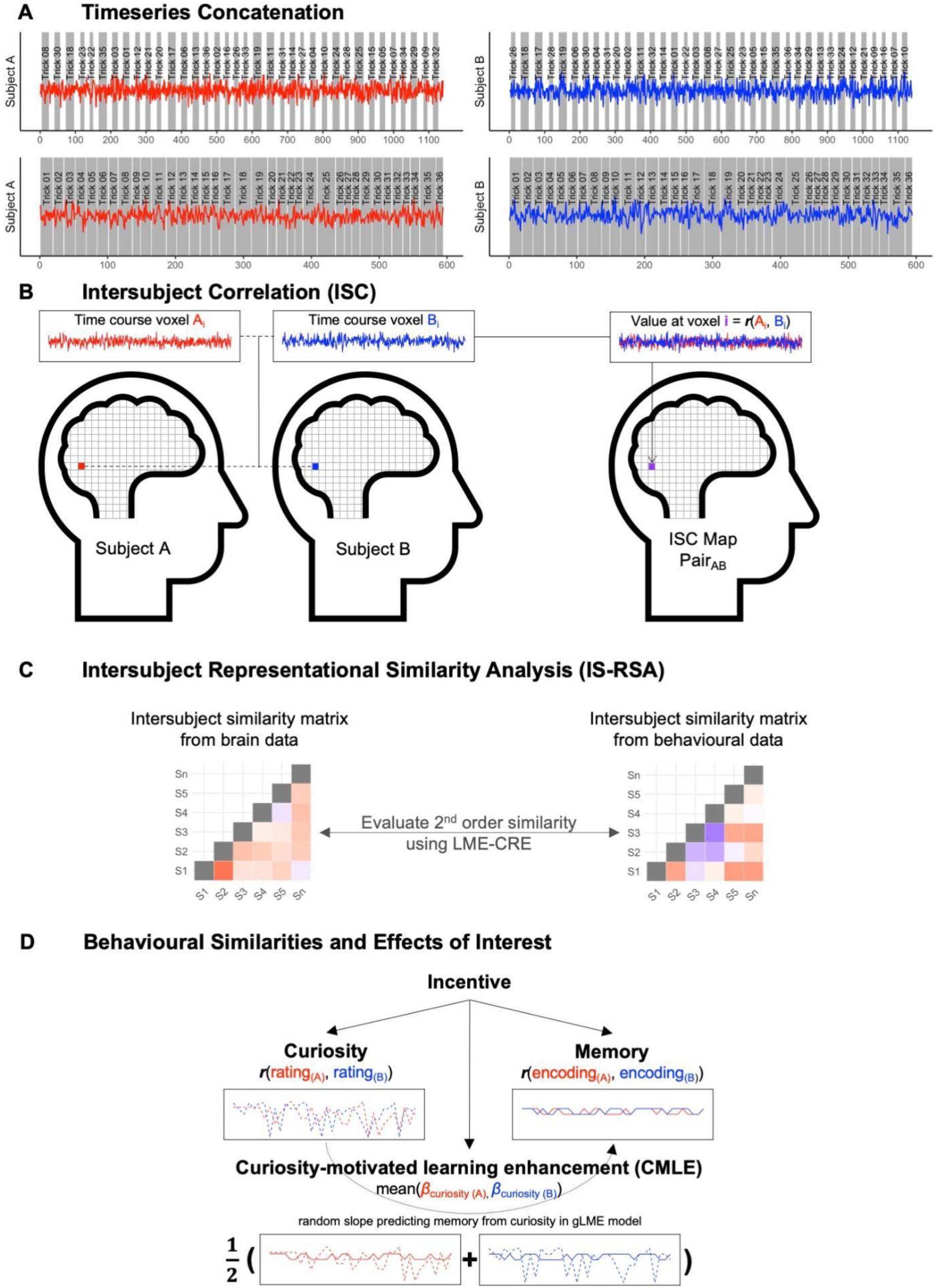
Illustration Of Processing And Analysis Methodology Within The Intersubject Framework. *Note.* To account for the dynamic nature of the stimuli, intersubject correlation (ISC) analysis was applied. (A) In the first step, data were concatenated to remove volumes of no interest, reorder volumes, and account for the lag in the HRF. (B) The concatenated time series of each voxel were correlated for each pair of participants creating pairwise ISC maps representing similarity in the brain response between participants (figure adapted from Nastase et al., 2019). (C) To anchor idiosyncratic response patterns to behavioural measurements, intersubject representational similarity analysis (IS-RSA) was used to relate similarities in the brain response to similarities in behavioural measurements (figure adapted from Finn et al., 2020). (D) Behavioural measures of interest were curiosity, encoding and curiosity-motivated learning enhancement (CMLE). To determine behavioural similarities in curiosity and encoding, the time course of rating and encoding were correlated for each pair of participants. For CMLE, each subject’s random slope predicting memory encoding with curiosity estimated by the behavioural gLME was extracted and the mean as a non-parametric difference measure was calculated for each pair.

As assumptions regarding the duration of the HRF lag to account for in ISC analyses vary (Hasson et al., 2004; Nummenmaa et al., 2012; Zadbood et al., 2017), a preliminary intersubject pattern correlation (ISPC; J. Chen et al., 2017) - a spatial form of ISC - was computed to determine the optimal HRF lag. This preliminary analysis indicated the optimal HRF lag to be 4 TRs (see supplementary material and Figure S2). The concatenated time series consisting of 594 volumes were correlated for each pair of participants (using AFNI’s *‘3dTcorrelate’*, Figure 2B) resulting in 1225 pairwise ISC maps that were Fisher’s *z*-transformed.

To determine brain areas showing significant synchronicity between subjects, linear mixed effect models with crossed random effects (LME-CRE; G. Chen et al., 2017) were specified to predict the pairwise Fisher’s *z*-transformed ISC maps (using AFNI’s *‘3dISC’*). The LME-CRE framework does not only account for the interrelatedness in the pairwise ISC map data by specifying crossed random intercepts for both subjects in each pair but also offers analytical flexibility to specify grouping variables to investigate the effects of incentives on ISC during magic trick watching as well as of other covariates (see below). To specify the fixed effect of incentives, deviation coding was adopted where 0.5 was assigned to subjects in the control group and -0.5 was assigned to subjects in the incentive group. By adding up these values for each pair, group was defined as 1 (both subjects in control group), 0 (both subjects in different groups), or -1 (both subjects in the incentive group).

##### Intersubject Representational Similarity Analysis (IS-RSA)

Nastase and colleagues (2019) proposed a formal definition of ISC where they divided the stimulus-driven component further into processes that are consistent across subjects and idiosyncratic responses that are nonetheless induced by the stimulus but characterised by timings and intensities specific to each subject. The consistent response can be estimated by averaging the ISC given that subject-specific and spontaneous responses will average out. To quantify the subject-specific responses in the time courses, other known information about the subjects can be used to ‘anchor’ the response - an approach known as intersubject representational similarity analysis (IS-RSA; Finn et al., 2020; Nummenmaa et al., 2012). More specifically, the similarity in participants’ behavioural data (e.g., trait scores, Finn et al., 2018; age, Moraczewski et al., 2018; recall performance, Nguyen et al., 2019; behavioural ratings, Nummenmaa et al., 2012) can be used to predict the similarity in the brain response (Figure 2C) by, firstly, calculating subject-by-subject similarity matrices separately for behavioural and brain data. In a second step, the geometry of both matrices can be compared or matched correlationally based on the second-order isomorphism within representational similarity analysis (RSA; Kriegeskorte et al., 2008). The second-order similarity can be evaluated using LME-CRE. Importantly, the pseudo-randomisation of trials allows that similarities in brain responses between participants can be attributed to the behavioural anchor rather than to similarities in the trial order.

Here, we were interested in how similarity in (1) curiosity, (2) memory encoding, and (c) curiosity-motivated learning enhancement (CMLE) predicts similarity in the neural responses across subjects (Figure 2D). To calculate the subject-by-subject similarity matrices in the first two instances, the trial-by-trial values (subject-wise mean-centred curiosity ratings and dummy-coded encoding performance on the high confidence criteria, respectively) were correlated for each pair of participants (after re-ordering the values for each subject to account for the pseudo-randomisation). To control for potentially shared variance between the similarity matrix of curiosity and the similarity matrix of memory, Fisher’s *z*-transformed pairwise curiosity correlations were residualised by removing the proportion of variance that can be linearly predicted by Fisher’s *z*-transformed pairwise memory correlations. Likewise, Fisher’s *z*-transformed pairwise memory correlations were residualised by removing the proportion of variance that can be linearly predicted by Fisher’s *z*-transformed pairwise curiosity correlations. In doing so, the unique effects of curiosity and memory could be investigated.

CMLE was quantified by extracting the individual curiosity beta values (estimated by the specification of random slopes predicting memory with curiosity) from the gLME model for high confidence recognition and mean-centring them^4^. The beta value quantifies the magnitude of the association between curiosity and memory for each individual. Because there was only one value per subject (rather than a time course), the similarity matrix was calculated using the Anna Karenina (AnnaK) model providing a metric reflecting the absolute position on the scale, i.e., the mean of both subjects (Finn et al., 2020). This is preferable compared to using a relative distance metric like the Euclidean distance as it allows for a scenario where low scoring subjects are more similar to other low scoring ones, but high scoring subjects are less similar to each other. Previous studies using working memory in IS-RSA found that the AnnaK model fitted the data better than the Euclidean distance and yielded to higher replicability between samples (Finn et al., 2020). Another benefit of using the mean is that effects in both directions can be captured: if the correlation between brain and behaviour is positive, then high scorers are alike and low scorers different whereas a negative sign indicates that low scorers are alike and high scorers different.

To link idiosyncratic responses in the stimulus-driven brain response to the behavioural effects of interest, LME-CRE were used to predict the pairwise Fisher’s *z*-transformed ISC maps. As described above, separate models were estimated for unique curiosity, unique memory, and CMLE, again specifying crossed random intercepts for both subjects in each pair. Fixed effects were specified for group (incentive vs. no incentive, effect-coded), the respective behavioural similarity (of curiosity, memory, or CMLE) as well as their interaction. Behavioural similarity was grand-mean centred before computing the interaction term with the group variable.

##### Thresholding And Regions-Of-Interest (ROI) Approach

All analyses were conducted at the whole brain level specifying a grey matter (GM) mask: during pre-processing, each subject’s grey matter (GM) mask based on FreeSurfer parcellation was transformed to echo-planar imaging (EPI) resolution. After averaging the masks across the sample, the mean image was thresholded so that a voxel was included in the group GM mask if it was GM in at least 50% of the sample.

To account for the multiple testing problem due to mass-univariate testing, cluster-extent based thresholding was performed using the recommended initial threshold of *p* value = 0.001 (Woo et al., 2014). The resulting map was cluster-extent corrected based on the output of simulations performed using ‘3dClustSim’ for first nearest neighbours clustering (NN = 1; faces of voxels touch) and a cluster threshold of *α* = 0.05 resulting in a threshold of *k* = 20 voxels. Unthresholded statistical maps were uploaded to NeuroVault (https://neurovault.org/collections/12980/).

In addition to whole brain analysis, we were also interested in regions previously implicated in motivated learning and *a prior*i defined the following regions-of-interest (ROIs): aHPC, NAcc, CN, and VTA/SN. The aHPC has been chosen as increased activity for remembered compared to forgotten items is predominantly centred in anterior parts of the HPC (Kim, 2011; Spaniol et al., 2009). The aHPC is also sensitive to the effects of incentives and motivationally relevant information on encoding (Adcock et al., 2006; Poppenk et al., 2013). To create the aHPC ROI, AFNI’s ‘whereami’ was used to extract the bilateral HPC from the Glasser Human Connectome Project atlas (Glasser et al., 2016). Following the recommendations by Poppenk et al. (2013), the aHPC was created by using the MNI coordinate y = 21P to determine the uncal apex as a landmark to divide anterior and posterior HPC (‘3dZeropad’). To create ROI masks for NAcc, CN, and VTA/SN; atlaskit (https://github.com/jmtyszka/atlaskit) was used to extract the NAcc, CN, Substantia Nigra pars reticulata (SNr), Substantia Nigra pars compacta (SNc), and Ventral Tegmental Area (VTA) from a high-resolution probabilistic subcortical nuclei atlas in MNI space (Pauli et al., 2018) specifying a probability threshold of 15%. This is similar to procedures by others presenting magic tricks inside the MRI scanner (Lau et al., 2020). To create the VTA/SN mask, the masks for VTA, SNr, and SNc were combined. In total, the aHPC mask contained 162 voxels, the CN mask contained 573 voxels, and the NAcc and VTA/SN mask both contained 60 voxels each (see Figure S3). To correct for multiple comparisons within each ROI, False Discovery Rate (FDR) correction was applied at *q* = 0.05. Additionally, clusters were thresholded at *k* = 5 (NN = 1). ROI masks can be accessed in the NeuroVault collection (https://neurovault.org/collections/12980/).

## 3 Results

### 3.1 Behavioural Data

The groups did not differ in their motivation in any TMI scale in any of the assessments (all *p* > 0.09). Likewise, no group difference was observed in the curiosity ratings (all *p* > 0.199). The detailed results for TMI scores and curiosity ratings can be found in Table S1 and S2 in the supplementary material, respectively.

Next, we investigated the effects of curiosity, incentives, and their interaction on memory encoding specifying the same gLME model for each data collection and memory measurement (recognition, high confidence recognition, cued recall) to submit the estimates into fixed effects meta-analyses for each memory measurement separately. The results of the fixed effects meta-analyses are shown in Table 2. Curiosity had a positive effect on memory encoding: magic tricks for which participants reported higher curiosity were more likely to be encoded. While the overall curiosity effect was not significant for recognition per se, significant effects were observed for high confidence recognition and cued recall. With respect to the effect of monetary incentives on memory encoding, the effects were overall positive, i.e., participants in the incentive group were more likely to encode the magic tricks compared to participants in the control group. However, the overall effect only reached significance for the high confidence recognition memory measurement. The interaction between monetary incentives and curiosity did not reach significance for any of the memory thresholds investigated.

**Table 2.** Integrated Results Of gLME Models Predicting Memory Encoding Using Curiosity, Monetary Incentive, And Their Interaction. *Note*. Separate models were run for each memory threshold. gLME = Generalised Linear Mixed Effects. SE = standard error. OR = Odds Ratio, CI = confidence interval.

|  | *b* (*SE*) | OR [95%-CI] | *z* value | *p* value |
| --- | --- | --- | --- | --- |
| Curiosity |  |  |  |  |
| Recognition | 0.023 (0.023) | 1.02 [0.98; 1.07] | 0.988 | 0.323 |
| High confidence recognition | 0.084 (0.022) | 1.09 [1.04; 1.14] | 3.766 | < 0.001 |
| Cued recall | 0.098 (0.025) | 1.10 [1.05; 1.16] | 3.842 | < 0.001 |
| Monetary incentive |  |  |  |  |
| Recognition | 0.084 (0.050) | 1.09 [0.99; 1.20] | 1.676 | 0.094 |
| High confidence recognition | 0.155 (0.067) | 1.17 [1.03; 1.33] | 2.336 | 0.019 |
| Cued recall | 0.119 (0.075) | 1.13 [0.97; 1.30] | 1.599 | 0.110 |
| Interaction |  |  |  |  |
| Recognition | -0.002 (0.022) | 1.00 [0.96; 1.04] | -0.070 | 0.944 |
| High confidence recognition | -0.010 (0.021) | 0.99 [0.95; 1.03] | -0.479 | 0.632 |
| Cued recall | -0.025 (0.024) | 0.98 [0.93; 1.02] | -1.058 | 0.290 |

We then further examined the quality of recognition memory by changing the confidence cut-off threshold gradually (0 ≤ cut-off ≤ 5). Again, the same gLME model was run for each confidence threshold and each data collection and estimates were integrated using a fixed effects meta-analysis (for detailed results for each effect on each threshold, see Table S3) to extract the integrated b estimates for each effect at each confidence cut-off. Then, to examine how the cut-off is related to memory enhancement effect, the integrated fixed effects b estimates were predicted using the confidence cut-off in a linear model separately for each effect. The cut-off was scaled from 0 to 5 so that the intercept is interpretable.

The results of the exploratory analysis are illustrated in Figure 3 and the detailed regression table can be found in Table S6. More specifically, they show that when calculating a linear regression to predict the integrated curiosity effect b values based on the confidence cut-off, the confidence cut-off was a significant predictor in the model (*B* = 0.021, 95%-CI [0.011; 0.031], *p* = .004) indicating that the integrated curiosity effect increases as the confidence cut-off increases: the Odds Ratio (OR) of the curiosity effect was 1.02 for confidence cut-off = 0 and 1.12 for confidence cut-off = 5.

**Figure 3.**
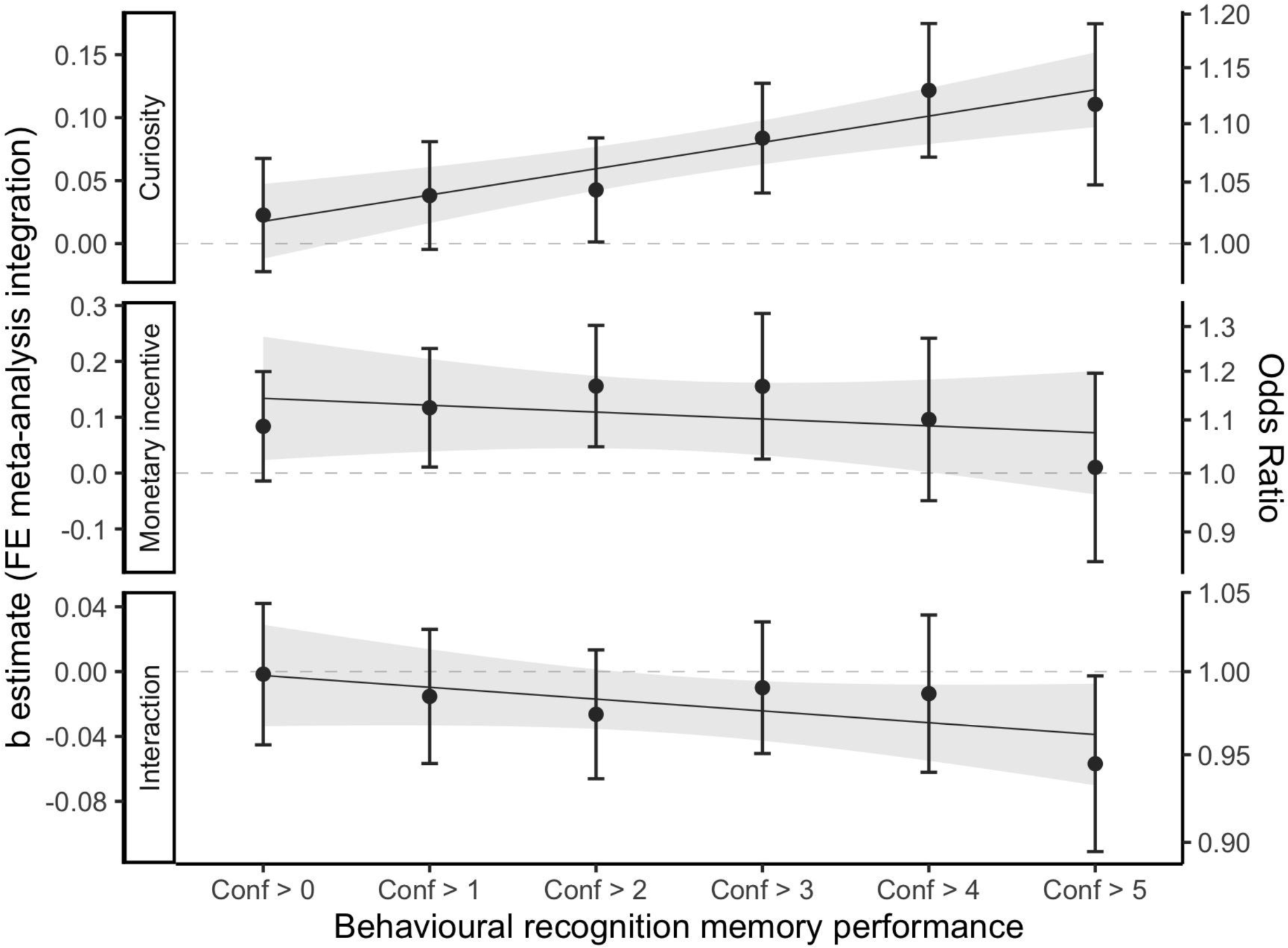
Integrated Fixed Effects Of Curiosity, Monetary Incentive, And Their Interaction As A Function Of Confidence Cut-Off. *Note.* The x-axis shows the gradual confidence cut-off and y-axis illustrates the integrated effect size (left - unstandardised, right - OR). Each panel shows one of the fixed effects specified in the gLME model. The integrated b estimate for each effect and confidence threshold is plotted and error bars indicate 95%-CI. The regression line illustrates the linear model predicting the effect with the gradual confidence cut-off.

However, in the model predicting the integrated monetary incentive effect b values with the confidence cut-off, the cut-off was not a significant predictor in the model (*B* = - 0.012, 95%-CI [-0.049; 0.024], *p* = .402). Likewise, using a linear model to predict the integrated interaction effect b values using the confidence cut-off, confidence cut-off was not a significant predictor (*B* = -0.007, 95%-CI [-0.018; 0.003], *p* = .122).

The results suggest that only the curiosity effect, but not the monetary incentive or the interaction effect, is sensitive to the confidence cut-off. More specifically, they show that the more confidently participants recognise the correct answer option, the larger the effect of curiosity on encoding. Monetary incentive and interaction effect, on the other hand, remain invariant regarding the confidence thresholds.

The results of all 21 individual gLME models (seven memory measurements in three experiments) can be found in Table S4. Additionally, Figure S4 contains the equivalent of Figure 3 plotting the effects from each data collection individually. Because 8 of 21 gLME models produced a singular fit warning during execution, all analyses were repeated using a simplified gLME model with a reduced random effects structure omitting the random slopes for the curiosity effect. Applying this reduced gLME model, however, did not affect the results of the meta-analyses and associated confidence cut-off linear model (see Table S5, S6, and S7 as well as Figure S5 and S6).

### 3.1 fMRI Data

#### 3.1.1 Intersubject Correlation (ISC)

ISC analyses were carried out to identify brain areas with activity driven by magic trick watching. Significant ISC was found bilaterally in all four ROIs (aHPC, VTA/SN, NAcc, and CN; see Figure S7). Shared activity measured as significant ISC in the reward network has previously been observed in naturalistic viewing paradigms when presenting comedy movie clips to participants (Jääskeläinen et al., 2016).

At the whole brain level, widespread cortical and subcortical synchronisation (Figure 4A, Table S8) was observed, especially dominant in the bilateral visual cortex as well as bilateral parietal somatosensory (BA 2, BA 5, BA 40, BA 1/2/3) and in attention-related areas (BA 7 and BA 39) as well as bilateral premotor and supplementary motor areas (BA 6, BA 8). Overall, this is in line with other studies showing that dynamic stimuli synchronise brain activity in visual areas (e.g., Aliko et al., 2020; Baldassano et al., 2017; Hasson et al., 2004; Nguyen et al., 2019), but also with prepositions linking motor and somatosensory areas to the observation of actions (Keysers et al., 2010; Thomas et al., 2018). Likewise, the decline of the ISC from posterior to anterior as well as from lateral to medial areas in the brain can be attributed to higher intersubject variability in the stimulus-induced response in ‘intrinsic systems’ (e.g., prefrontal and cingulate cortices; Ren et al., 2017).

**Figure 4.**
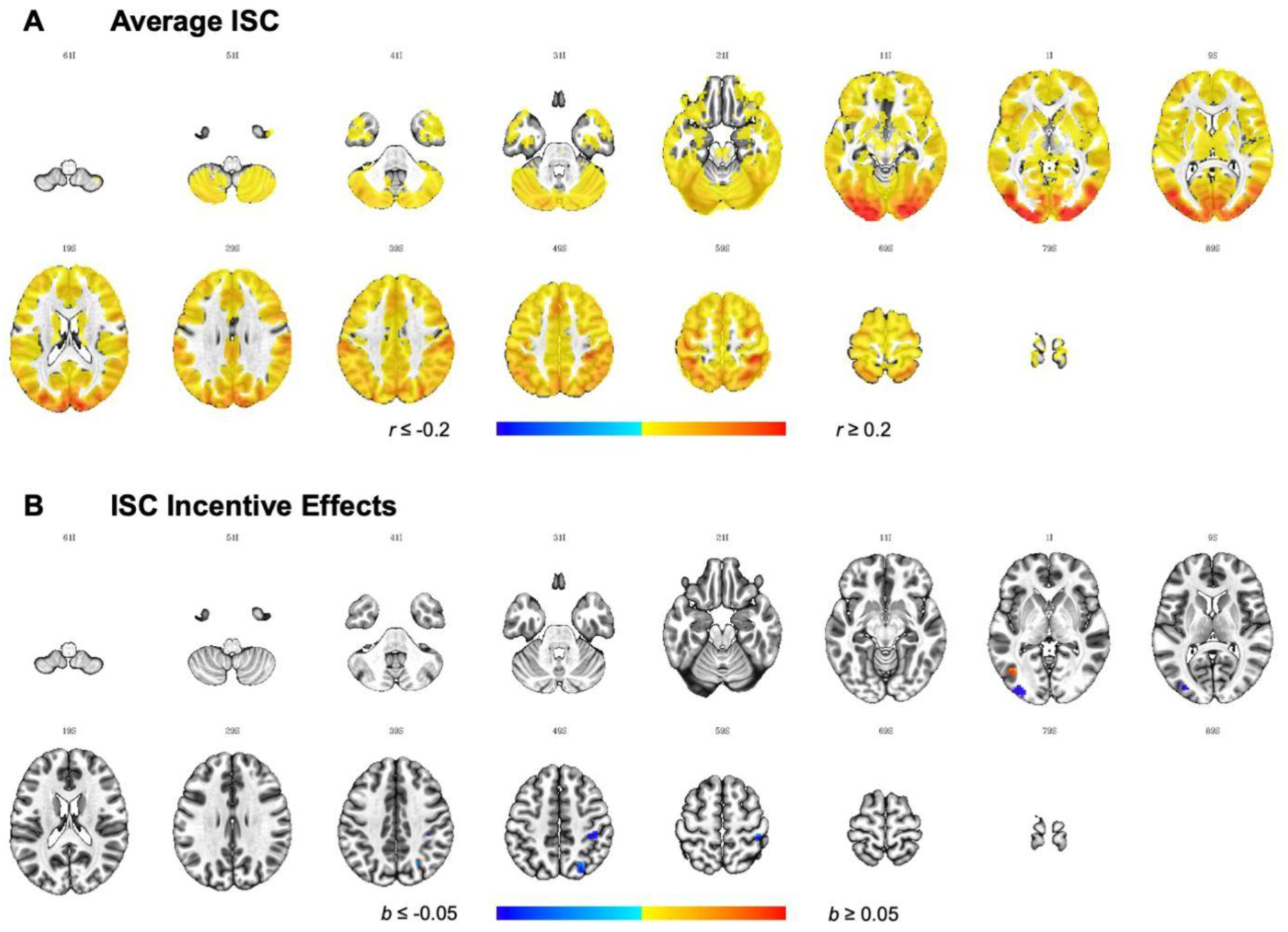
Whole Brain ISC And Incentive Effects Therein. *Note*. Results are thresholded at *p* < 0.001, cluster-extent corrected at *k* = 20 (equivalent to per-cluster α = 0.05) and plotted on the ICBM 2009c Nonlinear Asymmetric Template. Images are displayed in neurological orientation, where the left side of the brain is depicted on the left side of the image. While (A) highlights widespread ISC across cortical and subcortical areas during magic trick watching across both groups, (B) shows clusters where the ISC is higher in the incentive group compared to the control group in blue, and a cluster where ISC is higher in the control group in red.

We also investigated whether the availability of incentives had an effect on the ISC. While no effects were found in the ROIs, four clusters were found at the whole brain level (Figure 4B, Table S8). More specifically, in the incentive group, we found higher ISC in areas in the left middle occipital gyrus, right postcentral gyrus (BA 2), and right intraparietal sulcus (IPS). Higher ISC in the control group was observed in the left lateral occipital cortex (Area V5/MT+).

#### 3.1.2 Intersubject Representational Similarity Analysis (IS-RSA)

IS-RSA were carried out to identify brain regions with intersubject temporal dynamics reflecting the intersubject variability in our behavioural effects of interest as well as brain regions where this association was influenced by the incentive manipulation. For this purpose, for each behavioural effect of interest, an LME-CRE model was specified with fixed effects for group, behavioural similarity, as well as their interaction. The inclusion of the covariate and the interaction effect did not affect the main effect of incentive (all correlations with unthresholded incentive effects reported above ≥ 0.92), hence the incentive effects are not further discussed. Below, the main effects of each behavioural variable are described before discussing the interaction effects and results for the ROI analysis are reported followed by whole brain analysis.

##### IS-RSA For Each Behavioural Effect Of Interest

Here, the main effects of each behavioural variable are reported highlighting clusters where the behavioural similarity matrix was predictive of the neural similarity matrix. The underlying assumption is that participants similar in behavioural effects of interest (e.g., curiosity ratings) will process the magic trick videos more similarly and regions involved in these processes will reflect this similarity correspondingly and hence are detected in this analysis.

##### Curiosity Effect

The curiosity effect was defined as the pairwise correlation of trial-by-trial curiosity ratings. Importantly, we here used unique effects of curiosity where Fisher’s *z*-transformed pairwise curiosity correlations were residualised by removing the proportion that can be linearly predicted by Fisher’s *z*-transformed pairwise memory correlations. No activity in the four ROIs survived thresholding. At the whole brain level, seven positive clusters were found (Figure 5A, Table S9) where idiosyncratic patterns in curiosity were anchored to the brain response. These clusters were located in the left primary visual cortex (V1), right inferior frontal gyrus (pars opercularis), bilateral supplementary motor area (BA 8), left postcentral gyrus (primary somatosensory cortex), left precuneus (BA 7), right anterior insula cortex (AIC) and right supramarginal gyrus (BA 40).

**Figure 5.**
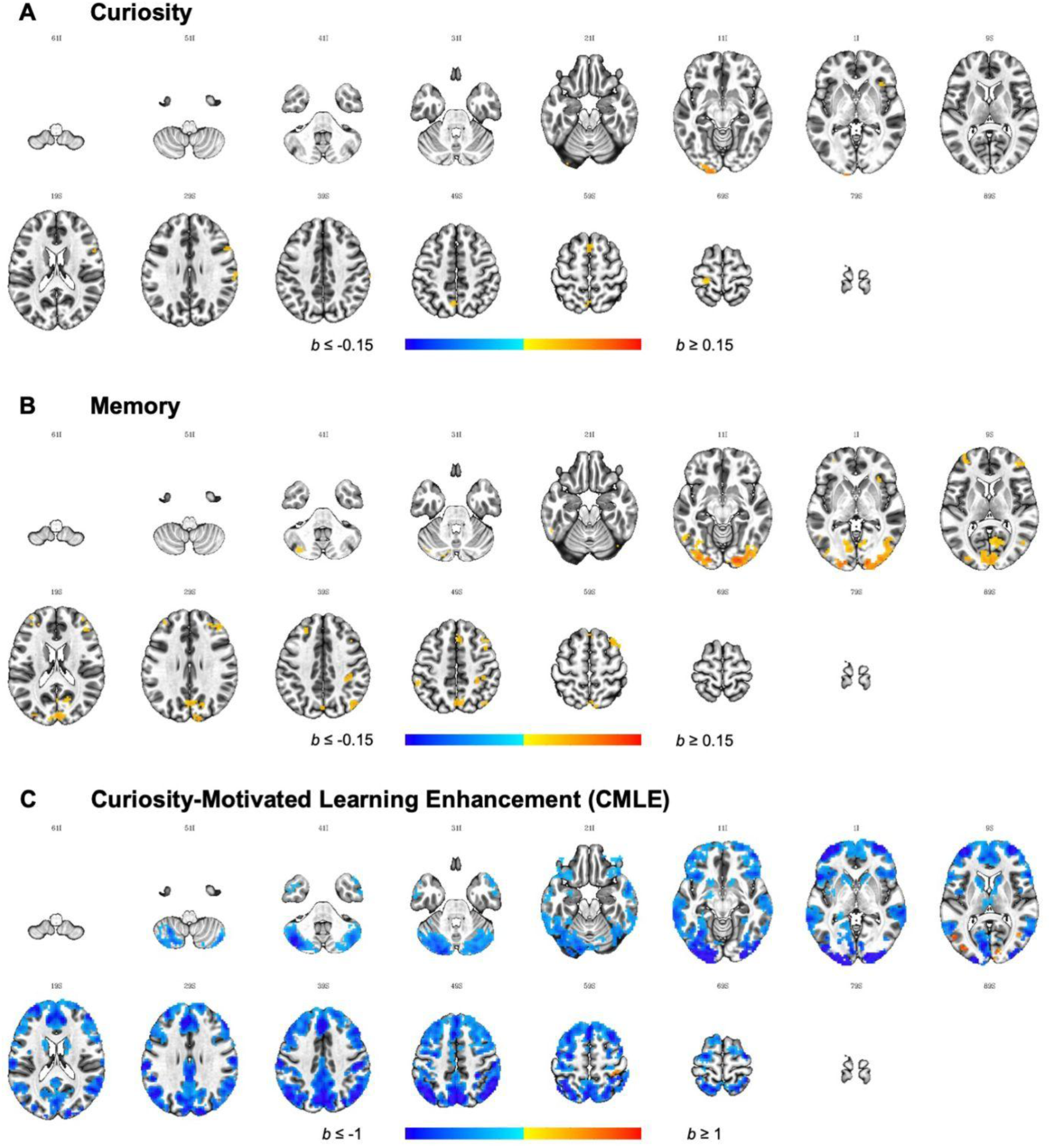
Whole Brain IS-RSA For Each Behavioural Effect Of Interest. *Note*. Results are thresholded at *p* < 0.001, cluster-extent corrected at *k* = 20 (equivalent to per-cluster α = 0.05) and plotted on the ICBM 2009c Nonlinear Asymmetric Template. Images are displayed in neurological orientation, where the left side of the brain is depicted on the left side of the image.

##### Memory Effect

The memory effect was defined as pairwise correlation of trial-by-trial encoding performance ratings. Again, the unique contribution of memory was investigated akin to what was described above in the context of curiosity. Similarity in brain response could be anchored to similarity in memory encoding in a bilateral cluster in the CN ROI (Figure 6, Table S9), however, no effects were observed for the other three ROIs. At the whole brain level, 21 clusters were found (Figure 5B, Table S9). More specifically, similarity in memory encoding positively predicted similarly in brain response bilateral visual areas as well as the left cerebellum, the bilateral superior (BA 46, BA 9-46, medial BA 8) and middle frontal gyrus (BA 6, BA 8), precuneus (BA 7) and lateral parietal areas including the right angular gyrus (BA 39) and somatosensory areas (BA 2, BA 40), the left lateral temporal gyrus (BA 37, fusiform and inferior temporal gyrus), right middle occipital gyrus (Area V5/MT+) and the right AIC.

**Figure 6.**
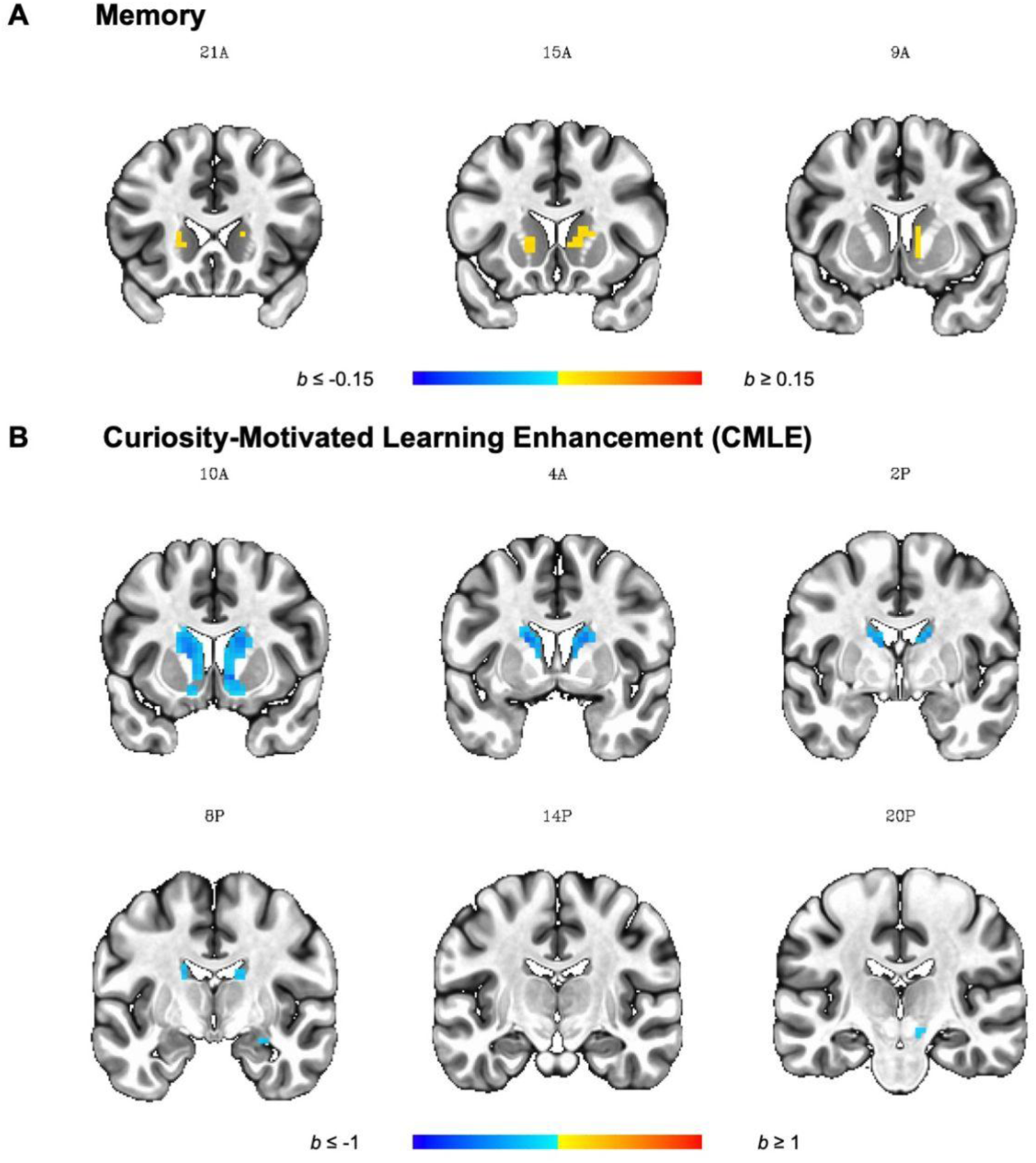
Effects Of Memory And CMLE In The ROIs. *Note.* Results are FDR-corrected at *q* < 0.05, cluster-extent corrected at *k* = 5 and plotted on the ICBM 2009c Nonlinear Asymmetric Template. Images are displayed in neurological orientation, where the left side of the brain is depicted on the left side of the image.

##### Curiosity-Motivated Learning Enhancement (CMLE) Effect

CMLE was defined based on the random slope predicting memory from curiosity in the gLME model and individual values were extracted. Using the AnnaK model to determine the behavioural similarity matrix, the prediction was tested whether participants high in CMLE share similar patterns of brain activity while people low in CMLE show more variability and vice versa (rather than testing for brain areas where similarity is predicted by similarity in CMLE in a linear fashion).

In the ROI analysis, IS-RSA CMLE were found in all 4 ROIs (Figure 6B, Table S9), all of them in a negative direction suggesting that participants with high CMLE scores had less similar brain activity compared to participants with low scores. More specifically, clusters were identified in the right aHPC, right VTA/SN, bilateral CN, and bilateral NAcc. Additionally, 15 clusters survived cluster-extent thresholding at the whole brain level out of which 5 were positively and 10 negatively directed (Figure 5C, Table S9). Positive clusters were located in the bilateral middle temporal gyrus, the left middle occipital gyrus, the right calcarine gyrus, and the right postcentral gyrus. In these positive clusters, subjects high in CMLE are more alike than subjects low in CMLE who are more different in their brain response.

In negative clusters, on the other hand, subjects low in CMLE are more alike and subjects high in CMLE are more different. Negative clusters were spread across large portions of the brain, in subcortical (e.g., striatum and thalamus) as well as cortical areas along the anterior and posterior midline (e.g., ACC, SMA, superior medial gyrus, precuneus, PCC, and cuneus), visual cortex, cerebellum, postcentral gyrus and posterior parietal cortex (PPC), the bilateral middle temporal gyrus, bilateral anterior insula cortex (AIC), as well as dorsolateral prefrontal cortex (dlPFC; centred around the MFG) and anterior PFC stretching into the frontal operculum/anterior insula (fO/aI).

##### IS-RSA For The Interaction Between The Incentive Manipulation And Each Behavioural Effect Of Interest

Due to the inclusion of group as a fixed effect in the LME-CRE model, it was further possible to determine brain areas where the behavioural similarity matrix predicted the neural similarity matrix differently depending on the availability of monetary incentives. In doing so, clusters could be identified where the behavioural effect is only predictive in one group or more strongly predictive in one group.

##### Curiosity Incentive Interaction

When looking at whether the incentive manipulation influences how similarity in curiosity predicts similarity in the brain response in the *a priori* defined ROI, no clusters survived thresholding. At the whole brain level, two clusters in the bilateral occipital cortex survived thresholding (Figure 7A, Table S10). In both clusters, similarity in curiosity was more predictive of similarity in the neural responses during magic trick watching in the control compared to the incentive group.

**Figure 7.**
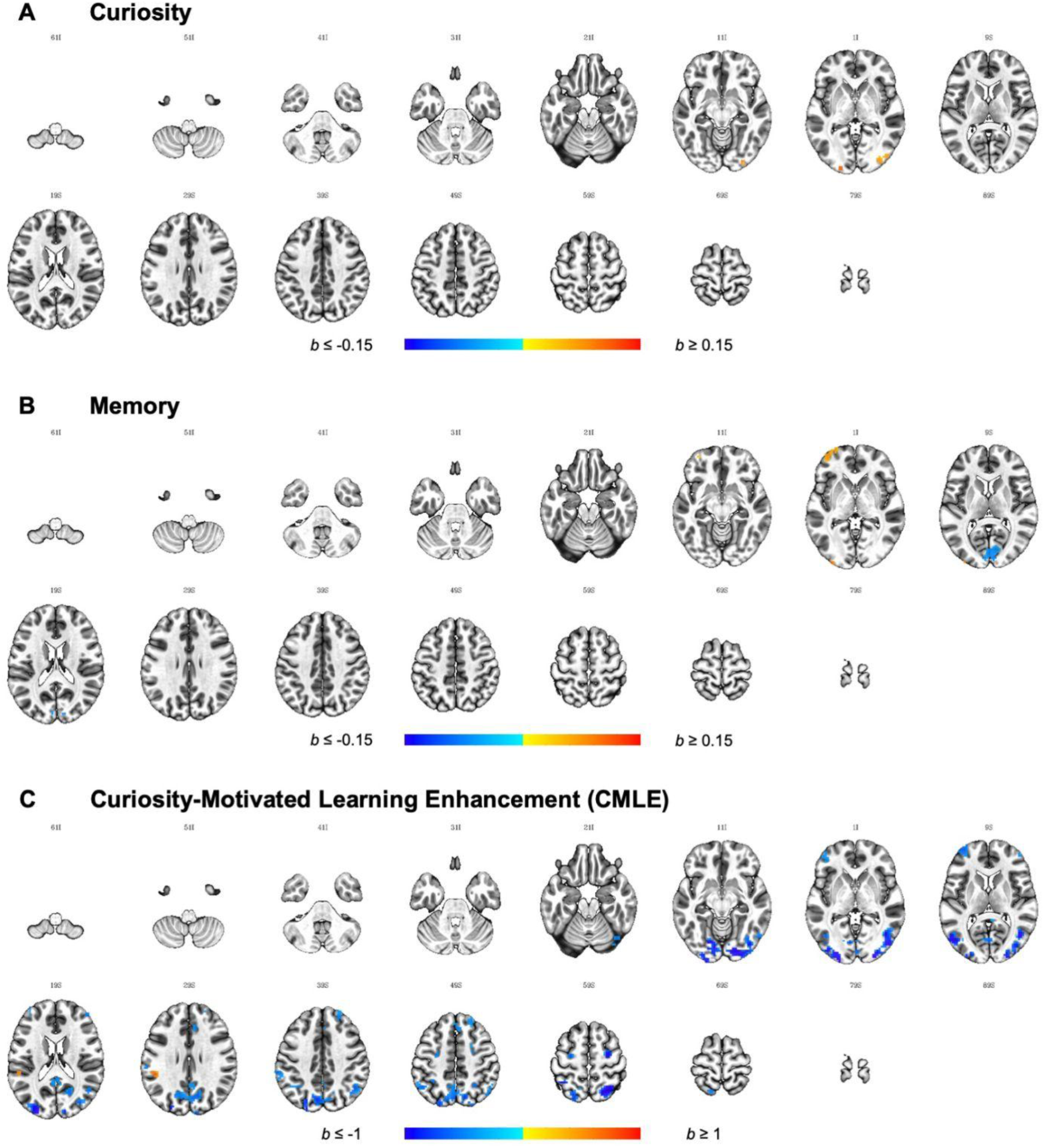
Whole Brain IS-RSA For The Interaction Between The Incentive Manipulation And Each Behavioural Effect Of Interest. *Note*. Results are thresholded at *p* < 0.001, cluster-extent corrected at *k* = 20 (equivalent to per-cluster α = 0.05) and plotted on the ICBM 2009c Nonlinear Asymmetric Template. Images are displayed in neurological orientation, where the left side of the brain is depicted on the left side of the image. Positive clusters (shown in red) indicate more positive values in the control compared to the incentive group whereas negative clusters (shown in blue) indicate more positive values in the incentive group.

##### Memory Incentive Interaction

ROI analysis did not reveal any clusters where incentive influenced how similarity in memory predicted the similarity in the neural response. In the whole brain analysis, three clusters were found (Figure 7B, Table S10) showing a differential predictive effect of similarity in memory depending on the availability of monetary incentives: One cluster in the bilateral Calcarine gyrus showed a more positive predictive effect of memory in the incentive compared to the control group. Two clusters were found where the predictive effect of memory was larger in the control compared to the incentive group. Those were located in the left dorsolateral prefrontal cortex (dlPFC, BA 10/BA 46) and left lateral middle occipital gyrus.

##### Curiosity-Motivated Learning Enhancement (CMLE) Incentive Interaction

While effects of curiosity and memory can be understood in a linear manner, the similarity matrix for CMLE was computed based on a nonlinear AnnaK model formulation further influencing the interpretation of any effects observed. More specifically, positive clusters represent brain regions where CMLE high scorers share similar patterns and low scorers show variability whereas negative clusters represent regions where CMLE low scorers share similar patterns and high scorers show variability.

As with the effects of curiosity and memory, the availability of monetary incentives did not affect the relationship between the similarity in CMLE and brain activity in any of the ROIs. At the whole brain level, 20 clusters were found (Figure 7C, Table S10). One cluster showed positive values indicating that values were more positive in the control compared to the incentive group. This cluster was located in the left supramarginal gyrus where values were negative in the incentive group but weakly positive in the control group. Additionally, 19 clusters showed negative values in which the values in the incentive group were more positive compared to the control group. These clusters were predominantly located in posterior regions, stretching from the occipital poles towards the temporo-parietal-occipital junction laterally and the cuneus medially. In the parietal cortex, clusters were found in the precuneus as well as the superior parietal lobe. Frontally, bilateral clusters in the MFG were found as well as in the right superior frontal gyrus and the superior medial gyrus stretching into the ACC.

## 4 Discussion

The goal of the present study was to examine the effects of curiosity on incidental encoding using different stimuli and a new way to elicit curiosity compared to the well-established trivia question paradigm. Further, we were interested in the combined effects of curiosity and monetary incentives on memory and neural responses. Behavioural results from three experiments showed that curiosity caused by the induced violation of expectations and surprise using magic trick videos facilitated incidental encoding independently of the availability of monetary incentives, but curiosity and monetary incentives did not interact with one another with respect to behavioural measures of learning. fMRI analysis accounting for the dynamic nature of the stimuli revealed that effects of curiosity elicitation, memory encoding, curiosity-motivated learning enhancement (CMLE) as well as monetary incentive effects were associated with activity across widespread cortical areas. Additionally, while the effects of memory encoding and CMLE were supported by activity within the often implicated mesolimbic regions within the hippocampal-VTA loop, we did not find any indication that the effects of curiosity elicitation and monetary incentives were supported by shared, stimulus-induced activity in those regions.

### 4.1 Effects Of Curiosity And Monetary Incentive On Memory

In contrast to the previous studies manipulating monetary reward within the trivia question paradigm (Murayama & Kuhbandner, 2011; Swirsky et al., 2021), we did not find a significant interaction between curiosity and incentive on any of our main measures of interest (recognition, high confidence recognition, cued recall). While the non-significant interaction effect may be explained by the differences in the design (e.g., materials, memory measures, and procedure to manipulate incentives compared to rewards), we also found an interesting dissociation between the effect of curiosity and that of incentives on memory. Specifically, the effects of curiosity on encoding were only found in recollection-based memory measurements (i.e., high confidence recognition and cued recall), but not on recognition regardless of confidence that is assumed to reflect familiarity and recollection (Yonelinas, 2002). On the other hand, the effect of incentives on memory (i.e., high confidence recognition as well as recognition regardless of confidence on trend level) does not seem to be influenced by confidence. Likewise, our exploratory analysis showed that while the effects of curiosit on memory encoding increases as confidence in the recognition task increases, this is not the case for the incentive effects. These findings suggest that curiosity only affects recollection-based, but not familiarity-based processes whereas the influence of monetary rewards is less selective.

These findings were unexpected but on scrutiny of the literature, they were somewhat consistent with findings previously reported. For example, Gruber and colleagues (2014) reported that the curiosity-related recognition advantage in a delayed memory test was specific to confidently recognised faces and did not emerge in overall recognition rates. These results were replicated with short delays (Galli et al., 2018 (Exp. 1, but not in Exp. 2); Murphy, Dehmelt, et al., 2021), and it has been suggested that curiosity-related memory facilitation is specific to recollection (Gruber et al., 2019; Murayama & Elliot, 2011; cf. Stare et al., 2018 for an exception). On the other hand, studies on incentives/rewards and memory have suggested that rewards may influence both recollection and familiarity components of memory (Bunzeck et al., 2010, 2012; Patil et al., 2017; cf. Wittmann et al., 2011). Although not specifically about memory effects, the findings are also consistent with a meta-analysis showing that extrinsic rewards/incentives better predicted *quantity* of performance whereas *quality* was better explained by intrinsic motivation, which is a critical source of curiosity (Cerasoli et al., 2014).

### 4.2 Neural Correlates Of Curiosity-And Incentive-Motivated Learning Within Reward-Related Areas And The Hippocampus

fMRI research on the effects of curiosity (Gruber et al., 2014) and monetary rewards/incentives (Murty & Adcock, 2014; Wittmann et al., 2005, 2008) on incidental encoding has repeatedly implicated the striatum, VTA/SN and hippocampus in motivated learning. Although we found that watching magic tricks led to significant synchronisation of brain activity across subjects in all of these areas, the incentive manipulation did not lead to differential synchronisation in these *a priori* defined ROI (aHPC, VTA/SN, NAcc, and CN). While some of the effects of interest (i.e., memory and CMLE) were located within the ROIs, others (i.e., curiosity) were not. Importantly, the interaction between any effects of interest and monetary incentives were only found outside these brain regions.

The biggest difference between this study and previous studies on the effects of curiosity and monetary incentives/rewards on encoding lies in the nature of stimuli used. Compared to the simplistic, static stimuli used by previous studies (blurred images, trivia questions), magic tricks have added complexity due to their dynamic nature. Critically, we analysed the fMRI data from dynamic stimuli based on intersubject synchronisation (or intersubject correlation (ISC); Hasson et al., 2004), focusing on the intrinsic correlation of the voxel-wise time courses across participants to determine (clusters of) voxels exhibiting a consistent response to the naturalistic stimuli (Nastase et al., 2019). The obtained ISC maps were further contrasted between different types of participant pairs in terms of incentive condition, curiosity rating, memory encoding, and CMLE. As such, the current analysis captures different types of brain dynamics from the classical approach based on the General Linear Model.

For instance, the lack of ISC effects of monetary incentives in reward-related structures does not necessarily imply that there is no difference in brain activation in response to incentives. In fact, it is possible that the activation in reward-related structures was overall increased in the reward compared to the control group, but such an overall increase would not affect the correlation. Using the ISC analysis, we instead tested whether the manipulation of incentives increased or reduced the individual differences in time course pattern within a voxel (e.g., voxels within the reward-related structures). In other words, significant differences in ISC are expected when incentives made participants similarly (or differently) attend and comprehend the magic tricks (Hasson, Furman, et al., 2008), and should manifest in brain areas that are responsible for the synchronised psychological functions (e.g., attention, comprehension). As such, we do not have a strong reason to believe that the reward network responds in an as synchronised fashion. Similar logic should apply to our IS-RSA analysis of the effects of curiosity and memory performance and the incentive effects therein.

An interesting observation from the ROI analysis, however, is that an effect of memory was found in the bilateral CN, replicating previous studies linking declarative memory to the CN (Blumenfeld et al., 2011; Schott, 2006). While meta-analyses have linked the CN to reward processing (Diekhof et al., 2012; Sescousse et al., 2013), the CN has also been implicated in goal-directed action and learning (for a review, see Grahn et al., 2008), and more specifically, in error learning (Delgado et al., 2005) and reward-motivated learning (Wittmann et al., 2005). However, even in the absence of feedback or reward, enhanced activity in the CN has also been found when expectations are violated in a movement observation paradigm (Schiffer & Schubotz, 2011), hence linking the CN to perceptual prediction errors (when “what is happening now” differs from the internally generated prediction; Zacks et al., 2007). Enhanced CN activity has further been found when participants watch magic tricks compared to matched control scenes not violating expectations (Danek et al., 2015), suggesting that magic tricks, because they violate expectations, trigger perceptual prediction errors, signalled in the CN. We here found that similarity in encoding magic tricks predicts similarity in CN activity. This suggests that the CN is not only important in signalling perceptual prediction errors, but might also play a role in updating internal models, or schemata, by supporting the encoding of incongruent events (see also exploratory intersubject functional connectivity analysis (ISFC; Simony et al., 2016) to support this view).

Lastly, significant CMLE effects were observed in all four ROIs, but importantly, these effects were negative. Negative clusters essentially indicate that participants who have low beta values (i.e., participants in which curiosity did not predict memory performance) showed more similar brain activation time courses in response to the magic trick stimuli. Put differently, in negative clusters, the response in the low scorers suggests a more exogenous and stimulus-driven process whereas the response in high scorers is likely more endogenous and individual-based — participants who have a high curiosity-memory association have more divergent and diverse time courses between individuals. Using the trivia question paradigm, Gruber and colleagues (2014) were the first to link the effects of curiosity on incidental encoding (i.e., the interaction between curiosity and memory) to activity in the bilateral NAcc and the right HPC (but not the left). Likewise, activity in the CN and NAcc supports the effects of curiosity on intentional encoding (Duan et al., 2020). The current study suggests that these brain areas are involved in curiosity-based memory encoding, but likely in a more time-varying and idiosyncratic manner.

### 4.3 Curiosity-And Incentive-Motivated Learning Outside The Reward-Related Areas And The Hippocampus

In addition to the results within the *a priori* ROIs (the reward-related areas and the hippocampus), our whole brain IS-RSA showed broader networks of the brain supporting curiosity, memory, and curiosity-motivated learning enhancement (CMLE) than previously implicated. While a more extended discussion of these results can be found in the supplementary material, certain observations deserve attention.

#### 4.3.1 Curiosity

With respect to the effects of curiosity elicitation, we found that similarity in the curiosity ratings predicted similarity in the brain response in visual areas, the inferior frontal gyrus (IFG), the supplementary motor area (SMA), the postcentral gyrus, precuneus, anterior insula, and the supramarginal gyrus in the IPL. While initial fMRI research using the trivia question paradigm suggested that curiosity - operationalised as the anticipation of rewarding information - is supported in dopaminergic regions in the striatum and midbrain (Gruber et al., 2014; Kang et al., 2009), the elicitation of curiosity has more recently been linked to a state of uncertainty, potentially due to a violation of expectations (Gruber & Ranganath, 2019; Murayama et al., 2019). Indeed, both the SMA (Cheung et al., 2019; Volz et al., 2005) and the anterior insula (Grinband et al., 2006; Huettel et al., 2005, 2006; Volz et al., 2003) have been implicated in the processing of uncertainty (for an extended discussion of the role of the anterior insula, please see the supplementary material). The IPL has previously been linked to signalling the moment of expectation violation in magic tricks (Danek et al., 2015), the induction of curiosity in a lottery task (van Lieshout et al., 2018) as well as within the trivia question paradigm (Duan et al., 2020; Ligneul et al., 2018), and even more broadly to knowledge uncertainty (Volz et al., 2004). Likewise, the IFG has previously been implicated in the elicitation of curiosity within the trivia question paradigm (Gruber et al., 2014; Kang et al., 2009) and might be involved in the appraisal processes determining whether prediction error and associated uncertainty elicits curiosity or anxiety (Gruber & Ranganath, 2019). The IFG has also been linked to the violation of expectations (Danek et al., 2015) and causal relationships (Parris et al., 2009) in magic tricks. This suggests that as participants watch magic tricks, the curiosity IS-RSA effects reported here could reflect that uncertainty-related signals and their appraisal processes of the experienced prediction errors share a similar signature when experienced curiosity is shared across individuals.

#### 4.3.2 Memory

The IS-RSA memory effect was found in broadly distributed cortical areas including visual cortices, medial and lateral parietal lobe, lateral temporal areas and dorsal PFC. Overall, the IS-RSA memory effects reported here replicate previous findings obtained using dynamic stimuli. Nguyen and colleagues (2019) also used IS-RSA to anchor similarity in encoding (immediate recall) to similarity in brain response and reported clusters in the default mode network (DMN; e.g., angular gyrus) and the fronto-parietal network (FPN; e.g., bilateral MFG). Both networks have been found to be co-activated during movie watching which could be related to mentalizing, emotional processes and social reasoning (Dixon et al., 2018; Nguyen et al., 2019). This is in alignment with previous studies implicating the DMN in the processing of complex narratives, suggesting that the DMN might accumulate information over longer time scales to integrate them at higher levels of the processing hierarchy (Hasson et al., 2015; Hasson, Yang, et al., 2008). The FPN, on the other hand, supports top-down adaptive, online control (Dosenbach et al., 2007). This suggests that adaptive control mechanisms in the FPN triggered by the cognitive conflict caused by observing the violation of causal relationships in magic tricks could facilitate successful encoding thereof.

#### 4.3.3 Curiosity-Motivated Learning Enhancement (CMLE)

In addition to the ROI results discussed above, significant CMLE effects were observed in broad cortical areas but, importantly, these effects were mostly negative. Indeed, negative clusters were found across largely distributed cortical and subcortical areas including large parts of the DMN (e.g., bilateral ACC, angular gyrus, middle temporal gyrus), FPN (e.g., bilateral MFG, SMA), dorsal attention network (e.g., bilateral posterior superior parietal lobe), ventral attention network (e.g., anterior insula/frontal operculum (aI/fO)) as well as visual network. A recent re-analysis of the dataset from Gruber and colleagues (2014) showed that the DMN and a subnetwork within the FPN (i.e., lateral PFC, posterior inferior temporal gyrus, and superior parietal lobe) show a curiosity-by-memory interaction (Murphy, Ranganath, et al., 2021). Replicating and expanding on these results and in alignment with the ROI analysis discussed above, we found that participants with high CMLE compared to those with low scores show a more individualised and variable activation in these brain networks (for a discussion of the implications of these findings, please refer to the supplementary material).

### 4.4 Influences Of The Availability Of Extrinsic Incentives

While we did not find effects of extrinsic incentives in the hippocampal-VTA loop or other dopaminergic regions, effects were found at the whole brain level. For instance, we found that monetary incentives are associated with increased synchronisation in the early visual areas, the postcentral gyrus and the intraparietal sulcus (IPS). Overall, these areas have been implicated in top-down spatial attention and form the dorsal attention network (Corbetta & Shulman, 2002; Vossel et al., 2014) in general, but also in value-driven attentional processes in the context of monetary rewards (Anderson, 2017) and memory-guided attention (Salsano et al., 2021) in particular. This suggests that participants in the incentive group might have attended to the magic tricks in a more similar manner because they have the common goal of maximising their rewards in the judgement task, perhaps attending to similar parts of the video clip. In the control group, on the other hand, increased ISC was found in Area V5/MT+ which is critically involved in motion perception and has feed-forward connections to the parietal cortex (Zeki, 2015). Likewise, the IPS is also more activated during the encoding of items compared to their associations (Kim, 2011), suggesting that the IPS supports familiarity-based compared to recollection-based recognition. Hence, the difference in ISC depending on the availability of monetary incentives provides further support for the proposition regarding the dissociation between curiosity and monetary incentives discussed above.

We further found IS-RSA clusters for each behavioural effect of interest where the second-order similarity between behaviour and brain response differed depending on the availability of monetary incentives, suggesting that the incentive manipulation influenced not only the degree of synchronisation during the naturalistic viewing per se, but also the way that similarity in the stimulus-elicited brain response could be anchored to similarity in each of the behavioural variables of interest. The clusters showing an interaction between the behavioural effects of interest and the incentive manipulation are primarily located in the visual or dorsal attention network, further suggesting that the incentive manipulation influenced attentional mechanisms during incidental encoding. Of note, for the memory-incentive interaction, we found lower synchronisation in the incentive group compared to the control group on the left lateral PFC, an area in the FPN previously implicated in the neural basis of the undermining effect (Murayama et al., 2010), but also in memory effects in general (Kim, 2011). Overall, our analysis showed that CMLE in and of itself is supported by endogenous rather than exogenous responses to the magic tricks in participants with high CMLE scores compared to those with low scores (reflected in negative effects). Investigating the interaction between CMLE and the incentive manipulation, most clusters were negative, meaning that the values in the incentive group were more positive compared to the control group. This suggests that in the incentive groups, participants’ tendency to prioritise internal over external processes was less pronounced in support of curiosity-motivated learning.

### 4.5 Overall Conclusion

We found that the curiosity effect of memory can be replicated using naturalistic stimuli. Using analysis approaches to account for the dynamic nature of the magic trick stimuli, the effects of curiosity and incentives per se were not located within the reward network of the brain, but across distributed cortical areas. While effects of memory and CMLE were found within the hippocampal-VTA loop, they too showed widely distributed cortical clusters. This supports the claim that the effects of motivated incidental encoding of dynamic stimuli are actually more distributed across higher-order cortices and not solely based on mesolimbic structures, often identified using reductionist simple stimuli that do not reflect everyday perception and cognition and analysis approaches based on rigorous modelling of the hemodynamic response. This suggests that a too stringent focus on narrow ROIs could lead to an oversimplification and might miss important insights in how the brain works when processing and encoding naturalistic stimuli. To derive a better understanding on how curiosity influences memory and to inform practitioners in educational settings, more research with various stimuli and tasks is needed.

## Footnotes

1 In previous literature on motivated learning, the terms ‘rewards’ and ‘incentives’ have been used rather interchangeably (e.g., Adcock et al., 2006), but some attach distinct definitions to them. Specifically, incentives are ‘plans that have predetermined criteria and standards, as well as understood policies for determining and allocating rewards’ (Greene, 2010, p. 219). As such, incentives can be seen as a promise of later rewards, hence incentives can be seen as expected rewards (Berridge, 2000), whereas rewards are the outcome of motivated behaviour that are received/perceived/consumed (Matyjek et al., 2020; Schultz, 2015). In this paper, we adopt these differential definitions.

2 Due to an error, one item was not included into the inventory. The pressure scale was computed based on 4 instead of 5 items.

3 50% additional bonus payment should have translated to £0.40 per correct answer. However, no participant reported to have noticed this error.

4 Due to singular fit warnings for the dependent variable high confidence recognition in the fMRI data, the model was also executed using a simplified random effects structure where the random intercepts of subject and random slopes of subjects for the curiosity effect were specified, but random intercepts of stimuli were removed allowing the model to converge without warnings. The individual curiosity beta values from both models were highly correlated (r = 0.992) and the gLME model specification did not affect the IS-RSA whole brain results (correlation unthresholded effect size map = .996, correlation unthresholded statistics map = .997, dice coefficient of masked cluster-extent thresholded results = 0.960) nor reported ROI results.

## 5 Acknowledgements

We are grateful to the Centre for Integrative Neuroscience and Neurodynamics (CINN) and the MeMo Lab for helpful discussions and support in the realisation of this project. We would like to especially thank Cristina Pascua Martin for her support during the fMRI data collection and with respect to the coding of recall answers. We would like to express our gratitude to AFNI Team and Jeremy Skipper for their input regarding the design of the fMRI experiment, pre-processing, and analysis. Lastly, we would like to acknowledge Anthony Haffey’s ongoing support with regards to data collection using Collector. This research was supported by Leverhulme Trust Research Leadership Award (RL-2016-030), Jacobs Foundation Advanced Research Fellowship, and the Alexander von Humboldt Foundation (the Alexander von Humboldt Professorship endowed by the German Federal Ministry of Education and Research) to Kou Murayama.

## 6 CRediT Author Statement

**Stefanie Meliss:** Conceptualisation, Methodology, Software, Validation, Formal analysis, Investigation, Resources, Data Curation, Writing – Original Draft, Visualisation, Project administration. **Carien van Reekum:** Conceptualisation, Writing – Review & Editing, Supervision. **Kou Murayama:** Conceptualisation, Methodology, Resources, Writing – Review & Editing, Supervision, Project Administration, Funding acquisition.

## 7 Data and Code Availability Statement

Scripts used to process and analyse the data were uploaded to GitHub (https://github.com/stefaniemeliss/MIL_paper). The repository further contains the behavioural data. Raw and pre-processed fMRI were uploaded on OpenNeuro (https://doi.org/10.18112/openneuro.ds004182.v1.0.0). ROI masks and unthresholded fMRI results at group-level are available as a NeuroVault collection (https://neurovault.org/collections/12980/).

## References

Adcock, R. A., Thangavel, A., Whitfield-Gabrieli, S., Knutson, B., & Gabrieli, J. D. E. (2006). Reward-Motivated Learning: Mesolimbic Activation Precedes Memory Formation. Neuron, 50(3), 507–517.

Aliko, S., Huang, J., Gheorghiu, F., Meliss, S., & Skipper, J. I. (2020). A naturalistic neuroimaging database for understanding the brain using ecological stimuli. Scientific Data, 7(1), 347.

Anderson, B. A. (2017). Reward processing in the value-driven attention network: reward signals tracking cue identity and location. Social Cognitive and Affective Neuroscience, 12(3), 461–467.

Baldassano, C., Chen, J., Zadbood, A., Pillow, J. W., Hasson, U., & Norman, K. A. (2017). Discovering Event Structure in Continuous Narrative Perception and Memory. Neuron, 95(3), 709–721.e5.

Bates, D., Mächler, M., Bolker, B., & Walker, S. (2015). Fitting Linear Mixed-Effects Models Using lme4. Journal of Statistical Software, Articles, 67(1), 1–48.

Bennett, D., Bode, S., Brydevall, M., Warren, H., & Murawski, C. (2016). Intrinsic Valuation of Information in Decision Making under Uncertainty. PLoS Computational Biology, 12(7), e1005020.

Berridge, K. C. (2000). Reward learning: Reinforcement, incentives, and expectations. In Psychology of Learning and Motivation (Vol. 40, pp. 223–278). Academic Press.

Blumenfeld, R. S., Parks, C. M., Yonelinas, A. P., & Ranganath, C. (2011). Putting the pieces together: the role of dorsolateral prefrontal cortex in relational memory encoding. Journal of Cognitive Neuroscience, 23(1), 257–265.

Brainard, D. H. (1997). The Psychophysics Toolbox. Spatial Vision, 10(4), 433–436.

Brod, G., & Breitwieser, J. (2019). Lighting the wick in the candle of learning: generating a prediction stimulates curiosity. NPJ Science of Learning, 4, 1–7.

Brod, G., Hasselhorn, M., & Bunge, S. A. (2018). When generating a prediction boosts learning: The element of surprise. Learning and Instruction, 55, 22–31.

Bromberg-Martin, E. S., & Hikosaka, O. (2009). Midbrain dopamine neurons signal preference for advance information about upcoming rewards. Neuron, 63(1), 119–126.

Brydevall, M., Bennett, D., Murawski, C., & Bode, S. (2018). The neural encoding of information prediction errors during non-instrumental information seeking. Scientific Reports, 8(1), 6134.

Bunzeck, N., Dayan, P., Dolan, R. J., & Duzel, E. (2010). A common mechanism for adaptive scaling of reward and novelty. Human Brain Mapping, 31(9), 1380–1394.

Bunzeck, N., Doeller, C. F., Dolan, R. J., & Duzel, E. (2012). Contextual interaction between novelty and reward processing within the mesolimbic system. Human Brain Mapping, 33(6), 1309–1324.

Cen, D., Gkoumas, C., & Gruber, M. J. (2021). Anticipation of novel environments enhances memory for incidental information. Learning & Memory, 28(8), 254–259.

Chen, G., Taylor, P. A., Shin, Y. W., Reynolds, R. C., & Cox, R. W. (2017). Untangling the relatedness among correlations, Part II: Inter-subject correlation group analysis through linear mixed-effects modeling. NeuroImage, 147(August 2016), 825–840.

Chen, J., Leong, Y. C., Honey, C. J., Yong, C. H., Norman, K. A., & Hasson, U. (2017). Shared memories reveal shared structure in neural activity across individuals. Nature Neuroscience, 20(1), 115–125.

Cheung, V. K. M., Harrison, P. M. C., Meyer, L., Pearce, M. T., Haynes, J.-D., & Koelsch, S. (2019). Uncertainty and Surprise Jointly Predict Musical Pleasure and Amygdala, Hippocampus, and Auditory Cortex Activity. Current Biology: CB, 29(23), 4084–4092.e4.

Corbetta, M., & Shulman, G. L. (2002). Control of goal-directed and stimulus-driven attention in the brain. Nature Reviews. Neuroscience, 3(3), 201–215.

Danek, A. H., Öllinger, M., Fraps, T., Grothe, B., & Flanagin, V. L. (2015). An fMRI investigation of expectation violation in magic tricks. Frontiers in Psychology, 6, 84.

Delgado, M. R., Miller, M. M., Inati, S., & Phelps, E. A. (2005). An fMRI study of reward-related probability learning. NeuroImage, 24(3), 862–873.

Diekhof, E. K., Kaps, L., Falkai, P., & Gruber, O. (2012). The role of the human ventral striatum and the medial orbitofrontal cortex in the representation of reward magnitude - An activation likelihood estimation meta-analysis of neuroimaging studies of passive reward expectancy and outcome processing. Neuropsychologia, 50(7), 1252–1266.

Dixon, M. L., De La Vega, A., Mills, C., Andrews-Hanna, J., Spreng, R. N., Cole, M. W., & Christoff, K. (2018). Heterogeneity within the frontoparietal control network and its relationship to the default and dorsal attention networks. Proceedings of the National Academy of Sciences of the United States of America, 115(7), E1598–E1607.

Dosenbach, N. U. F., Fair, D. A., Miezin, F. M., Cohen, A. L., Wenger, K. K., Dosenbach, R. A. T., Fox, M. D., Snyder, A. Z., Vincent, J. L., Raichle, M. E., Schlaggar, B. L., & Petersen, S. E. (2007). Distinct brain networks for adaptive and stable task control in humans. Proceedings of the National Academy of Sciences of the United States of America, 104(26), 11073–11078.

Duan, H., Fernández, G., van Dongen, E., & Kohn, N. (2020). The effect of intrinsic and extrinsic motivation on memory formation: insight from behavioral and imaging study. Brain Structure & Function, 225(5), 1561–1574.

Elliot, A. J., & Harackiewicz, J. M. (1996). Approach and avoidance achievement goals and intrinsic motivation: A mediational analysis. Journal of Personality and Social Psychology, 70(3), 461–475.

Fastrich, G. M., Kerr, T., Castel, A. D., & Murayama, K. (2018). The role of interest in memory for trivia questions: An investigation with a large-scale database. Motivation Science, 4(3), 227–250.

Finn, E. S., Corlett, P. R., Chen, G., Bandettini, P. A., & Constable, R. T. (2018). Trait paranoia shapes inter-subject synchrony in brain activity during an ambiguous social narrative. Nature Communications, 9(1), 2043.

Finn, E. S., Glerean, E., Khojandi, A. Y., Nielson, D., Molfese, P. J., Handwerker, D. A., & Bandettini, P. A. (2020). Idiosynchrony: From shared responses to individual differences during naturalistic neuroimaging. NeuroImage, 215, 116828.

FitzGibbon, L., Komiya, A., & Murayama, K. (2021). The Lure of Counterfactual Curiosity: People Incur a Cost to Experience Regret. Psychological Science, 32(2), 241–255.

FitzGibbon, L., Lau, J. K. L., & Murayama, K. (2020). The seductive lure of curiosity: information as a motivationally salient reward. Current Opinion in Behavioral Sciences, 35, 21–27.

Galli, G., Sirota, M., Gruber, M. J., Ivanof, B. E., Ganesh, J., Materassi, M., Thorpe, A., Loaiza, V., Cappelletti, M., & Craik, F. I. M. (2018). Learning facts during aging: the benefits of curiosity. Experimental Aging Research, 44(4), 311–328.

Glasser, M. F., Coalson, T. S., Robinson, E. C., Hacker, C. D., Harwell, J., Yacoub, E., Ugurbil, K., Andersson, J., Beckmann, C. F., Jenkinson, M., Smith, S. M., & Van Essen, D. C. (2016). A multi-modal parcellation of human cerebral cortex. Nature, 536(7615), 171–178.

Grahn, J. A., Parkinson, J. A., & Owen, A. M. (2008). The cognitive functions of the caudate nucleus. Progress in Neurobiology, 86(3), 141–155.

Greene, R. J. (2010). Rewarding performance: Guiding principles, custom strategies. Routledge.

Grinband, J., Hirsch, J., & Ferrera, V. P. (2006). A neural representation of categorization uncertainty in the human brain. Neuron, 49(5), 757–763.

Gruber, M. J., Gelman, B. D., & Ranganath, C. (2014). States of curiosity modulate hippocampus-dependent learning via the dopaminergic circuit. Neuron, 84(2), 486–496.

Gruber, M. J., & Otten, L. J. (2010). Voluntary control over prestimulus activity related to encoding. The Journal of Neuroscience: The Official Journal of the Society for Neuroscience, 30(29), 9793–9800.

Gruber, M. J., & Ranganath, C. (2019). How Curiosity Enhances Hippocampus-Dependent Memory: The Prediction, Appraisal, Curiosity, and Exploration (PACE) Framework. Trends in Cognitive Sciences, 23(12), 1014–1025.

Gruber, M. J., Ritchey, M., Wang, S. F., Doss, M. K., & Ranganath, C. (2016). Post-learning Hippocampal Dynamics Promote Preferential Retention of Rewarding Events. Neuron, 89(5), 1110–1120.

Gruber, M. J., Valji, A., & Ranganath, C. (2019). Curiosity and learning: a neuroscientific perspective. In K. A. Renniger & S. Hidi (Eds.), The Cambridge Handbook of Motivation and Learning (pp. 397–417). Cambridge University Press.

Gruber, M. J., Watrous, A. J., Ekstrom, A. D., Ranganath, C., & Otten, L. J. (2013). Expected reward modulates encoding-related theta activity before an event. NeuroImage, 64(1), 68–74.

Haffey, A., Plat, K. T., Mane, P., Blake, A., & Chakrabarti, B. (2020). Open source online behavioural experimentation using Collector: Proof of principle & sample size considerations. In PsyArXiv. https://doi.org/10.31234/osf.io/u3saf

Halamish, V., Madmon, I., & Moed, A. (2019). Motivation to Learn. The Long-Term Mnemonic Benefit of Curiosity in Intentional Learning. Experimentational Psychology, 66(5), 319–330.

Hasson, U., Chen, J., & Honey, C. J. (2015). Hierarchical process memory: Memory as an integral component of information processing. Trends in Cognitive Sciences, 19(6), 304– 313.

Hasson, U., Furman, O., Clark, D., Dudai, Y., & Davachi, L. (2008). Enhanced Intersubject Correlations during Movie Viewing Correlate with Successful Episodic Encoding. Neuron, 57(3), 452–462.

Hasson, U., Nir, Y., Levy, I., Fuhrmann, G., & Malach, R. (2004). Intersubject Synchronisation of Cortical Activity During Natural Vision. Science, 303(5664), 1634– 1640.

Hasson, U., Yang, E., Vallines, I., Heeger, D. J., & Rubin, N. (2008). A hierarchy of temporal receptive windows in human cortex. The Journal of Neuroscience: The Official Journal of the Society for Neuroscience, 28(10), 2539–2550.

Hawking, S. (2016). A Brief History Of Time: From The Big Bang To Black Holes (Updated Version). Penguin Random House UK.

Hidi, S., & Renninger, K. A. (2006). The Four-Phase Model of Interest Development. Educational Psychologist, 41(2), 111–127.

Huettel, S. A., Song, A. W., & McCarthy, G. (2005). Decisions under uncertainty: probabilistic context influences activation of prefrontal and parietal cortices. The Journal of Neuroscience: The Official Journal of the Society for Neuroscience, 25(13), 3304– 3311.

Huettel, S. A., Stowe, C. J., Gordon, E. M., Warner, B. T., & Platt, M. L. (2006). Neural signatures of economic preferences for risk and ambiguity. Neuron, 49(5), 765–775.

Jääskeläinen, I. P., Pajula, J., Tohka, J., Lee, H. J., Kuo, W. J., & Lin, F. H. (2016). Brain hemodynamic activity during viewing and re-viewing of comedy movies explained by experienced humor. Scientific Reports, 6(May), 1–14.

Jach, H. K., DeYoung, C. G., & Smillie, L. D. (2022). Why do people seek information? The role of personality traits and situation perception. Journal of Experimental Psychology. General, 151(4), 934–959.

Jepma, M., Verdonschot, R. G., van Steenbergen, H., Rombouts, S. A. R. B., & Nieuwenhuis, S. (2012). Neural mechanisms underlying the induction and relief of perceptual curiosity. Frontiers in Behavioral Neuroscience, 6, 5.

Kang, M. J., Hsu, M., Krajbich, I. M., Loewenstein, G., McClure, S. M., Wang, J. T.-Y., & Camerer, C. F. (2009). The wick in the candle of learning: epistemic curiosity activates reward circuitry and enhances memory. Psychological Science, 20(8), 963–973.

Keysers, C., Kaas, J. H., & Gazzola, V. (2010). Somatosensation in social perception. Nature Reviews. Neuroscience, 11(6), 417–428.

Kim, H. (2011). Neural activity that predicts subsequent memory and forgetting: a meta-analysis of 74 fMRI studies. NeuroImage, 54(3), 2446–2461.

Kobayashi, K., & Hsu, M. (2019). Common neural code for reward and information value. Proceedings of the National Academy of Sciences of the United States of America, 116(26), 13061–13066.

Kobayashi, K., Ravaioli, S., Baranès, A., Woodford, M., & Gottlieb, J. (2019). Diverse motives for human curiosity. Nature Human Behaviour, 3(6), 587–595.

Kriegeskorte, N., Mur, M., & Bandettini, P. (2008). Representational similarity analysis - connecting the branches of systems neuroscience. Frontiers in Systems Neuroscience, 2, 4.

Kuhn, G., Amlani, A. A., & Rensink, R. A. (2008). Towards a science of magic. Trends in Cognitive Sciences, 12(9), 349–354.

Lau, J. K. L., Ozono, H., Kuratomi, K., Komiya, A., & Murayama, K. (2020). Shared striatal activity in decisions to satisfy curiosity and hunger at the risk of electric shocks. Nature Human Behaviour, 4(5), 531–543.

Ligneul, R., Mermillod, M., & Morisseau, T. (2018). From relief to surprise: Dual control of epistemic curiosity in the human brain. NeuroImage, 181, 490–500.

Lisman, J. E., & Grace, A. A. (2005). The Hippocampal-VTA Loop: Controlling the Entry of Information into Long-Term Memory. Neuron, 46(5), 703–713.

Lisman, J. E., Grace, A. A., & Duzel, E. (2011). A neoHebbian framework for episodic memory; role of dopamine-dependent late LTP. Trends in Neurosciences, 34(10), 536– 547.

Marvin, C. B., & Shohamy, D. (2016). Curiosity and reward: Valence predicts choice and information prediction errors enhance learning. Journal of Experimental Psychology. General, 145(3), 266–272.

Matyjek, M., Meliss, S., Dziobek, I., & Murayama, K. (2020). A Multidimensional View on Social and Non-Social Rewards. Frontiers in Psychiatry / Frontiers Research Foundation, 11, 818.

Meliss, S., Pascua, C., Skipper, J. I., & Murayama, K. (2022). The Magic, Memory, and Curiosity fMRI Dataset of People Viewing Magic Tricks. https://doi.org/10.31234/osf.io/zq7gv

Miendlarzewska, E. A., Bavelier, D., & Schwartz, S. (2016). Influence of reward motivation on human declarative memory. Neuroscience and Biobehavioral Reviews, 61, 156–176.

Moraczewski, D., Chen, G., & Redcay, E. (2018). Inter-subject synchrony as an index of functional specialization in early childhood. Scientific Reports, 8(1), 1–12.

Mullaney, K. M., Carpenter, S. K., Grotenhuis, C., & Burianek, S. (2014). Waiting for feedback helps if you want to know the answer: the role of curiosity in the delay-of-feedback benefit. Memory and Cognition, 42(8), 1273–1284.

Murayama, K. (2022). A reward-learning framework of knowledge acquisition: An integrated account of curiosity, interest, and intrinsic-extrinsic rewards. Psychological Review, 129(1), 175–198.

Murayama, K., & Elliot, A. J. (2011). Achievement motivation and memory: achievement goals differentially influence immediate and delayed remember-know recognition memory. Personality & Social Psychology Bulletin, 37(10), 1339–1348.

Murayama, K., FitzGibbon, L., & Sakaki, M. (2019). Process Account of Curiosity and Interest: A Reward-Learning Perspective. Educational Psychology Review, 31(4), 875– 895.

Murayama, K., & Kitagami, S. (2014). Consolidation power of extrinsic rewards: reward cues enhance long-term memory for irrelevant past events. Journal of Experimental Psychology. General, 143(1), 15–20.

Murayama, K., & Kuhbandner, C. (2011). Money enhances memory consolidation - But only for boring material. Cognition, 119(1), 120–124.

Murayama, K., Matsumoto, M., Izuma, K., & Matsumoto, K. (2010). Neural basis of the undermining effect of monetary reward on intrinsic motivation. Proceedings of the National Academy of Sciences, 107(49), 20911–20916.

Murphy, C., Dehmelt, V., Yonelinas, A. P., Ranganath, C., & Gruber, M. J. (2021). Temporal proximity to the elicitation of curiosity is key for enhancing memory for incidental information. Learning & Memory, 28(2), 34–39.

Murphy, C., Ranganath, C., & Gruber, M. J. (2021). Connectivity between the hippocampus and default mode network during the relief – but not elicitation – of curiosity supports curiosity-enhanced memory enhancements. In bioRxiv (p. 2021.07.26.453739). https://doi.org/10.1101/2021.07.26.453739

Murty, V. P., & Adcock, R. A. (2014). Enriched encoding: reward motivation organizes cortical networks for hippocampal detection of unexpected events. Cerebral Cortex, 24(8), 2160–2168.

Nastase, S. A., Gazzola, V., Hasson, U., & Keysers, C. (2019). Measuring shared responses across subjects using intersubject correlation. Social Cognitive and Affective Neuroscience, 14(6), 669–687.

Nguyen, M., Vanderwal, T., & Hasson, U. (2019). Shared understanding of narratives is correlated with shared neural responses. NeuroImage, 184, 161–170.

Nummenmaa, L., Glerean, E., Viinikainen, M., Jääskeläinen, I. P., Hari, R., & Sams, M. (2012). Emotions promote social interaction by synchronizing brain activity across individuals. Proceedings of the National Academy of Sciences of the United States of America, 109(24), 9599–9604.

Nummenmaa, L., Lahnakoski, J. M., & Glerean, E. (2018). Sharing the social world via intersubject neural synchronisation. Current Opinion in Psychology, 24, 7–14.

Ozono, H., Komiya, A., Kuratomi, K., Hatano, A., Fastrich, G., Raw, J. A. L., Haffey, A., Meliss, S., Lau, J. K. L., & Murayama, K. (2021). Magic Curiosity Arousing Tricks (MagicCATs): A novel stimulus collection to induce epistemic emotions. Behavior Research Methods, 53(1), 188–215.

Pajula, J., & Tohka, J. (2016). How Many Is Enough? Effect of Sample Size in Inter-Subject Correlation Analysis of fMRI. Computational Intelligence and Neuroscience, 2016, 2094601.

Parris, B. A., Kuhn, G., Mizon, G. A., Benattayallah, A., & Hodgson, T. L. (2009). Imaging the impossible: an fMRI study of impossible causal relationships in magic tricks. NeuroImage, 45(3), 1033–1039.

Patil, A., Murty, V. P., Dunsmoor, J. E., Phelps, E. A., & Davachi, L. (2017). Reward retroactively enhances memory consolidation for related items. Learning & Memory, 24(1), 65–69.

Pauli, W. M., Nili, A. N., & Michael Tyszka, J. (2018). A high-resolution probabilistic in vivo atlas of human subcortical brain nuclei. Scientific Data, 5(1), 1–13.

Pekrun, R., Goetz, T., Titz, W., & Perry, R. P. (2002). Academic Emotions in Students’ Self-Regulated Learning and Achievement: A Program of Qualitative and Quantitative Research. Educational Psychologist, 37(2), 91–105.

Poh, J.-H., Vu, M.-A. T., Stanek, J. K., Hsiung, A., Egner, T., & Alison Adcock, R. (2021). Tuned to Learn: An anticipatory hippocampal convergence state conducive to memory formation revealed during midbrain activation. In bioRxiv (p. 2021.07.15.452391). https://doi.org/10.1101/2021.07.15.452391

Poppenk, J., Evensmoen, H. R., Moscovitch, M., & Nadel, L. (2013). Long-axis specialization of the human hippocampus. Trends in Cognitive Sciences, 17(5), 230–240.

R Core Team. (2020). R: A language and environment for statistical computing.

Rensink, R. A., & Kuhn, G. (2014). A framework for using magic to study the mind. Frontiers in Psychology, 5, 1508.

Ren, Y., Nguyen, V. T., Guo, L., & Guo, C. C. (2017). Inter-subject Functional Correlation Reveal a Hierarchical Organization of Extrinsic and Intrinsic Systems in the Brain. Scientific Reports, 7(1), 1–12.

Ryan, R. M. (1982). Control and information in the intrapersonal sphere: An extension of cognitive evaluation theory. Journal of Personality and Social Psychology, 43(3), 450– 461.

Salsano, I., Santangelo, V., & Macaluso, E. (2021). The lateral intraparietal sulcus takes viewpoint changes into account during memory-guided attention in natural scenes. Brain Structure & Function, 226(4), 989–1006.

Schiffer, A.-M., & Schubotz, R. I. (2011). Caudate nucleus signals for breaches of expectation in a movement observation paradigm. Frontiers in Human Neuroscience, 5, 38.

Schott, B. H. (2006). The Dopaminergic Midbrain Participates in Human Episodic Memory Formation: Evidence from Genetic Imaging. Journal of Neuroscience, 26(5), 1407– 1417.

Schultz, W. (2015). Neuronal reward and decision signals: From theories to data. Physiological Reviews, 95(3), 853–951.

Sescousse, G., Caldú, X., Segura, B., & Dreher, J.-C. (2013). Processing of primary and secondary rewards: a quantitative meta-analysis and review of human functional neuroimaging studies. Neuroscience and Biobehavioral Reviews, 37(4), 681–696.

Shamay-Tsoory, S. G., & Mendelsohn, A. (2019). Real-Life Neuroscience: An Ecological Approach to Brain and Behavior Research. Perspectives on Psychological Science: A Journal of the Association for Psychological Science, 14(5), 841–859.

Sharot, T., & Sunstein, C. R. (2020). How people decide what they want to know. Nature Human Behaviour, 4(1), 14–19.

Shohamy, D., & Adcock, R. A. (2010). Dopamine and adaptive memory. Trends in Cognitive Sciences, 14(10), 464–472.

Simony, E., Honey, C. J., Chen, J., Lositsky, O., Yeshurun, Y., Wiesel, A., & Hasson, U. (2016). Dynamic reconfiguration of the default mode network during narrative comprehension. *Nature Communications*, May 2015. https://doi.org/10.1038/ncomms12141

Sonkusare, S., Breakspear, M., & Guo, C. (2019). Naturalistic Stimuli in Neuroscience: Critically Acclaimed. Trends in Cognitive Sciences, 23(8), 699–714.

Spaniol, J., Davidson, P. S. R., Kim, A. S. N., Han, H., Moscovitch, M., & Grady, C. L. (2009). Event-related fMRI studies of episodic encoding and retrieval: meta-analyses using activation likelihood estimation. Neuropsychologia, 47(8-9), 1765–1779.

Stanek, J. K., Dickerson, K. C., Chiew, K. S., Clement, N. J., & Adcock, R. A. (2019). Expected Reward Value and Reward Uncertainty Have Temporally Dissociable Effects on Memory Formation. Journal of Cognitive Neuroscience, 31(10), 1443–1454.

Stare, C. J., Gruber, M. J., Nadel, L., Ranganath, C., & Gómez, R. L. (2018). Curiosity-driven memory enhancement persists over time but does not benefit from post-learning sleep. Cognitive Neuroscience, 9(3-4), 100–115.

Swirsky, L. T., Shulman, A., & Spaniol, J. (2021). The interaction of curiosity and reward on long-term memory in younger and older adults. Psychology and Aging, 36(5), 584–603.

Thomas, R. M., De Sanctis, T., Gazzola, V., & Keysers, C. (2018). Where and how our brain represents the temporal structure of observed action. NeuroImage, 183, 677–697.

van Lieshout, L. L. F., Traast, I. J., de Lange, F. P., & Cools, R. (2021). Curiosity or savouring? Information seeking is modulated by both uncertainty and valence. PloS One, 16(9), e0257011.

van Lieshout, L. L. F., Vandenbroucke, A. R. E., Müller, N. C. J., Cools, R., & de Lange, F. P. (2018). Induction and relief of curiosity elicit parietal and frontal activity. The Journal of Neuroscience: The Official Journal of the Society for Neuroscience, 38(10), 2816– 2817.

Viechtbauer, W. (2010). Conducting Meta-Analyses in R with the metafor Package. *Journal of Statistical Software*, Articles, 36(3), 1–48.

Vogl, E., Pekrun, R., Murayama, K., Loderer, K., & Schubert, S. (2019). Surprise, Curiosity, and Confusion Promote Knowledge Exploration: Evidence for Robust Effects of Epistemic Emotions. Frontiers in Psychology, 10, 2474.

Volz, K. G., Schubotz, R. I., & von Cramon, D. Y. (2003). Predicting events of varying probability: uncertainty investigated by fMRI. NeuroImage, *19*(2 Pt 1), 271–280.

Volz, K. G., Schubotz, R. I., & von Cramon, D. Y. (2004). Why am I unsure? Internal and external attributions of uncertainty dissociated by fMRI. NeuroImage, 21(3), 848–857.

Volz, K. G., Schubotz, R. I., & von Cramon, D. Y. (2005). Variants of uncertainty in decision-making and their neural correlates. Brain Research Bulletin, 67(5), 403–412.

Vossel, S., Geng, J. J., & Fink, G. R. (2014). Dorsal and ventral attention systems: distinct neural circuits but collaborative roles. *The Neuroscientist: A Review Journal Bringing Neurobiology*, Neurology and Psychiatry, 20(2), 150–159.

Wade, S., & Kidd, C. (2019). The role of prior knowledge and curiosity in learning. Psychonomic Bulletin & Review, 26(4), 1377–1387.

Wigfield, A., & Eccles, J. S. (2000). Expectancy–Value Theory of Achievement Motivation. Contemporary Educational Psychology, 25(1), 68–81.

Wittmann, B. C., Dolan, R. J., & Duzel, E. (2011). Behavioral specifications of reward-associated long-term memory enhancement in humans. Learning & Memory, 18(5), 296– 300.

Wittmann, B. C., Schiltz, K., Boehler, C. N., & Düzel, E. (2008). Mesolimbic interaction of emotional valence and reward improves memory formation. Neuropsychologia, 46(4), 1000–1008.

Wittmann, B. C., Schott, B. H., Guderian, S., Frey, J. U., Heinze, H.-J., & Düzel, E. (2005). Reward-related FMRI activation of dopaminergic midbrain is associated with enhanced hippocampus-dependent long-term memory formation. Neuron, 45(3), 459–467.

Wolosin, S. M., Zeithamova, D., & Preston, A. R. (2012). Reward modulation of hippocampal subfield activation during successful associative encoding and retrieval. Journal of Cognitive Neuroscience, 24(7), 1532–1547.

Woo, C. W., Krishnan, A., & Wager, T. D. (2014). Cluster-extent based thresholding in fMRI analyses: Pitfalls and recommendations. NeuroImage, 91, 412–419.

Yeshurun, Y., Swanson, S., Simony, E., Chen, J., Lazaridi, C., Honey, C. J., & Hasson, U. (2017). Same Story, Different Story: The Neural Representation of Interpretive Frameworks. Psychological Science, 28(3), 307–319.

Yonelinas, A. P. (2001). Components of episodic memory: the contribution of recollection and familiarity. Philosophical Transactions of the Royal Society of London. Series B, Biological Sciences, 356(1413), 1363–1374.

Yonelinas, A. P. (2002). The Nature of Recollection and Familiarity: A Review of 30 Years of Research. Journal of Memory and Language, 46(3), 441–517.

Zacks, J. M., Speer, N. K., Swallow, K. M., Braver, T. S., & Reynolds, J. R. (2007). Event perception: a mind-brain perspective. Psychological Bulletin, 133(2), 273–293.

Zadbood, A., Chen, J., Leong, Y. C., Norman, K. A., & Hasson, U. (2017). How We Transmit Memories to Other Brains: Constructing Shared Neural Representations Via Communication. Cerebral Cortex, 27(10), 4988–5000.

Zeki, S. (2015). Area V5—a microcosm of the visual brain. Frontiers in Integrative Neuroscience, 9, 21.

