## Supplementary Materials for "Broad Brain Networks Support Curiosity-Motivated Incidental Learning Of Naturalistic Dynamic Stimuli With And Without Monetary Incentives"

### Supplementary Tables

**Table S1**

*Welch's Two-Sample t-Test For Between-group Differences In The TMI Scales*

| | Intrinsic motivation | Task engagement | Interest | Boredom | Effort | Pressure |
|---|---|---|---|---|---|---|
| | | | Behavioural study | | | |
| Control group | 5.32 (1.88) [1.00; 7.00] | 5.28 (1.39) [2.33; 7.00] | 5.54 (1.86) [1.00; 7.00] | 2.98 (2.14) [1.00; 7.00] | 5.78 (1.20) [2.00; 7.00] | 1.89 (1.06) [1.00; 5.25] |
| Incentive group | 5.55 (1.46) [1.67; 7.00] | 5.46 (1.02) [3.00; 7.00] | 5.82 (1.42) [2.67; 7.00] | 2.62 (1.63) [1.00; 6.00] | 5.62 (1.33) [2.00; 7.00] | 1.72 (1.01) [1.00; 5.00] |
| Group comparison | t(69.774) = -0.603, p = 0.548 | t(67.698) = -0.648, p = 0.519 | t(69.181) = -0.732, p = 0.467 | t(69.156) = 0.824, p = 0.413 | t(74.599) = 0.567, p = 0.572 | t(74.619) = 0.749, p = 0.456 |
| Cohen's d | 0.14 [-0.31; 0.59] | 0.15 [-0.30; 0.60] | 0.17 [-0.28; 0.62] | -0.19 [-0.64; 0.26] | -0.13 [-0.58; 0.32] | -0.17 [-0.62; 0.28] |
| | | | Replication | | | |
| Control group | 5.58 (1.20) [2.67; 7.00] | 5.40 (1.01) [3.33; 7.00] | 5.87 (1.00) [3.67; 7.00] | 2.69 (1.48) [1.00; 5.67] | 5.62 (1.03) [3.00; 7.00] | 2.53 (1.22) [1.00; 5.20] |
| Incentive group | 5.80 (1.30) [2.67; 7.00] | 5.61 (0.98) [3.33; 7.00] | 5.95 (1.25) [2.33; 7.00] | 2.63 (1.46) [1.00; 5.67] | 5.78 (0.99) [3.40; 7.00] | 3.02 (1.27) [1.00; 5.60] |
| Group comparison | t(74.589) = -0.758, p = 0.451 | t(75.944) = -0.911, p = 0.365 | t(70.803) = -0.314, p = 0.754 | t(75.855) = 0.180, p = 0.857 | t(75.979) = -0.738, p = 0.463 | t(75.412) = -1.720, p = 0.090 |
| Cohen's d | 0.17 [-0.27; 0.62] | 0.21 [-0.24; 0.65] | 0.07 [-0.37; 0.52] | -0.04 [-0.49; 0.40] | 0.17 [-0.28; 0.61] | 0.40 [-0.06; 0.85] |

| | | | | | | |
|---|---|---|---|---|---|---|
| | | | fMRI study | | | |
| Control group | 5.24 (1.16) [2.33; 7.00] | 5.35 (0.88) [4.00; 6.67] | 5.32 (1.12) [3.00; 7.00] | 2.87 (1.10) [1.00; 5.00] | 4.93 (0.93) [3.00; 6.40] | 2.89 (1.41) [1.00; 5.40] |
| Incentive group | 5.24 (1.05) [3.00; 6.67] | 5.33 (0.71) [4.33; 6.67] | 5.75 (0.87) [3.67; 7.00] | 3.24 (1.30) [1.00; 6.00] | 5.40 (1.07) [3.60; 7.00] | 2.82 (1.13) [1.20; 4.40] |
| Group comparison | t(47.470) = 0.000, p = 1.000 | t(45.947) = 0.059, p = 0.953 | t(45.296) = -1.502, p = 0.140 | t(46.682) = -1.097, p = 0.278 | t(47.092) = -1.661, p = 0.103 | t(45.866) = 0.199, p = 0.843 |
| Cohen's d | 0.00 [-0.55; 0.55] | -0.02 [-0.57; 0.54] | 0.43 [-0.14; 1.00] | 0.32 [-0.25; 0.88] | 0.48 [-0.10; 1.05] | -0.06 [-0.61; 0.50] |

*Note*. For each TMI scale, the table shows mean (standard deviation) [minimum; maximum] separately for each group and data collection. To test for differences, Welch Two Sample t-tests were used and the table reports *t* statistics and *p* values. Effects were quantified using Cohen's d [95%-confidence interval]. TMI = Task Motivation Inventory.

**Table S2**

*Results Of LME Model Predicting Curiosity Ratings By Group*

| | Estimate | *SE* | *t* value |
|---|---|---|---|
| | Behavioural study | | |
| Intercept | 4.408 | 0.157 | 28.098 |
| Incentive effect | 0.058 | 0.148 | 0.393 |
| | Replication | | |
| Intercept | 4.793 | 0.136 | 35.164 |
| Incentive effect | 0.152 | 0.118 | 1.284 |
| | fMRI study | | |
| Intercept | 4.407 | 0.14 | 31.502 |
| Incentive effect | -0.056 | 0.119 | -0.465 |

*Note*. The LME model specified random intercepts for subject and stimulus ID. Incentive effect was effect-coded (control = -1, incentive = 1). *p* values are omitted as the *lme4* package does not compute them by default. LME = Linear mixed effects. *SE* = standard error.

**Table S3**

*Integrated Results Of gLME Models Predicting Recognition Memory With Gradual Increase In Confidence Using Curiosity, Monetary Incentive, And Their Interaction*

| Effect | b (*SE*) | OR [95%-CI] | *z* value | *p* value |
|---|---|---|---|---|
| | | Recog [Conf > 0] | | |
| Curiosity | 0.023 (0.023) | 1.02 [0.98; 1.07] | 0.988 | 0.323 |
| Monetary incentive | 0.084 (0.050) | 1.09 [0.99; 1.20] | 1.676 | 0.094 |
| Interaction | -0.002 (0.022) | 1.00 [0.96; 1.04] | -0.070 | 0.944 |
| | | Recog [Conf > 1] | | |
| Curiosity | 0.038 (0.022) | 1.04 [1.00; 1.08] | 1.747 | 0.081 |
| Monetary incentive | 0.117 (0.054) | 1.12 [1.01; 1.25] | 2.161 | 0.031 |
| Interaction | -0.015 (0.021) | 0.98 [0.94; 1.03] | -0.725 | 0.468 |
| | | Recog [Conf > 2] | | |
| Curiosity | 0.043 (0.021) | 1.04 [1.00; 1.09] | 2.021 | 0.043 |
| Monetary incentive | 0.156 (0.055) | 1.17 [1.05; 1.30] | 2.811 | 0.005 |
| Interaction | -0.026 (0.020) | 0.97 [0.94; 1.01] | -1.300 | 0.194 |
| | | Recog [Conf > 3] | | |
| Curiosity | 0.084 (0.022) | 1.09 [1.04; 1.14] | 3.766 | < 0.001 |
| Monetary incentive | 0.155 (0.067) | 1.17 [1.03; 1.33] | 2.336 | 0.019 |
| Interaction | -0.010 (0.021) | 0.99 [0.95; 1.03] | -0.479 | 0.632 |
| | | Recog [Conf > 4] | | |
| Curiosity | 0.122 (0.027) | 1.13 [1.07; 1.19] | 4.496 | < 0.001 |
| Monetary incentive | 0.096 (0.074) | 1.10 [0.95; 1.27] | 1.294 | 0.196 |
| Interaction | -0.014 (0.025) | 0.99 [0.94; 1.04] | -0.549 | 0.583 |
| | | Recog [Conf > 5] | | |
| Curiosity | 0.111 (0.033) | 1.12 [1.05; 1.19] | 3.388 | 0.001 |

| | | | | |
|---|---|---|---|---|
| Monetary incentive | 0.010 (0.086) | 1.01 [0.85; 1.20] | 0.118 | 0.906 |
| Interaction | -0.057 (0.028) | 0.94 [0.89; 1.00] | -2.056 | 0.040 |

*Note*. The same model specifying the full random effects structure was run for each recognition memory threshold. Please note that Recog [Conf > 0] and Recog [Conf > 3] are equal to Recognition and High confidence recognition in Table 2, respectively. gLME = Generalised Linear Mixed Effects. b = unstandardised regression coefficient. *SE* = standard error. OR = Odds Ratio, CI = confidence interval. Recog = recognition. Conf > [0:5] = confidence above given threshold.

**Table S4**

*Results Of gLME Models For Each Data Collection Specifying The Full Random Effects Structure*

| | Behavioural study | | | Replication | | | fMRI study | | |
|---|---|---|---|---|---|---|---|---|---|
| Fixed Effect | b (*SE*) | OR | 95%-CI | b (*SE*) | OR | 95%-CI | b (*SE*) | OR | 95%-CI |
| | | | | Recog [Conf > 0] | | | | | |
| Curiosity | 0.026 (0.037) | 1.03 | [0.96; 1.10] | 0.024 (0.039) | 1.02 | [0.95; 1.11] | 0.016 (0.044) | 1.02 | [0.93; 1.11] |
| Monetary incentive | 0.158 (0.094) | 1.17 | [0.98; 1.41] | 0.229 (0.083) | 1.26 | [1.07; 1.48] | -0.130 (0.085) | 0.88 | [0.74; 1.04] |
| Interaction | 0.027 (0.036) | 1.03 | [0.96; 1.10] | -0.048 (0.038) | 0.95 | [0.89; 1.03] | 0.018 (0.043) | 1.02 | [0.94; 1.11] |
| | | | | Recog [Conf > 1] | | | | | |
| Curiosity | 0.042 (0.036) | 1.04 | [0.97; 1.12] | 0.047 (0.038) | 1.05 | [0.97; 1.13] | * 0.024 (0.040) | 1.02 | [0.95; 1.11] |
| Monetary incentive | 0.161 (0.108) | 1.18 | [0.95; 1.45] | 0.257 (0.084) | 1.29 | [1.10; 1.52] | * -0.096 (0.094) | 0.91 | [0.75; 1.09] |
| Interaction | 0.006 (0.035) | 1.01 | [0.94; 1.08] | -0.046 (0.036) | 0.95 | [0.89; 1.02] | * -0.005 (0.039) | 1.00 | [0.92; 1.07] |
| | | | | Recog [Conf > 2] | | | | | |
| Curiosity | 0.013 (0.037) | 1.01 | [0.94; 1.09] | * 0.075 (0.034) | 1.08 | [1.01; 1.15] | * 0.034 (0.039) | 1.03 | [0.96; 1.12] |
| Monetary incentive | 0.247 (0.118) | 1.28 | [1.02; 1.62] | * 0.265 (0.084) | 1.30 | [1.11; 1.54] | * -0.044 (0.095) | 0.96 | [0.79; 1.15] |
| Interaction | 0.009 (0.036) | 1.01 | [0.94; 1.08] | * -0.058 (0.033) | 0.94 | [0.89; 1.01] | * -0.024 (0.038) | 0.98 | [0.91; 1.05] |
| | | | | Recog [Conf > 3] | | | | | |
| Curiosity | * 0.048 (0.038) | 1.05 | [0.97; 1.13] | 0.136 (0.037) | 1.15 | [1.07; 1.23] | * 0.061 (0.040) | 1.06 | [0.98; 1.15] |

| | | | | | | | | | |
|---|---|---|---|---|---|---|---|---|---|
| Monetary incentive | * 0.298 (0.128) | 1.35 | [1.05; 1.73] | 0.245 (0.106) | 1.28 | [1.04; 1.57] | * -0.062 (0.114) | 0.94 | [0.75; 1.18] |
| Interaction | * 0.006 (0.036) | 1.01 | [0.94; 1.08] | -0.021 (0.034) | 0.98 | [0.92; 1.05] | * -0.015 (0.038) | 0.99 | [0.91; 1.06] |
| | | | | Recog [Conf > 4] | | | | | |
| Curiosity | 0.088 (0.043) | 1.09 | [1.00; 1.19] | 0.138 (0.044) | 1.15 | [1.05; 1.25] | 0.152 (0.057) | 1.16 | [1.04; 1.30] |
| Monetary incentive | 0.280 (0.140) | 1.32 | [1.01; 1.74] | 0.192 (0.123) | 1.21 | [0.95; 1.54] | -0.148 (0.124) | 0.86 | [0.68; 1.10] |
| Interaction | 0.004 (0.039) | 1.00 | [0.93; 1.08] | -0.038 (0.040) | 0.96 | [0.89; 1.04] | -0.002 (0.053) | 1.00 | [0.90; 1.11] |
| | | | | Recog [Conf > 5] | | | | | |
| Curiosity | * 0.118 (0.051) | 1.13 | [1.02; 1.24] | 0.071 (0.055) | 1.07 | [0.96; 1.19] | 0.158 (0.068) | 1.17 | [1.03; 1.34] |
| Monetary incentive | * 0.255 (0.149) | 1.29 | [0.96; 1.73] | 0.053 (0.165) | 1.05 | [0.76; 1.46] | -0.227 (0.137) | 0.80 | [0.61; 1.04] |
| Interaction | * -0.003 (0.042) | 1.00 | [0.92; 1.08] | -0.148 (0.047) | 0.86 | [0.79; 0.95] | -0.019 (0.060) | 0.98 | [0.87; 1.10] |
| | | | | Recall | | | | | |
| Curiosity | 0.033 (0.052) | 1.03 | [0.93; 1.14] | * 0.107 (0.038) | 1.11 | [1.03; 1.20] | * 0.135 (0.046) | 1.14 | [1.05; 1.25] |
| Monetary incentive | 0.096 (0.171) | 1.10 | [0.79; 1.54] | * 0.228 (0.105) | 1.26 | [1.02; 1.54] | * -0.043 (0.134) | 0.96 | [0.74; 1.25] |
| Interaction | -0.015 (0.048) | 0.99 | [0.90; 1.08] | * -0.063 (0.036) | 0.94 | [0.88; 1.01] | * 0.019 (0.043) | 1.02 | [0.94; 1.11] |

*Note.* For each data collection and memory measurement, the same gLME model was run. Full random effects structure indicates random intercepts for participant and stimulus, as well as random slopes for the curiosity effect. If the model produced a singular fit error, an asterisk was added before reporting the coefficients. gLME = generalised linear mixed effects. b = unstandardised regression coefficient. *SE* = standard error. OR = Odds Ratio. CI = confidence interval. Recog = recognition. Conf > [0:5] = confidence above given threshold.

**Table S5**

*Integrated Results Of gLME Models With Reduced Random Effects Structure Predicting Memory Encoding Using Monetary Incentive, Curiosity, And Their Interaction*

| Effect | b (*SE*) | OR [95%-CI] | *z* value | *p* value |
|---|---|---|---|---|
| | | Recog [Conf > 0] | | |
| Curiosity | 0.024 (0.021) | 1.02 [0.98; 1.07] | 1.152 | 0.249 |
| Monetary incentive | 0.083 (0.050) | 1.09 [0.99; 1.20] | 1.675 | 0.094 |
| Interaction | -0.005 (0.020) | 0.99 [0.96; 1.03] | -0.264 | 0.792 |
| | | Recog [Conf > 1] | | |
| Curiosity | 0.038 (0.021) | 1.04 [1.00; 1.08] | 1.838 | 0.066 |
| Monetary incentive | 0.116 (0.054) | 1.12 [1.01; 1.25] | 2.156 | 0.031 |
| Interaction | -0.017 (0.020) | 0.98 [0.95; 1.02] | -0.849 | 0.396 |
| | | Recog [Conf > 2] | | |
| Curiosity | 0.042 (0.021) | 1.04 [1.00; 1.09] | 2.041 | 0.041 |
| Monetary incentive | 0.155 (0.055) | 1.17 [1.05; 1.30] | 2.803 | 0.005 |
| Interaction | -0.026 (0.020) | 0.97 [0.94; 1.01] | -1.295 | 0.195 |
| | | Recog [Conf > 3] | | |
| Curiosity | 0.085 (0.021) | 1.09 [1.04; 1.13] | 4.002 | < 0.001 |
| Monetary incentive | 0.155 (0.066) | 1.17 [1.03; 1.33] | 2.336 | 0.019 |
| Interaction | -0.011 (0.020) | 0.99 [0.95; 1.03] | -0.541 | 0.589 |
| | | Recog [Conf > 4] | | |
| Curiosity | 0.111 (0.022) | 1.12 [1.07; 1.17] | 4.968 | < 0.001 |
| Monetary incentive | 0.092 (0.073) | 1.10 [0.95; 1.27] | 1.251 | 0.211 |
| Interaction | -0.015 (0.021) | 0.99 [0.94; 1.03] | -0.702 | 0.483 |
| | | Recog [Conf > 5] | | |
| Curiosity | 0.115 (0.026) | 1.12 [1.07; 1.18] | 4.456 | < 0.001 |
| Monetary incentive | 0.005 (0.085) | 1.00 [0.85; 1.19] | 0.054 | 0.957 |
| Interaction | -0.060 (0.025) | 0.94 [0.90; 0.99] | -2.448 | 0.014 |

| | | | | |
|---|---|---|---|---|
| | | Recall | | |
| Curiosity | 0.102 (0.024) | 1.11 [1.06; 1.16] | 4.335 | < 0.001 |
| Monetary incentive | 0.119 (0.074) | 1.13 [0.97; 1.30] | 1.606 | 0.108 |
| Interaction | -0.024 (0.023) | 0.98 [0.93; 1.02] | -1.063 | 0.288 |

*Note*. Reduced random effects structure specifies random intercepts for participant and stimulus, but omits random slopes for the curiosity effect. Recog [Conf > 0] and Recog [Conf > 3] are the same as Recognition and High confidence recognition in Table 2, respectively. gLME = Generalised Linear Mixed Effects. b = unstandardised regression coefficient. *SE* = standard error. OR = Odds Ratio, CI = confidence interval. Recog = recognition. Conf > [0:5] = confidence above given threshold.

**Table S6**

*Predicting Integrated Effects Using The Gradual Confidence Cut-off*

| | Full gLME model | | | | | Reduced gLME model | | | | |
|---|---|---|---|---|---|---|---|---|---|---|
| | B (*SE*) | 95%-CI | *t* value | *p* value | Fit | B (*SE*) | 95%-CI | *t* value | *p* value | Fit |
| | | | | | Curiosity | | | | | |
| Intercept | 0.018 (0.011) | [-0.012; 0.047] | 1.652 | 0.174 | | 0.018 (0.008) | [-0.004; 0.041] | 2.230 | 0.090 | |
| Slope | 0.021 (0.004) | [ 0.011; 0.031] | 5.920 | 0.004 | | 0.020 (0.003) | [ 0.013; 0.028] | 7.562 | 0.002 | |
| | | | | | $R2 = 0.898$ | | | | | $R2 = 0.935$ |
| | | | | | Monetary incentive | | | | | |
| Intercept | 0.134 (0.040) | [ 0.023; 0.244] | 3.361 | 0.028 | | 0.134 (0.041) | [ 0.021; 0.247] | 3.297 | 0.030 | |
| Slope | -0.012 (0.013) | [-0.049; 0.024] | -0.937 | 0.402 | | -0.013 (0.013) | [-0.051; 0.024] | -0.990 | 0.378 | |
| | | | | | $R2 = 0.180$ | | | | | $R2 = 0.197$ |
| | | | | | Interaction | | | | | |
| Intercept | -0.002 (0.011) | [-0.034; 0.029] | -0.213 | 0.842 | | -0.004 (0.012) | [-0.037; 0.028] | -0.362 | 0.735 | |
| Slope | -0.007 (0.004) | [-0.018; 0.003] | -1.954 | 0.122 | | -0.007 (0.004) | [-0.018; 0.003] | -1.895 | 0.131 | |
| | | | | | $R2 = 0.488$ | | | | | $R2 = 0.473$ |

*Note*. For each fixed effect, a linear model was used to predict the integrated effect using the confidence cut-off value. The integrated results are based on the full and reduced gLME model, respectively. The confidence cut-off value was defined from 0 to 5 so that the intercept can be interpreted. gLME = Generalised Linear Mixed Effects. *SE* = standard error. CI = confidence interval.

**Table S7**

*Results Of gLME Models For Each Data Collection Specifying The Reduced Random Effects Structure*

| | Behavioural study | | | Replication | | | fMRI study | | |
|---|---|---|---|---|---|---|---|---|---|
| Fixed Effect | b (*SE*) | OR | 95%-CI | b (*SE*) | OR | 95%-CI | b (*SE*) | OR | 95%-CI |
| | | | | Recog [Conf > 0] | | | | | |
| Curiosity | 0.026 (0.035) | 1.03 | [0.96; 1.10] | 0.023 (0.035) | 1.02 | [0.96; 1.10] | 0.023 (0.039) | 1.02 | [0.95; 1.10] |
| Monetary incentive | 0.158 (0.093) | 1.17 | [0.98; 1.41] | 0.227 (0.082) | 1.25 | [1.07; 1.47] | -0.129 (0.084) | 0.88 | [0.75; 1.04] |
| Interaction | 0.024 (0.034) | 1.02 | [0.96; 1.10] | -0.052 (0.034) | 0.95 | [0.89; 1.01] | 0.018 (0.037) | 1.02 | [0.95; 1.10] |
| | | | | Recog [Conf > 1] | | | | | |
| Curiosity | 0.041 (0.035) | 1.04 | [0.97; 1.12] | 0.045 (0.035) | 1.05 | [0.98; 1.12] | 0.026 (0.038) | 1.03 | [0.95; 1.11] |
| Monetary incentive | 0.161 (0.108) | 1.18 | [0.95; 1.45] | 0.255 (0.083) | 1.29 | [1.10; 1.52] | -0.096 (0.094) | 0.91 | [0.76; 1.09] |
| Interaction | 0.005 (0.035) | 1.01 | [0.94; 1.08] | -0.049 (0.033) | 0.95 | [0.89; 1.02] | -0.002 (0.037) | 1.00 | [0.93; 1.07] |
| | | | | Recog [Conf > 2] | | | | | |
| Curiosity | 0.015 (0.036) | 1.02 | [0.95; 1.09] | 0.075 (0.034) | 1.08 | [1.01; 1.15] | 0.032 (0.038) | 1.03 | [0.96; 1.11] |
| Monetary incentive | 0.247 (0.118) | 1.28 | [1.02; 1.61] | 0.265 (0.084) | 1.30 | [1.11; 1.54] | -0.044 (0.094) | 0.96 | [0.80; 1.15] |
| Interaction | 0.008 (0.035) | 1.01 | [0.94; 1.08] | -0.058 (0.032) | 0.94 | [0.89; 1.01] | -0.022 (0.037) | 0.98 | [0.91; 1.05] |
| | | | | Recog [Conf > 3] | | | | | |

| | | | | | | | | | |
|---|---|---|---|---|---|---|---|---|---|
| Curiosity | 0.058 (0.037) | 1.06 | [0.99; 1.14] | 0.132 (0.035) | 1.14 | [1.07; 1.22] | 0.055 (0.039) | 1.06 | [0.98; 1.14] |
| Monetary incentive | 0.298 (0.128) | 1.35 | [1.05; 1.73] | 0.244 (0.106) | 1.28 | [1.04; 1.57] | -0.062 (0.114) | 0.94 | [0.75; 1.18] |
| Interaction | 0.004 (0.036) | 1.00 | [0.94; 1.08] | -0.021 (0.033) | 0.98 | [0.92; 1.04] | -0.015 (0.037) | 0.99 | [0.92; 1.06] |
| | | | | Recog [Conf > 4] | | | | | |
| Curiosity | 0.089 (0.038) | 1.09 | [1.01; 1.18] | 0.139 (0.036) | 1.15 | [1.07; 1.23] | 0.101 (0.043) | 1.11 | [1.02; 1.20] |
| Monetary incentive | 0.279 (0.140) | 1.32 | [1.01; 1.74] | 0.193 (0.123) | 1.21 | [0.95; 1.54] | -0.148 (0.121) | 0.86 | [0.68; 1.09] |
| Interaction | 0.002 (0.037) | 1.00 | [0.93; 1.08] | -0.038 (0.034) | 0.96 | [0.90; 1.03] | -0.003 (0.040) | 1.00 | [0.92; 1.08] |
| | | | | Recog [Conf > 5] | | | | | |
| Curiosity | 0.107 (0.043) | 1.11 | [1.02; 1.21] | 0.130 (0.042) | 1.14 | [1.05; 1.24] | 0.102 (0.050) | 1.11 | [1.00; 1.22] |
| Monetary incentive | 0.256 (0.149) | 1.29 | [0.96; 1.73] | 0.064 (0.165) | 1.07 | [0.77; 1.47] | -0.234 (0.133) | 0.79 | [0.61; 1.03] |
| Interaction | -0.003 (0.042) | 1.00 | [0.92; 1.08] | -0.141 (0.040) | 0.87 | [0.80; 0.94] | -0.021 (0.048) | 0.98 | [0.89; 1.08] |
| | | | | Recall | | | | | |
| Curiosity | 0.068 (0.041) | 1.07 | [0.99; 1.16] | 0.109 (0.038) | 1.12 | [1.04; 1.20] | 0.134 (0.045) | 1.14 | [1.05; 1.25] |
| Monetary incentive | 0.096 (0.169) | 1.10 | [0.79; 1.53] | 0.229 (0.105) | 1.26 | [1.02; 1.55] | -0.043 (0.134) | 0.96 | [0.74; 1.25] |
| Interaction | -0.015 (0.040) | 0.99 | [0.91; 1.06] | -0.062 (0.036) | 0.94 | [0.88; 1.01] | 0.020 (0.043) | 1.02 | [0.94; 1.11] |

*Note.* For each data collection and memory measurement, the same gLME model was run. Reduced random effects structure specifies random intercepts for participant and stimulus, but omits random slopes for the curiosity effect. gLME = generalised linear mixed effects. b =

unstandardised regression coefficient. *SE* = standard error. OR = Odds Ratio. CI = confidence interval. Recog = recognition. Conf > [0:5] = confidence above given threshold.

**Table S8**

*ROI And Whole Brain Results Of ISC And Incentive Effects Therein*

| Cluster | Macro Label | Direction | Cluster Size | Maximum Intensity | | | |
|---|---|---|---|---|---|---|---|
| | | | | *t* | RL | AP | IS |
| | | | Average ISC | | | | |
| Regions-of-Interest Approach | | | | | | | |
| 1 | R aHPC | positive | 62 | 7.1708 | -22.5 | 1.5 | -19.5 |
| 2 | L aHPC | positive | 45 | 5.8873 | 31.5 | 7.5 | -19.5 |
| 1 | L VTA/SN | positive | 30 | 6.927 | 7.5 | 13.5 | -10.5 |
| 2 | R VTA/SN | positive | 29 | 4.9475 | -4.5 | 16.5 | -16.5 |
| 1 | R NAcc | positive | 31 | 6.8451 | -16.5 | -7.5 | -10.5 |
| 2 | L Nacc | positive | 27 | 6.0561 | 13.5 | -10.5 | -10.5 |
| 1 | R CN | positive | 273 | 13.686 | -13.5 | -22.5 | 4.5 |
| 2 | L CN | positive | 270 | 12.944 | 19.5 | -16.5 | 10.5 |
| Whole Brain Analysis | | | | | | | |
| 1 | R Middle Frontal Gyrus<br>L Middle Frontal Gyrus<br>L Middle Temporal Gyrus<br>R Middle Temporal Gyrus<br>R Superior Frontal Gyrus<br>L Precuneus<br>L Precentral Gyrus<br>L Middle Occipital Gyrus<br>R Precuneus8<br>L Postcentral Gyrus<br>L Superior Frontal Gyrus<br>L Superior Medial Gyrus<br>R Postcentral Gyrus<br>R Precentral Gyrus | positive | 45359 | 18.783 | -46.5 | 70.5 | -7.5 |

R Inferior Temporal
Gyrus
R Superior Temporal
Gyrus
L Inferior Parietal
Lobule
L Cerebellum (Crus 1)
L Calcarine Gyrus
R Cerebellum (Crus 1)
R Lingual Gyrus
L Inferior Frontal Gyrus
(p. Triangularis)
L Lingual Gyrus
L SMA
R Middle Occipital
Gyrus
L Inferior Temporal
Gyrus
R Fusiform Gyrus
R Superior Parietal
Lobule
L Superior Parietal
Lobule
R SMA
R SupraMarginal Gyrus
L Middle Cingulate
Cortex
L Cerebellum (Crus 2)3
R Superior Medial
Gyrus
L Superior Temporal
Gyrus
R Middle Cingulate
Cortex
R Angular Gyrus
R Cerebellum (Crus 2)
R Cerebellum (VI)
R Calcarine Gyrus
L Fusiform Gyrus
R Inferior Frontal Gyrus
(p. Triangularis)
R Insula Lobe
L Cuneus
L Cerebellum (VI)
R Cerebellum (VIII)
R Inferior Parietal
Lobule
R Cuneus
L Inferior Frontal Gyrus
(p. Orbitalis)

L Insula Lobe
R Inferior Frontal Gyrus
(p. Orbitalis)
L Anterior Cingulate
Cortex
R Superior Occipital
Gyrus
L Superior Occipital
Gyrus
R Inferior Frontal Gyrus
(p. Opercularis)
R Rolandic Operculum
L SupraMarginal Gyrus
L Angular Gyrus
R Inferior Occipital
Gyrus
L Cerebellum (VIII)
L Thalamus
R Anterior Cingulate
Cortex
L Rolandic Operculum
L Inferior Occipital
Gyrus
R Middle Orbital Gyrus
L Putamen
L Inferior Frontal Gyrus
(p. Opercularis)
R Putamen
L Paracentral Lobule
R Caudate Nucleus
R Temporal Pole
L Caudate Nucleus
R Thalamus
L Middle Orbital Gyrus
L Mid Orbital Gyrus
R Medial Temporal
Pole
R Mid Orbital Gyrus
L Temporal Pole
R Paracentral Lobule
L Hippocampus
L Cerebellum (VII)
R Cerebellum (IV-V)
R Superior Orbital
Gyrus
R Cerebellum (VII)
R Hippocampus
L Posterior Cingulate
Cortex
Cerebellar Vermis (6)

| | |
|---|---|
| | L Superior Orbital Gyrus |
| | R ParaHippocampal Gyrus |
| | Cerebellar Vermis (4/5) |
| | R Heschls Gyrus |
| | L Cerebellum (IX) |
| | L Heschls Gyrus9 |
| | L Medial Temporal Pole |
| | R Cerebellum (IX) |
| | L Cerebellum (IV-V) |
| | L ParaHippocampal Gyrus |
| | R Pallidum |
| | R Amygdala |
| | R Posterior Cingulate Cortex |
| | L Amygdala |
| | L Olfactory cortex |
| | Cerebellar Vermis (7) |
| | L Rectal Gyrus |
| | L Pallidum |
| | R Rectal Gyrus |
| | R Olfactory cortex |
| | R Cerebellum (X) |
| | Cerebellar Vermis (8) |
| | Cerebellar Vermis (3) |
| | L Cerebellum (X) |

ISC Incentive Effects

Regions-of-Interest Approach

*no clusters survive thresholding*

Whole Brain Analysis

| | | | | | | | |
|---|---|---|---|---|---|---|---|
| 1 | L Middle Occipital Gyrus | I > C | 85 | -8.0421 | 43.5 | 91.5 | -1.5 |
| 2 | R Postcentral Gyrus | I > C | 53 | -11.744 | -37.5 | 31.5 | 49.5 |
| | Superior Parietal Lobule | | | | | | |
| 3 | R Superior Occipital Gyrus | I > C | 38 | -6.101 | -25.5 | 70.5 | 46.5 |
| | L Middle Occipital Gyrus | | | | | | |
| 4 | L Inferior Occipital Gyrus | I < C | 28 | 6.0377 | 43.5 | 67.5 | -1.5 |

*Note.* Results are thresholded at $q < 0.05$ and $k = 5$ for the regions-of-interest approach and at $p < 0.001$, cluster-extent corrected at $k = 20$ (equivalent to per-cluster $\alpha = 0.05$) for whole brain analysis. The table corresponds to Figure 4 and S6. $t$ = $t$ value, RL = right to left, AP = anterior to posterior, IS = inferior to superior. RAI orientation is applied where negative values indicate right, anterior, inferior.

**Table S9**

*ROI And Whole Brain Results Of IS-RSA For Each Behavioural Effect Of Interest*

| Cluster | Macro Label | Direction | Cluster Size | Maximum Intensity | | | |
|---|---|---|---|---|---|---|---|
| | | | | *t* | RL | AP | IS |
| | | | Curiosity | | | | |
| Regions-of-Interest Approach | | | | | | | |
| | | *no clusters survive thresholding* | | | | | |
| Whole Brain Analysis | | | | | | | |
| 1 | L Inferior Occipital Gyrus<br>L Lingual Gyrus<br>L Calcarine Gyrus | positive | 63 | 6.2884 | 13.5 | 106.5 | -4.5 |
| 2 | R Inferior Frontal Gyrus (p. Opercularis) | positive | 34 | 4.9267 | -52.5 | -16.5 | 25.5 |
| 3 | R SMA | positive | 33 | 4.9359 | -1.5 | -13.5 | 61.5 |
| 4 | L Postcentral Gyrus | positive | 29 | 4.9618 | 25.5 | 31.5 | 73.5 |
| 5 | L Precuneus | positive | 23 | 4.2003 | 1.5 | 64.5 | 61.5 |
| 6 | R Insula Lobe | positive | 20 | 5.2905 | -37.5 | -22.5 | 1.5 |
| 7 | R SupraMarginal Gyrus | positive | 20 | 4.202 | -67.5 | 25.5 | 37.5 |
| | | | Memory | | | | |
| Regions-of-Interest Approach | | | | | | | |
| 1 | R CN | positive | 30 | 4.1269 | -13.5 | -19.5 | 4.5 |
| 2 | L CN | positive | 20 | 4.253 | 19.5 | -22.5 | 4.5 |
| Whole Brain Analysis | | | | | | | |
| 1 | R Inferior Occipital Gyrus<br>Middle Occipital Gyrus<br>Lingual Gyrus<br>Middle Temporal Gyrus | positive | 455 | 7.9417 | -34.5 | 85.5 | -10.5 |

| | | | | | | | |
|---|---|---|---|---|---|---|---|
| 2 | R Calcarine Gyrus<br>L Lingual Gyrus<br>R Lingual Gyrus<br>L Cuneus<br>R Cuneus | positive | 320 | 5.5756 | -7.5 | 64.5 | 4.5 |
| 3 | L Middle Occipital Gyrus<br>L Inferior Occipital Gyrus<br>L Fusiform Gyrus | positive | 270 | 7.7376 | 34.5 | 64.5 | -13.5 |
| 4 | L Calcarine Gyrus<br>L Cuneus<br>R Cuneus<br>R Calcarine Gyrus | positive | 233 | 7.1555 | -4.5 | 94.5 | 10.5 |
| 5 | L Middle Frontal Gyrus | positive | 81 | 5.1844 | 34.5 | -52.5 | 7.5 |
| 6 | R Precuneus<br>L Precuneus | positive | 75 | 5.5491 | -1.5 | 70.5 | 55.5 |
| 7 | R Middle Frontal Gyrus | positive | 59 | 5.5209 | -49.5 | -46.5 | 13.5 |
| 8 | R Middle Frontal Gyrus | positive | 55 | 4.973 | -31.5 | -43.5 | 31.5 |
| 9 | R Angular Gyrus | positive | 50 | 5.3212 | -43.5 | 70.5 | 43.5 |
| 10 | R Postcentral Gyrus | positive | 45 | 6.2064 | -31.5 | 34.5 | 43.5 |
| 11 | R Middle Frontal Gyrus | positive | 42 | 5.2976 | -31.5 | -19.5 | 58.5 |
| 12 | R SMA | positive | 42 | 5.2509 | -7.5 | -19.5 | 49.5 |
| 13 | L Cerebellum (Crus 1)<br>L Cerebellum (Crus 2) | positive | 34 | 4.5986 | 16.5 | 82.5 | -28.5 |
| 14 | L Cerebellum (Crus 2)<br>L Cerebellum (Crus 1) | positive | 33 | 5.0131 | 31.5 | 76.5 | -37.5 |
| 15 | L Inferior Temporal Gyrus | positive | 33 | 5.3724 | 55.5 | 58.5 | -7.5 |
| 16 | R Insula Lobe | positive | 33 | 5.2887 | -31.5 | -25.5 | -4.5 |
| 17 | L Middle Occipital Gyrus | positive | 32 | 5.7279 | 46.5 | 85.5 | 4.5 |
| 18 | R Middle Frontal Gyrus | positive | 29 | 4.8888 | -43.5 | -16.5 | 43.5 |
| 19 | L Cerebellum (Crus 1) | positive | 22 | 5.3657 | 43.5 | 73.5 | -25.5 |
| 20 | L Middle Frontal Gyrus | positive | 22 | 5.0285 | 25.5 | -37.5 | 37.5 |
| 21 | L Inferior Parietal Lobule | positive | 20 | 4.7087 | 55.5 | 40.5 | 52.5 |

| | | | | | | | |
|---|---|---|---|---|---|---|---|
| Regions-of-Interest Approach | | | | | | | |
| 1 | R aHPC | negative | 8 | -4.456 | -31.5 | 4.5 | -19.5 |
| 2 | R VTA/SN | negative | 5 | -3.401 | -13.5 | 22.5 | -10.5 |
| 1 | R NAcc | negative | 25 | -5.5516 | -10.5 | -10.5 | -4.5 |
| 2 | L Nacc | negative | 11 | -4.4598 | 13.5 | -13.5 | -10.5 |
| 1 | R CN | negative | 188 | -8.0879 | -13.5 | -16.5 | 7.5 |
| 2 | L CN | negative | 176 | -8.9741 | 16.5 | -4.5 | 16.5 |
| Whole Brain Analysis | | | | | | | |
| 1 | R Calcarine Gyrus | positive | 42 | 6.7942 | -16.5 | 88.5 | 7.5 |
| 2 | L Middle Occipital Gyrus | positive | 33 | 8.1115 | 37.5 | 82.5 | 7.5 |
| 3 | R Postcentral Gyrus | positive | 26 | 7.1842 | -40.5 | 31.5 | 58.5 |
| 4 | L Middle Temporal Gyrus | positive | 22 | 5.8015 | 49.5 | 64.5 | 7.5 |
| 5 | R Middle Temporal Gyrus | positive | 20 | 6.9451 | -43.5 | 61.5 | 7.5 |
| 6 | L Middle Frontal Gyrus<br>R Middle Frontal Gyrus<br>L Superior Medial Gyrus<br>L Precuneus<br>R Superior Frontal Gyrus<br>L Middle Temporal Gyrus<br>R Precuneus<br>R Middle Temporal Gyrus<br>L Superior Frontal Gyrus<br>L Cerebellum (Crus 1)<br>L Precentral Gyrus<br>L Inferior Frontal Gyrus (p. Triangularis)<br>R Superior Medial Gyrus<br>R Angular Gyrus<br>R Cerebellum (Crus 1)<br>L Cerebellum (Crus 2)<br>L Inferior Parietal Lobule<br>L Calcarine Gyrus<br>L Cuneus<br>L Superior Parietal Lobule<br>R Superior Parietal Lobule<br>R Cerebellum (Crus 2)<br>L Middle Occipital Gyrus<br>R Inferior Parietal Lobule | negative | 20326 | -16.137 | -49.5 | 55.5 | 49.5 |

| | | | | | | | |
|---|---|---|---|---|---|---|---|
| | L Lingual Gyrus | | | | | | |
| | R Inferior Frontal Gyrus (p. Orbitalis) | | | | | | |
| | L Inferior Frontal Gyrus (p. Orbitalis) | | | | | | |
| | L Angular Gyrus | | | | | | |
| | R Inferior Frontal Gyrus (p. Triangularis) | | | | | | |
| | L Fusiform Gyrus | | | | | | |
| | R SupraMarginal Gyrus | | | | | | |
| | L SMA | | | | | | |
| | L Middle Cingulate Cortex | | | | | | |
| | L Inferior Temporal Gyrus | | | | | | |
| | R Middle Occipital Gyrus | | | | | | |
| | L Postcentral Gyrus | | | | | | |
| | L Anterior Cingulate Cortex | | | | | | |
| | R Cuneus | | | | | | |
| | L Inferior Occipital Gyrus | | | | | | |
| | R Postcentral Gyrus | | | | | | |
| | R Inferior Temporal Gyrus | | | | | | |
| | R Precentral Gyrus | | | | | | |
| | R Caudate Nucleus | | | | | | |
| | L Thalamus | | | | | | |
| | R Thalamus | | | | | | |
| 7 | R Putamen | negative | 208 | -8.0879 | -13.5 | -16.5 | 7.5 |
| | L Caudate Nucleus | | | | | | |
| 8 | L Putamen | negative | 183 | -8.9741 | 16.5 | -4.5 | 16.5 |
| | L Rolandic Operculum | | | | | | |
| 9 | L Insula Lobe | negative | 60 | -10.656 | 40.5 | 1.5 | 16.5 |
| | L Amygdala | | | | | | |
| 10 | L Hippocampus | negative | 57 | -6.7116 | 16.5 | 7.5 | -13.5 |
| 11 | L Putamen | negative | 42 | -6.1443 | 28.5 | 4.5 | 4.5 |
| | R Postcentral Gyrus | | | | | | |
| 12 | R Precentral Gyrus | negative | 37 | -6.0096 | -28.5 | 25.5 | 58.5 |
| | R Rolandic Operculum | | | | | | |
| 13 | R Insula Lobe | negative | 36 | -8.0841 | -43.5 | 1.5 | 16.5 |
| | R Lingual Gyrus | | | | | | |
| 14 | R Calcarine Gyrus | negative | 33 | -7.401 | -7.5 | 79.5 | 1.5 |
| 15 | L SupraMarginal Gyrus | negative | 28 | -8.1472 | 52.5 | 43.5 | 28.5 |

*Note.* Results are thresholded at $q < 0.05$ and $k = 5$ for the regions-of-interest approach and at $p < 0.001$, cluster-extent corrected at $k = 20$ (equivalent to per-cluster $\alpha = 0.05$) for whole brain analysis. The table corresponds to Figure 5 and 6. $t$ = $t$ value, RL = right to left, AP = anterior to posterior, IS = inferior to superior. RAI orientation is applied where negative values indicate right, anterior, inferior.

**Table S10**

*ROI And Whole Brain Results Of IS-RSA For The Interaction Between The Incentive Manipulation And Each Behavioural Effect Of Interest*

| Cluster | Macro Label | Direction | Cluster Size | Maximum Intensity | | | |
|---|---|---|---|---|---|---|---|
| | | | | t | RL | AP | IS |
| | | | Curiosity | | | | |
| Regions-of-Interest Approach | | | | | | | |
| | | *no clusters survive thresholding* | | | | | |
| Whole Brain Analysis | | | | | | | |
| 1 | R Inferior Occipital Gyrus<br>R Middle Occipital Gyrus | I < C | 60 | 4.9726 | -31.5 | 88.5 | -4.5 |
| 2 | L Middle Occipital Gyrus | I < C | 36 | 5.4253 | 22.5 | 97.5 | 1.5 |
| | | | Memory | | | | |
| Regions-of-Interest Approach | | | | | | | |
| | | *no clusters survive thresholding* | | | | | |
| Whole Brain Analysis | | | | | | | |
| 1 | R Calcarine Gyrus<br>L Calcarine Gyrus<br>L Cuneus<br>R Cuneus | I > C | 120 | -5.3723 | -10.5 | 82.5 | 13.5 |
| 2 | L Middle Orbital Gyrus<br>L Middle Frontal Gyrus | I < C | 76 | 5.7046 | 37.5 | -58.5 | 1.5 |
| 3 | L Middle Occipital Gyrus | I < C | 25 | 5.4122 | 34.5 | 97.5 | 4.5 |
| | | | Curiosity-Motivated Learning Enhancement | | | | |
| Regions-of-Interest Approach | | | | | | | |
| | | *no clusters survive thresholding* | | | | | |

Whole Brain Analysis

| | | | | | | | |
|---|---|---|---|---|---|---|---|
| 1 | R Middle Occipital Gyrus<br>R Inferior Occipital Gyrus<br>R Middle Temporal Gyrus<br>L Lingual Gyrus<br>R Lingual Gyrus<br>L Calcarine Gyrus<br>R Inferior Temporal Gyrus<br>R Fusiform Gyrus<br>R Calcarine Gyrus | I > C | 711 | -9.0868 | -40.5 | 79.5 | -7.5 |
| 2 | L Precuneus<br>R Precuneus<br>R Cuneus<br>L Cuneus<br>L Superior Parietal Lobule | I > C | 544 | -7.0127 | -13.5 | 67.5 | 46.5 |
| 3 | L Middle Occipital Gyrus<br>L Middle Temporal Gyrus<br>L Inferior Occipital Gyrus | I > C | 297 | -8.1443 | 31.5 | 91.5 | 16.5 |
| 4 | R Superior Parietal Lobule<br>R Inferior Parietal Lobule | I > C | 131 | -9.7247 | -25.5 | 64.5 | 64.5 |
| 5 | L Middle Frontal Gyrus<br>L Middle Orbital Gyrus | I > C | 120 | -6.1818 | 37.5 | -58.5 | 10.5 |
| 6 | R Angular Gyrus<br>R Inferior Parietal Lobule | I > C | 86 | -5.9831 | -46.5 | 58.5 | 37.5 |
| 7 | R Superior Frontal Gyrus<br>R Precentral Gyrus | I > C | 54 | -6.5473 | -25.5 | 7.5 | 55.5 |
| 8 | R Superior Frontal Gyrus | I > C | 51 | -5.7977 | -22.5 | -43.5 | 43.5 |

| | | | | | | | |
|---|---|---|---|---|---|---|---|
| 9 | R Middle Cingulate Cortex<br>R Superior Medial Gyrus<br>L Superior Medial Gyrus<br>R Anterior Cingulate Cortex | I > C | 48 | -4.9674 | -4.5 | -28.5 | 43.5 |
| 10 | L Inferior Parietal Lobule<br>L Angular Gyrus | I > C | 46 | -6.486 | 46.5 | 55.5 | 46.5 |
| 11 | R Precuneus<br>R Middle Cingulate Cortex<br>R Posterior Cingulate Cortex | I > C | 45 | -5.0203 | -4.5 | 49.5 | 34.5 |
| 12 | L Superior Parietal Lobule<br>L Superior Occipital Gyrus | I > C | 41 | -7.5829 | 22.5 | 79.5 | 46.5 |
| 13 | L SupraMarginal Gyrus<br>L Inferior Parietal Lobule | I > C | 31 | -4.7041 | 64.5 | 34.5 | 40.5 |
| 14 | R Cuneus<br>R Superior Occipital Gyrus | I > C | 29 | -6.81 | -13.5 | 94.5 | 22.5 |
| 15 | R Middle Frontal Gyrus | I > C | 28 | -6.4763 | -46.5 | -46.5 | 16.5 |
| 16 | R Posterior Cingulate Cortex<br>R Precuneus<br>L Posterior Cingulate Cortex | I > C | 27 | -4.9755 | -4.5 | 43.5 | 19.5 |
| 17 | L Inferior Parietal Lobule<br>L Superior Parietal Lobule | I > C | 27 | -6.4452 | 34.5 | 49.5 | 58.5 |
| 18 | L Precentral Gyrus<br>L Superior Frontal Gyrus | I > C | 24 | -5.4886 | 25.5 | 13.5 | 55.5 |
| 19 | L Inferior Occipital Gyrus<br>L Inferior Temporal Gyrus | I > C | 23 | -6.4189 | 43.5 | 64.5 | -4.5 |

| | | | | | | | |
|---|---|---|---|---|---|---|---|
| | L Middle Temporal Gyrus | | | | | | |
| | L Middle Occipital Gyrus | | | | | | |
| | L SupraMarginal Gyrus | | | | | | |
| 20 | L Superior Temporal Gyrus | I < C | 31 | 5.331 | 49.5 | 37.5 | 28.5 |

*Note.* Results are thresholded at $q < 0.05$ and $k = 5$ for the regions-of-interest approach and at $p < 0.001$, cluster-extent corrected at $k = 20$ (equivalent to per-cluster $\alpha = 0.05$) for whole brain analysis. The table corresponds to Figure 7. $t = t$ value, RL = right to left, AP = anterior to posterior, IS = inferior to superior. RAI orientation is applied where negative values indicate right, anterior, inferior. I = Incentive group, C = Control group.

**Table S11**

*Results Of LME-CRE Models Predicting aHPC-VTA/SN-ISFC*

| | Estimate | *SE* | *t* value |
|---|---|---|---|
| | Incentive Effect | | |
| Intercept | 0.01 | 0.003 | 3.638 |
| Incentive effect | -0.002 | 0.003 | -0.962 |
| | Main Effects | | |
| Intercept | 0.01 | 0.003 | 3.757 |
| Incentive effect | -0.002 | 0.003 | -0.803 |
| Curiosity effect | 0.013 | 0.007 | 1.805 |
| Memory effect | -0.003 | 0.007 | -0.361 |
| CMLE effect | -0.073 | 0.066 | -1.102 |
| | Main and Interaction Effects | | |
| Intercept | 0.01 | 0.003 | 3.7 |
| Incentive effect | -0.002 | 0.003 | -0.84 |
| Curiosity effect | 0.013 | 0.007 | 1.705 |
| Curiosity incentive interaction | 0.002 | 0.011 | 0.172 |
| Memory effect | -0.003 | 0.007 | -0.417 |
| Memory incentive interaction | 0.015 | 0.011 | 1.362 |
| CMLE effect | -0.071 | 0.067 | -1.048 |
| CMLE incentive interaction | -0.031 | 0.09 | -0.343 |

*Note*. Three different models were computed targeting different effects. Each LME specified crossed random intercepts for both subjects in each pair. Incentive effect was effect-coded (control = -1, incentive = 1). Rather than residualized values, Fisher's *z*-transformed and grand-mean centred pairwise correlation values were used. *p* values are omitted as the *lme4* package does not compute them by default. SE = standard error. CMLE = Curiosity-motivated learning enhancement.

**Table S12**

*Whole Brain Results Of ISFC And Incentive Effects Therein Specifying aHPC And VTA/SN As Seeds*

| Cluster | Macro Label | Direction | Cluster Size | Maximum Intensity | | | |
|---|---|---|---|---|---|---|---|
| | | | | t | RL | AP | IS |
| | | Average ISFC | | | | | |
| aHPC seed | | | | | | | |
| | L Middle Temporal Gyrus | | | | | | |
| | L Middle Frontal Gyrus | | | | | | |
| | L Superior Medial Gyrus | | | | | | |
| | L Superior Frontal Gyrus | | | | | | |
| | R Middle Temporal Gyrus | | | | | | |
| | R Superior Medial Gyrus | | | | | | |
| | L Precuneus | | | | | | |
| | L Calcarine Gyrus | | | | | | |
| | L Lingual Gyrus | | | | | | |
| | R Superior Frontal Gyrus | | | | | | |
| | L Middle Cingulate Cortex | | | | | | |
| | L Cuneus | | | | | | |
| | R Precuneus | | | | | | |
| | R Calcarine Gyrus | | | | | | |
| | L Inferior Frontal Gyrus (p. Orbitalis) | | | | | | |
| | R Lingual Gyrus | | | | | | |
| | L SMA | | | | | | |
| | R Superior Temporal Gyrus | | | | | | |
| | L Inferior Temporal Gyrus | | | | | | |
| | L Insula Lobe | | | | | | |
| | L Anterior Cingulate Cortex | | | | | | |
| | R Middle Cingulate Cortex | | | | | | |
| | L Angular Gyrus | | | | | | |
| | L Inferior Frontal Gyrus (p. Triangularis) | | | | | | |
| | R Inferior Temporal Gyrus | | | | | | |
| | L Thalamus | | | | | | |
| | R Cuneus | | | | | | |
| | R Anterior Cingulate Cortex | | | | | | |
| | R Insula Lobe | | | | | | |
| | L Fusiform Gyrus | | | | | | |
| | L Putamen | | | | | | |
| | R Hippocampus | | | | | | |
| 1 | R Inferior Frontal Gyrus (p. | positive | 17870 | 9.9588 | -19.5 | -13.5 | -7.5 |

Orbitalis)
R Thalamus
L Hippocampus
R Medial Temporal Pole
R Middle Frontal Gyrus
L Temporal Pole
R Temporal Pole
R ParaHippocampal Gyrus
L Superior Temporal Gyrus
L Inferior Parietal Lobule
R Caudate Nucleus
L Mid Orbital Gyrus
R Putamen
L Caudate Nucleus
R Mid Orbital Gyrus
R Cerebellum (IV-V)
R Fusiform Gyrus
L Cerebellum (IV-V)
L Rolandic Operculum
L Superior Occipital Gyrus
L ParaHippocampal Gyrus
L Posterior Cingulate
Cortex
L Medial Temporal Pole
R Rolandic Operculum
R SMA
R Middle Orbital Gyrus
L Middle Occipital Gyrus
L Superior Orbital Gyrus
L Inferior Frontal Gyrus (p.
Opercularis)
R Heschls Gyrus
R Cerebellum (VI)
R Superior Orbital Gyrus
L Middle Orbital Gyrus
L Heschls Gyrus
R Superior Occipital Gyrus
R Amygdala
R Olfactory cortex
L Olfactory cortex
R Posterior Cingulate
Cortex
L Amygdala
R Pallidum
L Cerebellum (VI)
L Precentral Gyrus
L Rectal Gyrus
L Pallidum
L Postcentral Gyrus
R Inferior Frontal Gyrus (p.

| | | | | | | | |
|---|---|---|---|---|---|---|---|
| | Opercularis)<br>Cerebellar Vermis (4/5)<br>L Paracentral Lobule<br>L Superior Parietal Lobule<br>L SupraMarginal Gyrus<br>R Rectal Gyrus<br>R Paracentral Lobule<br>L Inferior Occipital Gyrus<br>R Inferior Frontal Gyrus (p. Triangularis)<br>Cerebellar Vermis (3)<br>L Cerebellum (III)<br>L Cerebellum (Crus 1)<br>R Cerebellum (III) | | | | | | |
| 2 | L Postcentral Gyrus<br>L Precentral Gyrus | positive | 535 | 8.5217 | 16.5 | 34.5 | 64.5 |
| 3 | R Angular Gyrus<br>R Inferior Parietal Lobule<br>R Middle Occipital Gyrus | positive | 531 | 7.0422 | -58.5 | 58.5 | 28.5 |
| 4 | R Precentral Gyrus<br>R Postcentral Gyrus | positive | 305 | 7.3448 | -22.5 | 25.5 | 61.5 |
| 5 | L Cerebellum (Crus 2)<br>L Cerebellum (Crus 1) | positive | 241 | 5.837 | 40.5 | 82.5 | -34.5 |
| 6 | R Cerebellum (Crus 1)<br>R Cerebellum (Crus 2) | positive | 155 | 6.2307 | -40.5 | 76.5 | -34.5 |
| 7 | Brainstem | positive | 69 | 5.9908 | 4.5 | 28.5 | -40.5 |
| 8 | R Middle Frontal Gyrus<br>R Middle Orbital Gyrus | positive | 40 | 4.2492 | -40.5 | -52.5 | 7.5 |
| 9 | Cerebellar Vermis (8)<br>L Cerebellum (VIII) | positive | 22 | 4.8611 | -1.5 | 70.5 | -37.5 |
| 10 | R Superior Parietal Lobule<br>R Postcentral Gyrus<br>R Inferior Parietal Lobule<br>R Precuneus | negative | 713 | -7.6371 | -13.5 | 55.5 | 70.5 |
| 11 | L Superior Parietal Lobule<br>L Inferior Parietal Lobule<br>L Postcentral Gyrus<br>L Precuneus | negative | 495 | -6.4146 | 22.5 | 61.5 | 58.5 |

| | | | | | | | |
|---|---|---|---|---|---|---|---|
| 12 | L Middle Occipital Gyrus<br>L Inferior Occipital Gyrus<br>L Lingual Gyrus<br>L Calcarine Gyrus | negative | 488 | -6.6946 | 28.5 | 94.5 | -16.5 |
| 13 | R Superior Frontal Gyrus<br>R Precentral Gyrus<br>R Middle Frontal Gyrus | negative | 343 | -7.03 | -34.5 | 7.5 | 55.5 |
| 14 | R SupraMarginal Gyrus<br>R Postcentral Gyrus | negative | 334 | -6.3619 | -52.5 | 37.5 | 28.5 |
| 15 | R Precentral Gyrus<br>R Inferior Frontal Gyrus (p. Opercularis)<br>R Rolandic Operculum | negative | 208 | -7.545 | -64.5 | -10.5 | 16.5 |
| 16 | R Inferior Occipital Gyrus<br>R Lingual Gyrus<br>R Calcarine Gyrus | negative | 196 | -5.0473 | -22.5 | 100.5 | -13.5 |
| 17 | L Precentral Gyrus<br>L Superior Frontal Gyrus | negative | 159 | -6.6714 | 25.5 | 10.5 | 58.5 |
| 18 | L SupraMarginal Gyrus<br>L Postcentral Gyrus | negative | 97 | -5.0521 | 58.5 | 25.5 | 28.5 |
| 19 | L SupraMarginal Gyrus<br>L Superior Temporal Gyrus | negative | 69 | -6.7032 | 52.5 | 37.5 | 25.5 |
| 20 | L Precentral Gyrus<br>L Postcentral Gyrus | negative | 44 | -5.9077 | 58.5 | -1.5 | 43.5 |
| VTA/SN seed | | | | | | | |
| 1 | L Middle Frontal Gyrus<br>R Middle Frontal Gyrus<br>L Middle Temporal Gyrus<br>R Middle Temporal Gyrus<br>R Superior Frontal Gyrus<br>L Cerebellum (Crus 1)<br>L Inferior Temporal Gyrus<br>L Superior Medial Gyrus<br>L Superior Frontal Gyrus<br>L Inferior Frontal Gyrus (p. Triangularis)<br>R Cerebellum (Crus 1)<br>R Inferior Temporal Gyrus<br>L Lingual Gyrus<br>R Lingual Gyrus | positive | 32507 | 16.502 | 10.5 | 85.5 | 4.5 |

L Precuneus
L Calcarine Gyrus
R Inferior Frontal Gyrus (p. Triangularis)
L Cerebellum (Crus 2)
R Superior Medial Gyrus
R Angular Gyrus
R Precuneus
R Superior Temporal Gyrus
L Inferior Frontal Gyrus (p. Orbitalis)
R Inferior Frontal Gyrus (p. Orbitalis)
L Inferior Parietal Lobule
R Cerebellum (Crus 2)
L Cuneus
L Middle Cingulate Cortex
R Calcarine Gyrus
L Insula Lobe
R Cuneus
R Middle Cingulate Cortex
R Cerebellum (VI)
L Precentral Gyrus
R Insula Lobe
R Fusiform Gyrus
L Angular Gyrus
L SMA
L Thalamus
L Fusiform Gyrus
R Inferior Parietal Lobule
R Middle Orbital Gyrus
R Temporal Pole
L Anterior Cingulate Cortex
L Temporal Pole
L Putamen
R Inferior Frontal Gyrus (p. Opercularis)
R Thalamus
R Anterior Cingulate Cortex
L Inferior Frontal Gyrus (p. Opercularis)
R Caudate Nucleus
R Medial Temporal Pole
L Postcentral Gyrus
L Middle Orbital Gyrus
L Superior Temporal Gyrus
L Superior Occipital Gyrus
L Caudate Nucleus
R ParaHippocampal Gyrus
R SMA

R Putamen
R Cerebellum (VIII)
R Superior Orbital Gyrus
R SupraMarginal Gyrus
R Cerebellum (IV-V)
L Middle Occipital Gyrus
R Hippocampus
R Superior Occipital Gyrus
L Hippocampus
L Medial Temporal Pole
L Cerebellum (VI)
R Middle Occipital Gyrus
L Superior Orbital Gyrus
L ParaHippocampal Gyrus
Cerebellar Vermis (4/5)
R Cerebellum (VII)
R Pallidum
L Cerebellum (IV-V)
L Superior Parietal Lobule
L Pallidum
L Posterior Cingulate
Cortex
L Cerebellum (VII)
L SupraMarginal Gyrus
L Amygdala
R Amygdala
L Cerebellum (IX)
L Rolandic Operculum
L Cerebellum (VIII)
R Olfactory cortex
R Superior Parietal Lobule
R Precentral Gyrus
R Posterior Cingulate
Cortex
L Olfactory cortex
L Cerebellum (III)
R Rectal Gyrus
R Cerebellum (IX)
Cerebellar Vermis (8)
R Rolandic Operculum
Cerebellar Vermis (3)
Cerebellar Vermis (9)
Cerebellar Vermis (6)
L Inferior Occipital Gyrus
R Heschls Gyrus
R Mid Orbital Gyrus
Cerebellar Vermis (7)
L Rectal Gyrus
L Heschls Gyrus
R Cerebellum (III)

| | | | | | | | |
|---|---|---|---|---|---|---|---|
| | Cerebellar Vermis (1/2)<br>Cerebellar Vermis (10)<br>L Paracentral Lobule<br>R Paracentral Lobule<br>L Mid Orbital Gyrus | | | | | | |
| 2 | R Precentral Gyrus<br>R Postcentral Gyrus | positive | 33 | 7.5112 | -37.5 | 16.5 | 37.5 |
| 3 | L Middle Occipital Gyrus<br>L Inferior Occipital Gyrus<br>L Lingual Gyrus<br>L Middle Temporal Gyrus<br>L Calcarine Gyrus | negative | 2676 | -12.85 | -37.5 | 37.5 | 58.5 |
| 4 | R Inferior Occipital Gyrus<br>R Middle Occipital Gyrus<br>R Middle Temporal Gyrus<br>R Lingual Gyrus<br>R Calcarine Gyrus | negative | 890 | -11.985 | 19.5 | 103.5 | 4.5 |
| 5 | R Inferior Occipital Gyrus<br>R Middle Occipital Gyrus<br>R Middle Temporal Gyrus<br>R Lingual Gyrus<br>R Calcarine Gyrus | negative | 806 | -13.049 | -22.5 | 97.5 | -4.5 |
| 6 | L Postcentral Gyrus<br>L SupraMarginal Gyrus<br>L Precentral Gyrus | negative | 196 | -6.3416 | 61.5 | 16.5 | 37.5 |
| 7 | L SupraMarginal Gyrus<br>L Superior Temporal Gyrus | negative | 82 | -7.1019 | 49.5 | 40.5 | 28.5 |
| 8 | L Mid Orbital Gyrus<br>L Rectal Gyrus<br>R Rectal Gyrus<br>R Mid Orbital Gyrus | negative | 78 | -4.9271 | 1.5 | -52.5 | -13.5 |
| 9 | R Precentral Gyrus<br>R Postcentral Gyrus | negative | 65 | -7.109 | -61.5 | -10.5 | 34.5 |
| 10 | L Cerebellum (VIII) | negative | 58 | -6.1588 | 28.5 | 52.5 | -46.5 |
| 11 | R Superior Occipital Gyrus<br>R Superior Parietal Lobule | negative | 52 | -7.9278 | -28.5 | 82.5 | 43.5 |

| | | | | | | | |
|---|---|---|---|---|---|---|---|
| 12 | L Superior Occipital Gyrus<br>L Middle Occipital Gyrus<br>L Superior Parietal Lobule | negative | 50 | -6.8917 | 25.5 | 85.5 | 40.5 |
| 13 | R SupraMarginal Gyrus<br>R Rolandic Operculum | negative | 41 | -5.4704 | -40.5 | 25.5 | 22.5 |
| 14 | L Cerebellum (Crus 2)<br>L Cerebellum (VII)<br>Cerebellar Vermis (7) | negative | 25 | -5.8236 | -1.5 | 76.5 | -31.5 |
| 15 | R Rolandic Operculum<br>R Insula Lobe | negative | 21 | -6.0122 | -40.5 | -1.5 | 16.5 |
| | ISFC Incentive Effects | | | | | | |
| aHPC seed | | | | | | | |
| | *no clusters survive thresholding* | | | | | | |
| VTA/SN seed | | | | | | | |
| 1 | L Superior Temporal Gyrus<br>L SupraMarginal Gyrus | I > C | 23 | -4.3753 | 55.5 | 28.5 | 16.5 |
| 2 | L Calcarine Gyrus<br>L Inferior Occipital Gyrus<br>L Middle Occipital Gyrus<br>L Lingual Gyrus | I < C | 56 | 5.1537 | 10.5 | 100.5 | -4.5 |

*Note.* Results are thresholded at $p < 0.001$, cluster-extent corrected at $k = 20$ (equivalent to per-cluster $\alpha = 0.05$) for whole brain analysis. The table corresponds to Figure S9. $t = t$ value, RL = right to left, AP = anterior to posterior, IS = inferior to superior. RAI orientation is applied where negative values indicate right, anterior, inferior.

**Table S13**

*Whole Brain Results Of ISFC-RSA For Each Behavioural Effect Of Interest Specifying aHPC And VTA/SN As Seeds*

| Cluster | Macro Label | Direction | Cluster Size | Maximum Intensity | | | |
|---|---|---|---|---|---|---|---|
| | | | | t | RL | AP | IS |
| | | | Curiosity | | | | |
| aHPC seed | | | | | | | |
| | | *no clusters survive thresholding* | | | | | |
| VTA/SN seed | | | | | | | |
| 1 | R Superior Medial Gyrus<br>L Superior Medial Gyrus | positive | 49 | 5.1764 | -4.5 | -40.5 | 52.5 |
| 2 | L Inferior Frontal Gyrus (p. Orbitalis)<br>L Insula Lobe | positive | 44 | 5.288 | 31.5 | -25.5 | -7.5 |
| 3 | L Inferior Parietal Lobule<br>L Middle Occipital Gyrus | positive | 31 | 4.4912 | 34.5 | 82.5 | 37.5 |
| 4 | Dorsal pons | positive | 30 | 5.7828 | 1.5 | 22.5 | -25.5 |
| 5 | R Superior Medial Gyrus<br>R SMA | positive | 26 | 4.9863 | -10.5 | -22.5 | 58.5 |
| | | | Memory | | | | |
| aHPC seed | | | | | | | |
| | | *no clusters survive thresholding* | | | | | |
| VTA/SN seed | | | | | | | |
| 1 | L Cuneus<br>L Calcarine Gyrus<br>R Calcarine Gyrus<br>R Cuneus | positive | 480 | 5.3752 | -4.5 | 85.5 | 25.5 |
| 2 | R Middle Frontal Gyrus<br>R Superior Frontal Gyrus | positive | 227 | 5.7297 | -28.5 | -40.5 | 34.5 |

| | | | | | | | |
|---|---|---|---|---|---|---|---|
| 3 | R Anterior Cingulate Cortex<br>L Superior Medial Gyrus<br>R Middle Cingulate Cortex<br>R Superior Medial Gyrus<br>L Anterior Cingulate Cortex | positive | 205 | 5.3584 | -4.5 | -28.5 | 34.5 |
| 4 | L Postcentral Gyrus<br>L Precentral Gyrus | positive | 82 | 4.8762 | 46.5 | 22.5 | 55.5 |
| 5 | R Angular Gyrus<br>R Inferior Parietal Lobule | positive | 77 | 4.5695 | -43.5 | 70.5 | 46.5 |
| 6 | R Insula Lobe | positive | 65 | 5.1114 | -31.5 | -28.5 | 4.5 |
| 7 | R Inferior Parietal Lobule<br>R SupraMarginal Gyrus | positive | 52 | 4.7107 | -55.5 | 43.5 | 43.5 |
| 8 | R Middle Frontal Gyrus | positive | 47 | 5.2007 | -40.5 | -13.5 | 58.5 |
| 9 | L SMA<br>R SMA | positive | 42 | 4.3257 | 4.5 | -16.5 | 58.5 |
| 10 | L Angular Gyrus<br>L Inferior Parietal Lobule | positive | 39 | 4.7107 | 43.5 | 64.5 | 46.5 |
| 11 | R Middle Frontal Gyrus | positive | 35 | 4.7809 | -43.5 | -19.5 | 43.5 |
| 12 | R Inferior Temporal Gyrus | positive | 32 | 4.8716 | -58.5 | 55.5 | -13.5 |
| 13 | L Middle Frontal Gyrus | positive | 32 | 4.5784 | 37.5 | -52.5 | 19.5 |
| 14 | R Middle Temporal Gyrus | positive | 28 | 4.719 | -64.5 | 34.5 | -13.5 |
| 15 | R Cuneus<br>R Superior Occipital Gyrus | positive | 27 | 4.4552 | -16.5 | 91.5 | 28.5 |
| 16 | L Caudate Nucleus<br>L Putamen | positive | 23 | 5.1853 | 16.5 | -16.5 | -1.5 |
| 17 | L Cerebellum (Crus 2) | positive | 20 | 5.2813 | 46.5 | 64.5 | -52.5 |
| 18 | L Middle Occipital Gyrus<br>L Inferior Occipital Gyrus | negative | 21 | -4.4753 | 22.5 | 97.5 | -1.5 |

Curiosity-Motivated Learning Enhancement

aHPC seed

| | | | | | | | |
|---|---|---|---|---|---|---|---|
| 1 | R Lingual Gyrus<br>R Calcarine Gyrus<br>R Fusiform Gyrus<br>R Cuneus<br>R Superior Occipital Gyrus | positive | 377 | 6.7356 | -16.5 | 79.5 | -7.5 |
| 2 | L Calcarine Gyrus<br>L Superior Occipital Gyrus<br>L Cuneus<br>L Lingual Gyrus | positive | 144 | 6.2075 | 7.5 | 94.5 | 13.5 |
| 3 | L Middle Frontal Gyrus<br>L Inferior Frontal Gyrus (p. Triangularis)<br>L Inferior Frontal Gyrus (p. Opercularis) | positive | 137 | 5.858 | 43.5 | -28.5 | 34.5 |
| 4 | L Inferior Parietal Lobule<br>L Angular Gyrus<br>L Superior Parietal Lobule | positive | 112 | 6.1657 | 28.5 | 76.5 | 49.5 |
| 5 | L Inferior Frontal Gyrus (p. Orbitalis)<br>L Inferior Frontal Gyrus (p. Triangularis)<br>L Middle Orbital Gyrus<br>L Middle Frontal Gyrus | positive | 105 | 4.7188 | 49.5 | -34.5 | -7.5 |
| 6 | R Angular Gyrus<br>R Superior Occipital Gyrus<br>R Superior Parietal Lobule | positive | 93 | 5.3644 | -37.5 | 67.5 | 49.5 |
| 7 | R Middle Temporal Gyrus | positive | 71 | 4.6236 | 61.5 | 55.5 | 19.5 |
| 8 | R Middle Temporal Gyrus<br>R Superior Temporal Gyrus | positive | 55 | 4.5192 | -58.5 | 34.5 | -1.5 |
| 9 | L Lingual Gyrus<br>L Cerebellum (Crus 1) | positive | 53 | 5.3504 | 13.5 | 85.5 | -13.5 |
| 10 | L SMA<br>R SMA | positive | 51 | 4.6741 | 1.5 | -19.5 | 55.5 |
| 11 | R Inferior Parietal Lobule | positive | 49 | 4.6818 | -46.5 | 49.5 | 49.5 |
| 12 | L Middle Frontal Gyrus | positive | 45 | 4.657 | 28.5 | -49.5 | 19.5 |

| | | | | | | | |
|---|---|---|---|---|---|---|---|
| 13 | R Fusiform Gyrus<br>R Lingual Gyrus | positive | 35 | 5.2247 | -22.5 | 58.5 | -10.5 |
| 14 | L Superior Orbital Gyrus<br>L Middle Orbital Gyrus<br>L Superior Frontal Gyrus | positive | 33 | 4.731 | 22.5 | -61.5 | -7.5 |
| 15 | R Precuneus | positive | 26 | 4.306 | -7.5 | 73.5 | 49.5 |
| 16 | L Superior Frontal Gyrus<br>L Superior Medial Gyrus | positive | 20 | 4.4976 | 16.5 | -64.5 | 13.5 |
| 17 | L Inferior Occipital Gyrus<br>L Middle Occipital Gyrus | negative | 112 | -6.9394 | 19.5 | 100.5 | -10.5 |
| VTA/SN seed | | | | | | | |
| 1 | L Precentral Gyrus<br>R Superior Frontal Gyrus<br>L SMA<br>R SMA<br>R Precentral Gyrus<br>L Superior Frontal Gyrus | positive | 804 | 10.396 | 25.5 | 13.5 | 58.5 |
| 2 | L Superior Parietal Lobule<br>L Postcentral Gyrus<br>L Inferior Parietal Lobule<br>L SupraMarginal Gyrus | positive | 749 | 9.5435 | 25.5 | 67.5 | 64.5 |
| 3 | R Postcentral Gyrus<br>R Superior Parietal Lobule<br>R SupraMarginal Gyrus<br>R Inferior Parietal Lobule | positive | 407 | 9.5024 | -31.5 | 40.5 | 49.5 |
| 4 | R Inferior Occipital Gyrus<br>R Calcarine Gyrus<br>R Middle Occipital Gyrus | positive | 193 | 8.7441 | -25.5 | 103.5 | 1.5 |
| 5 | L Middle Occipital Gyrus<br>L Inferior Occipital Gyrus | positive | 155 | 7.9807 | 22.5 | 103.5 | -7.5 |
| 6 | L Precentral Gyrus<br>L Postcentral Gyrus | positive | 90 | 6.7871 | 61.5 | -7.5 | 28.5 |
| 7 | L Middle Occipital Gyrus | positive | 42 | 5.9038 | 37.5 | 70.5 | 4.5 |
| 8 | R Precentral Gyrus | positive | 42 | 6.2684 | -64.5 | -7.5 | 25.5 |
| 9 | L Cerebellum (VIII) | positive | 29 | 5.3834 | 25.5 | 64.5 | -52.5 |

| | | | | | | | |
|---|---|---|---|---|---|---|---|
| 10 | L Rolandic Operculum | positive | 28 | 6.5409 | 52.5 | 4.5 | 13.5 |
| | L Middle Frontal Gyrus | | | | | | |
| | R Middle Frontal Gyrus | | | | | | |
| | L Superior Medial Gyrus | | | | | | |
| | R Superior Frontal Gyrus | | | | | | |
| | L Cerebellum (Crus 1) | | | | | | |
| | L Inferior Frontal Gyrus (p. Triangularis) | | | | | | |
| | R Superior Medial Gyrus | | | | | | |
| | R Cerebellum (Crus 1) | | | | | | |
| | R Inferior Frontal Gyrus (p. Triangularis) | | | | | | |
| | R Inferior Frontal Gyrus (p. Orbitalis) | | | | | | |
| | R Middle Temporal Gyrus | | | | | | |
| | L Calcarine Gyrus | | | | | | |
| | L Cerebellum (Crus 2) | | | | | | |
| | L Cuneus | | | | | | |
| | L Superior Frontal Gyrus | | | | | | |
| | R Angular Gyrus | | | | | | |
| | L Middle Temporal Gyrus | | | | | | |
| | R Inferior Temporal Gyrus | | | | | | |
| | L Inferior Frontal Gyrus (p. Orbitalis) | | | | | | |
| | R Precuneus | | | | | | |
| | R Cuneus | | | | | | |
| | L Precuneus | | | | | | |
| | L Lingual Gyrus | | | | | | |
| | R Inferior Parietal Lobule | | | | | | |
| | R Calcarine Gyrus | | | | | | |
| | R Lingual Gyrus | | | | | | |
| | L Inferior Temporal Gyrus | | | | | | |
| | R Middle Orbital Gyrus | | | | | | |
| | R Cerebellum (Crus 2) | | | | | | |
| | R Caudate Nucleus | | | | | | |
| | L SMA | | | | | | |
| | R Anterior Cingulate Cortex | | | | | | |
| | R Inferior Frontal Gyrus (p. Opercularis) | | | | | | |
| | L Middle Orbital Gyrus | | | | | | |
| | L Anterior Cingulate Cortex | | | | | | |
| 11 | L Superior Occipital Gyrus | negative | 18122 | -11.709 | -49.5 | 52.5 | 52.5 |

| | | | | | | | |
|---|---|---|---|---|---|---|---|
| 12 | L Inferior Parietal Lobule<br>L Angular Gyrus<br>L Middle Occipital Gyrus | negative | 951 | -9.872 | 46.5 | 58.5 | 55.5 |
| 13 | L Inferior Temporal Gyrus | negative | 112 | -5.7714 | 43.5 | -4.5 | -40.5 |
| 14 | R ParaHippocampal Gyrus<br>R Amygdala<br>R Hippocampus | negative | 81 | -5.0832 | -25.5 | 10.5 | -34.5 |
| 15 | Fusiform Gyrus<br>Inferior Temporal Gyrus<br>ParaHippocampal Gyrus | negative | 55 | -5.4851 | 31.5 | 7.5 | -34.5 |
| 16 | L Insula Lobe<br>L Superior Temporal Gyrus | negative | 22 | -5.4636 | 37.5 | 4.5 | -7.5 |
| 17 | R ParaHippocampal Gyrus<br>R Hippocampus | negative | 21 | -4.6463 | -28.5 | 19.5 | -22.5 |

*Note.* Results are thresholded at $p < 0.001$, cluster-extent corrected at $k = 20$ (equivalent to per-cluster $\alpha = 0.05$) for whole brain analysis. The table corresponds to Figure S10. $t = t$ value, RL = right to left, AP = anterior to posterior, IS = inferior to superior. RAI orientation is applied where negative values indicate right, anterior, inferior.

**Table S14**

*Whole Brain Results Of ISFC-RSA For The Interaction Between The Incentive Manipulation And Each Behavioural Effect Of Interest Specifying aHPC And VTA/SN As Seeds*

| Cluster | Macro Label | Direction | Cluster Size | Maximum Intensity | | | |
|---|---|---|---|---|---|---|---|
| | | | | t | RL | AP | IS |
| | | | Curiosity | | | | |
| aHPC seed | | | | | | | |
| | | *no clusters survive thresholding* | | | | | |
| VTA/SN seed | | | | | | | |
| | | *no clusters survive thresholding* | | | | | |
| | | | Memory | | | | |
| aHPC seed | | | | | | | |
| | | *no clusters survive thresholding* | | | | | |
| VTA/SN seed | | | | | | | |
| 1 | R Middle Occipital Gyrus<br>R Superior Occipital Gyrus<br>R Calcarine Gyrus<br>R Inferior Occipital Gyrus | I > C | 20 | -4.2192 | -28.5 | 100.5 | 1.5 |
| 2 | L Posterior Cingulate Cortex<br>L Middle Cingulate Cortex<br>R Middle Cingulate Cortex<br>R Precuneus | I < C | 79 | 4.7595 | -1.5 | 49.5 | 25.5 |
| 3 | L Middle Frontal Gyrus<br>L Middle Orbital Gyrus | I < C | 31 | 5.3363 | 43.5 | -55.5 | 1.5 |
| | | Curiosity-Motivated Learning Enhancement | | | | | |
| aHPC seed | | | | | | | |
| 1 | L Inferior Occipital Gyrus | I < C | 28 | 5.2033 | 43.5 | 70.5 | -4.5 |
| VTA/SN seed | | | | | | | |
| 1 | L Precuneus<br>R Precuneus | I > C | 264 | -5.5263 | 13.5 | 73.5 | 46.5 |

| | | | | | | | |
|---|---|---|---|---|---|---|---|
| 2 | R Middle Frontal Gyrus<br>R Middle Orbital Gyrus | I > C | 149 | -5.7355 | -37.5 | -61.5 | 7.5 |
| 3 | L Middle Frontal Gyrus<br>L Middle Orbital Gyrus | I > C | 110 | -4.9382 | 40.5 | -49.5 | -1.5 |
| 4 | R Inferior Parietal Lobule | I > C | 98 | -5.4174 | -49.5 | 49.5 | 58.5 |
| 5 | L SupraMarginal Gyrus | I > C | 93 | -6.7075 | 64.5 | 49.5 | 31.5 |
| 6 | R Middle Temporal Gyrus | I > C | 67 | -4.9715 | -61.5 | 28.5 | -4.5 |
| 7 | L Inferior Parietal Lobule | I > C | 67 | -4.7524 | 37.5 | 52.5 | 40.5 |
| 8 | L Superior Medial Gyrus<br>R Superior Medial Gyrus | I > C | 51 | -4.3063 | -1.5 | -25.5 | 37.5 |
| 9 | R Cuneus<br>R Precuneus | I > C | 47 | -4.633 | -19.5 | 70.5 | 28.5 |
| 10 | R Angular Gyrus | I > C | 47 | -4.6916 | -43.5 | 73.5 | 46.5 |
| 11 | L Superior Frontal Gyrus<br>L SMA | I > C | 39 | -5.4058 | 19.5 | -16.5 | 67.5 |
| 12 | R Middle Frontal Gyrus | I > C | 33 | -5.2262 | -40.5 | -10.5 | 55.5 |
| 13 | L Middle Frontal Gyrus | I > C | 22 | -4.6668 | 52.5 | -28.5 | 34.5 |
| 14 | L Inferior Frontal Gyrus (p. Orbitalis) | I > C | 20 | -4.2697 | 40.5 | -22.5 | -10.5 |
| 15 | R Superior Parietal Lobule | I < C | 50 | 4.9234 | -25.5 | 67.5 | 64.5 |
| 16 | R Superior Frontal Gyrus<br>R Precentral Gyrus | I < C | 30 | 5.3659 | -25.5 | 10.5 | 55.5 |

*Note.* Results are thresholded at $p < 0.001$, cluster-extent corrected at $k = 20$ (equivalent to per-cluster $\alpha = 0.05$) for whole brain analysis. The table corresponds to Figure S11. $t$ = $t$ value, RL = right to left, AP = anterior to posterior, IS = inferior to superior. RAI orientation is applied where negative values indicate right, anterior, inferior. I = Incentive group, C = Control group.

### Supplementary Figures

#### Figure S1

*Trial Structure In The fMRI Experiment*

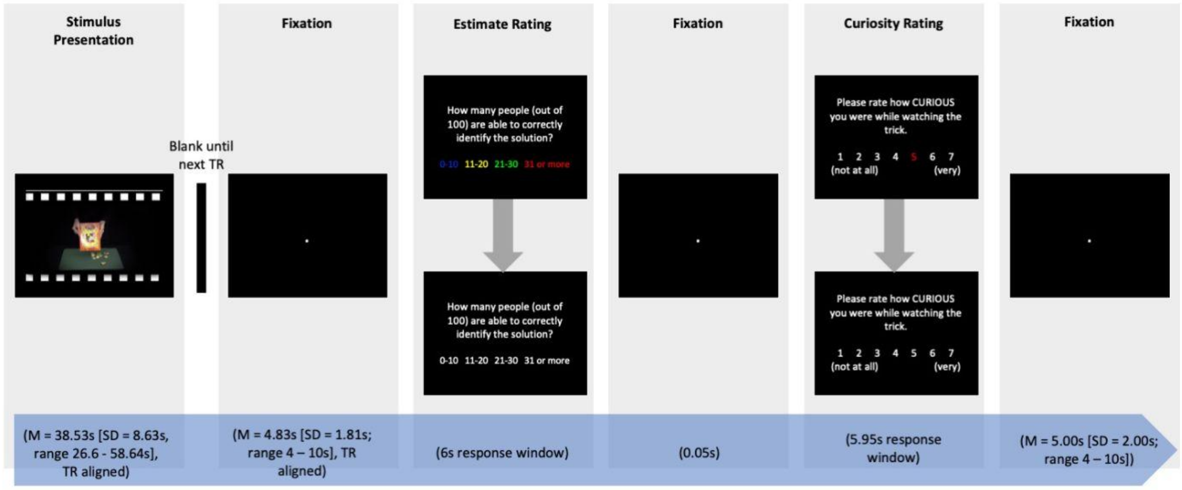

*Note*. This figure was adopted and reproduced from Meliss et al. (2022). The magic trick watching task was slightly adapted for the use inside the MRI scanner. More specifically, (1) the start of the magic trick presentation and the beginning of the fixation afterwards were aligned with the beginning of the acquisition of a volume ('TR aligned'), (2) jittered fixations were added between the stimulus presentation and ratings, and (3) and all response windows were fixed. The film framing surrounding the stimuli is for illustration purposes only and was not used in the experiment.

**Figure S2**

*Methodology To Determine The Optimal Lag To Apply During Concatenation*

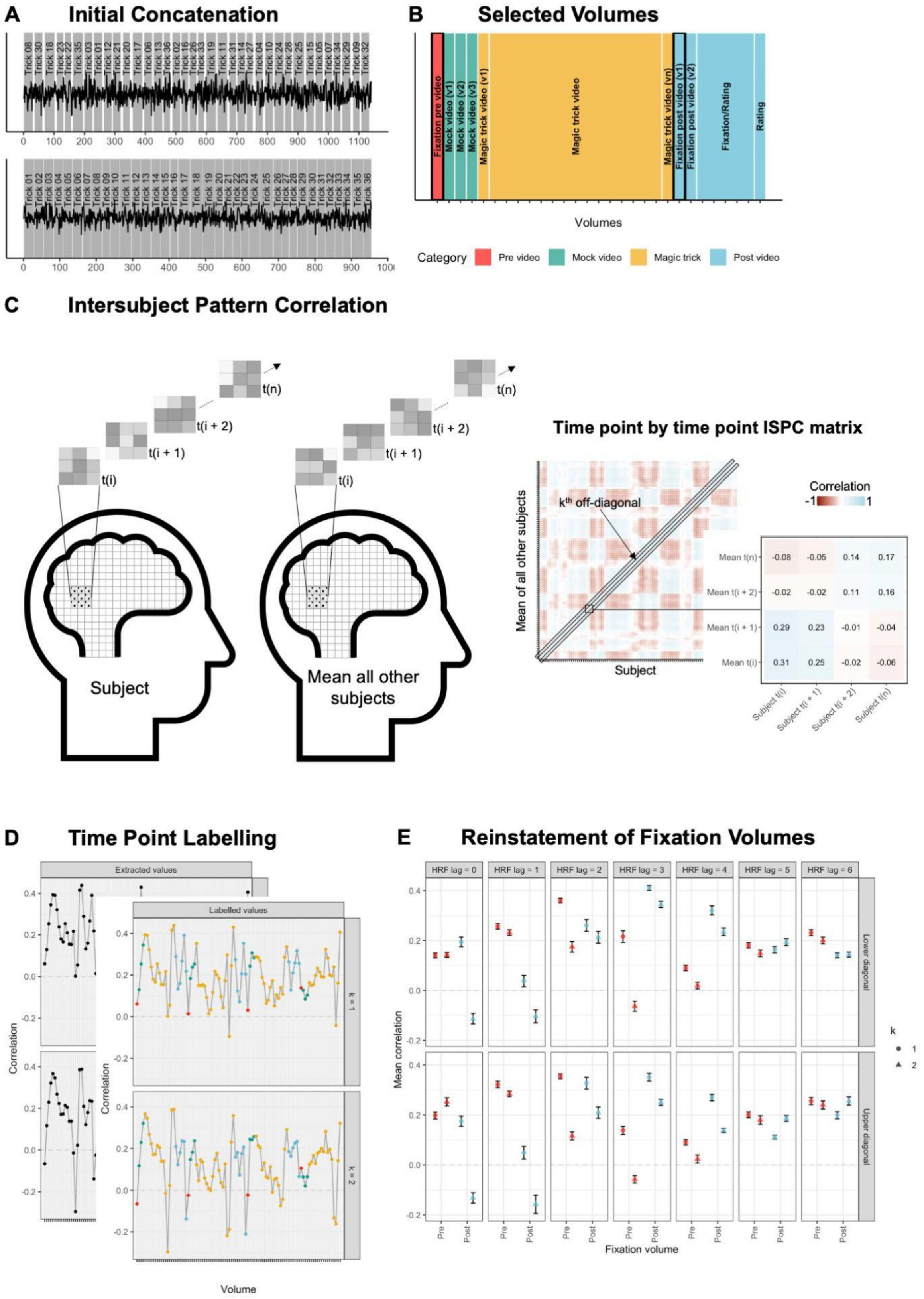

*Note*. Due to inconsistencies in the literature, the optimal HRF lag was determined based on data. (A) To reverse the pseudo-randomisation of stimuli, an initial concatenation step was applied. (B) During the initial concatenation, volumes before and after the magic trick video were selected and categorised depending on events in the experimental task. (C) The concatenated time series were used to extract the time course of the second visual cortex (V2) for each voxel at each time point. The data in time point t(i) in a given subject was correlated with time points from the mean time series of all other subjects creating a time point by time point intersubject pattern correlation (ISPC) matrix for each subject that was Fisher's *z*-transformed before computing the mean sample ISPC matrix. In the matrix, the $k^{th}$ off-diagonal represents the intersubject reinstatement of time point t(i) at time point t(i + k). The upper diagonal shows the reinstatement of the subject's response in the sample mean of all other subjects whereas the lower diagonal represents the reinstatement of the sample mean of all other subjects in the subject's response (figure adapted from Nastase et al., 2019). (D) For $1 \leq k \leq 2$, the upper and lower off-diagonals were extracted and labelled in correspondence to (B) without applying any HRF lags. Labels were also shifted for 1 volume ≤ HRF lag ≤ 6 volumes. (E) For the events 'Fixation pre video' and 'Fixation post video (volume 1)' (both highlighted in (B)), intersubject reinstatement (measured as correlation) data was averaged and plotted for each HRF lag and k. Error bars represent SE.

**Figure S3**

*Regions Of interest (ROIs)*

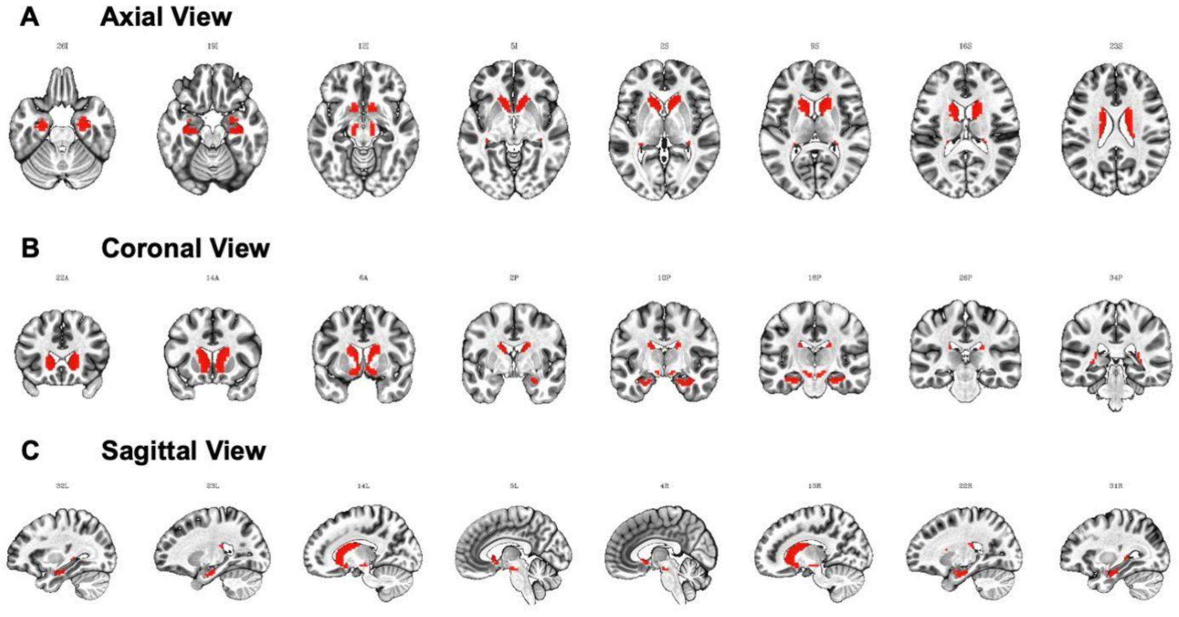

*Note*. The *a priori* defined ROIs were anterior hippocampus (aHPC), nucleus accumbens (NAcc), caudate nucleus (CN), and dopaminergic midbrain (VTA/SN) in (A) axial, (B) coronal, and (C) sagittal view. For details on how they were created, please refer to the main text. ROIs are shown on the ICBM 2009c Nonlinear Asymmetric Template and resampled to the EPI grid. Images are displayed in neurological orientation, where the left side of the brain is depicted on the left side of the image.

Figure S4

*Fixed Effects Estimates Of Curiosity, Monetary Incentive, And Their Interaction As A Function Of Confidence Cut-off Separately For Each Data Collection*

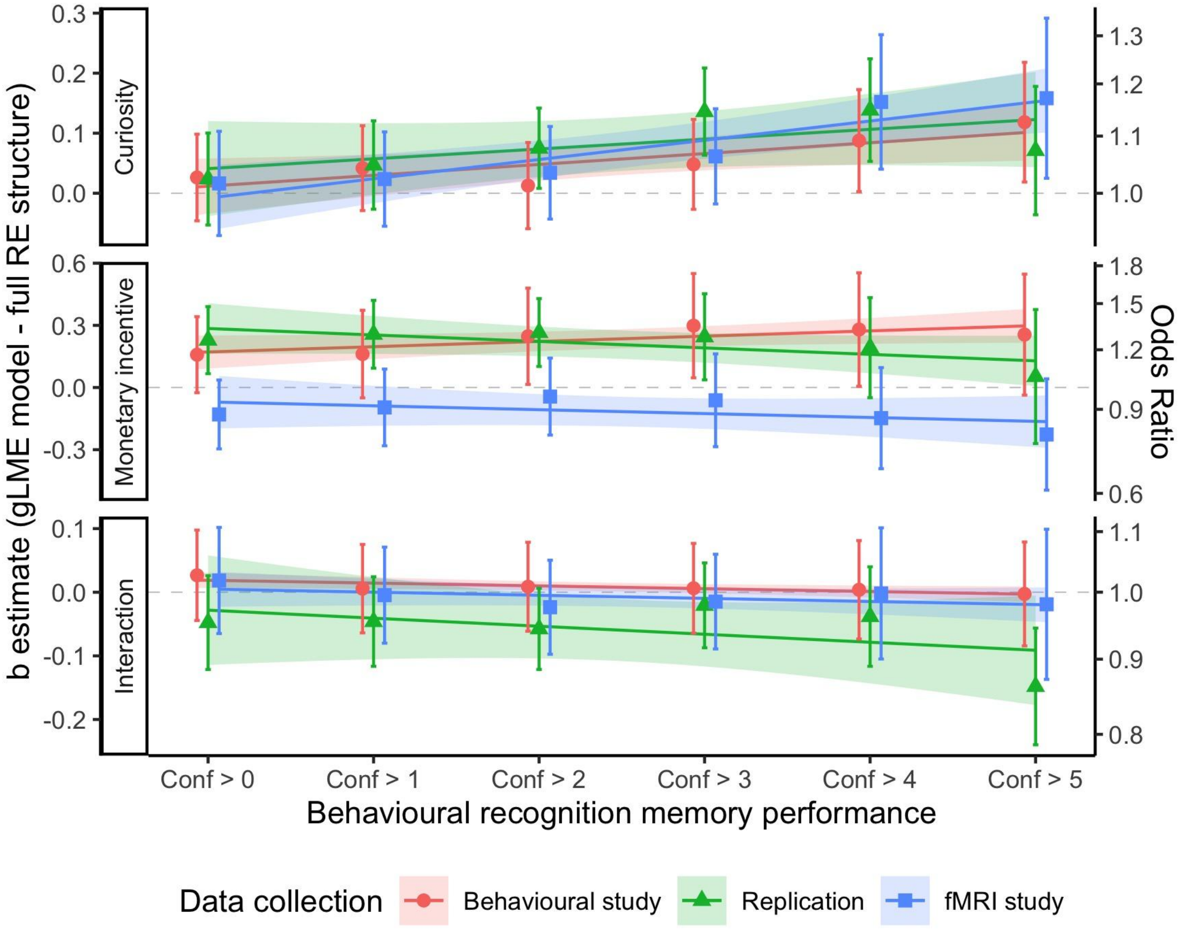

*Note*. The x-axis shows the gradual confidence cut-off and y-axis illustrates the integrated effect size (left - unstandardised, right - OR). Each panel shows one of the fixed effects specified in the gLME model. The b estimate from each data collection for each effect and confidence threshold is plotted and error bars indicate 95%-CI. The regression line illustrates the linear model predicting the effect with the gradual confidence cut-off. The data shown here was integrated and plotted in Figure 3 in the main text.

**Figure S5**

*Integrated Fixed Effects Of Curiosity, Monetary Incentive, And Their Interaction As A Function Of Confidence cut-off*

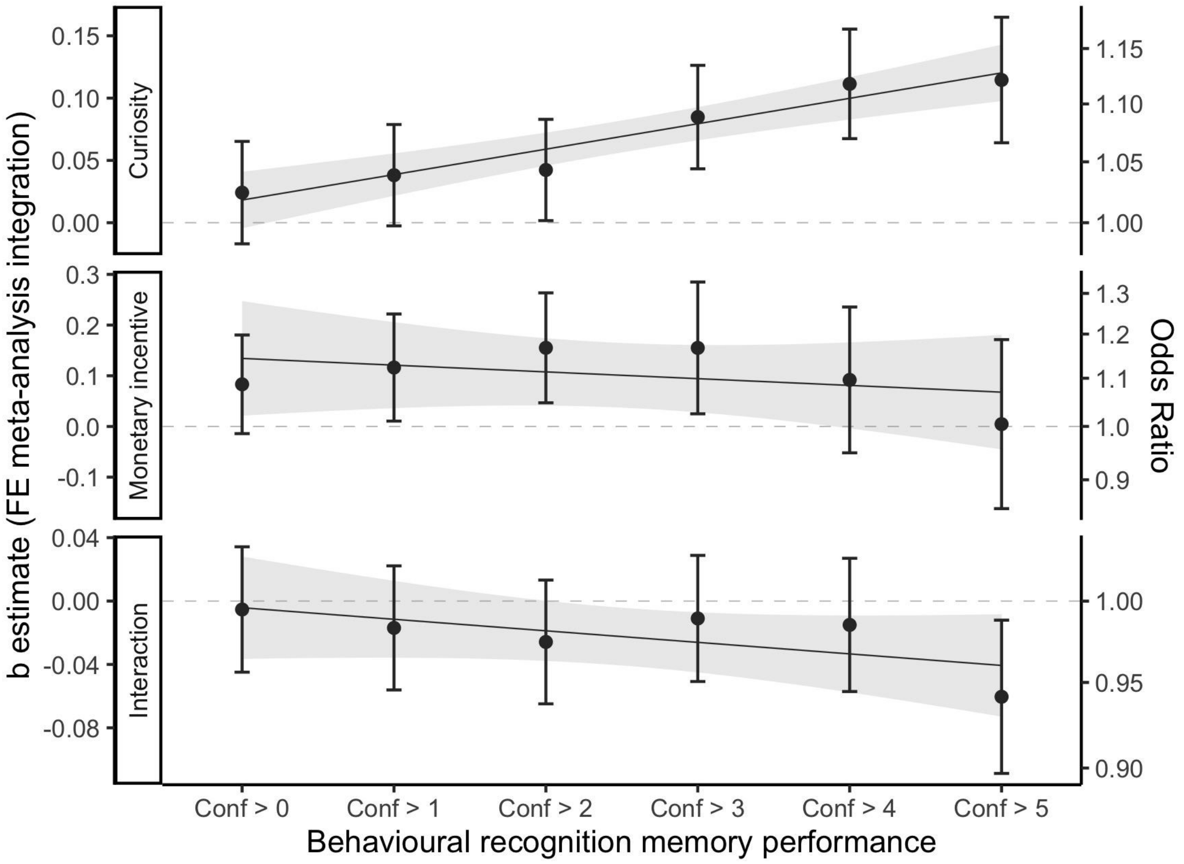

*Note*. The x-axis shows the gradual confidence cut-off and y-axis illustrates the integrated effect size (left - unstandardised, right - OR). Each panel shows one of the fixed effects specified in the gLME model. Importantly, a gLME model with reduced random effects structure (no random slope for the curiosity effect) was used to analyse the data from each data collection before integrating the results. The integrated b estimate for each effect and confidence threshold is plotted and error bars indicate 95%-CI. The regression line illustrates the linear model predicting the effect with the gradual confidence cut-off. This figure corresponds to Figure 3 in the main text where a gLME model with full random effects structure was specified to analyse the data before fixed effects meta-analysis integration.

**Figure S6**

*Fixed Effects Estimates Of Curiosity, Monetary Incentive, And Their Interaction As A Function Of Confidence Cut-off Separately For Each Data Collection*

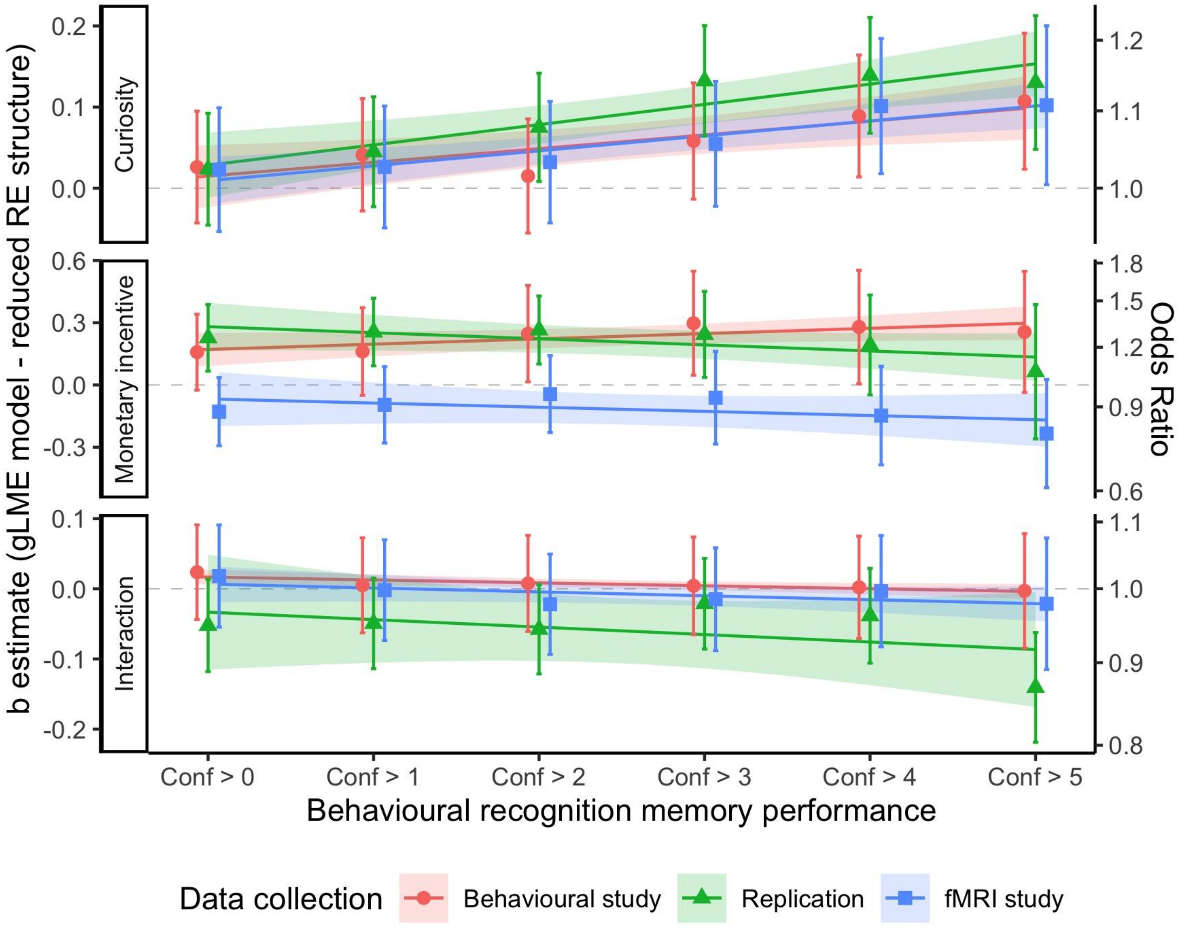

*Note*. The x-axis shows the gradual confidence cut-off and y-axis illustrates the integrated effect size (left - unstandardised, right - OR). Each panel shows one of the fixed effects specified in the gLME model. Importantly, a gLME model with reduced random effects structure (no random slope for the curiosity effect) was used. The b estimate from each data collection for each effect and confidence threshold is plotted and error bars indicate 95%-CI. The regression line illustrates the linear model predicting the effect with the gradual confidence cut-off. The data shown here was integrated and plotted in Figure S5 and this figure corresponds to Figure S4 where a gLME model with full random effects structure was specified.

**Figure S7**

*ISC In The ROIs*

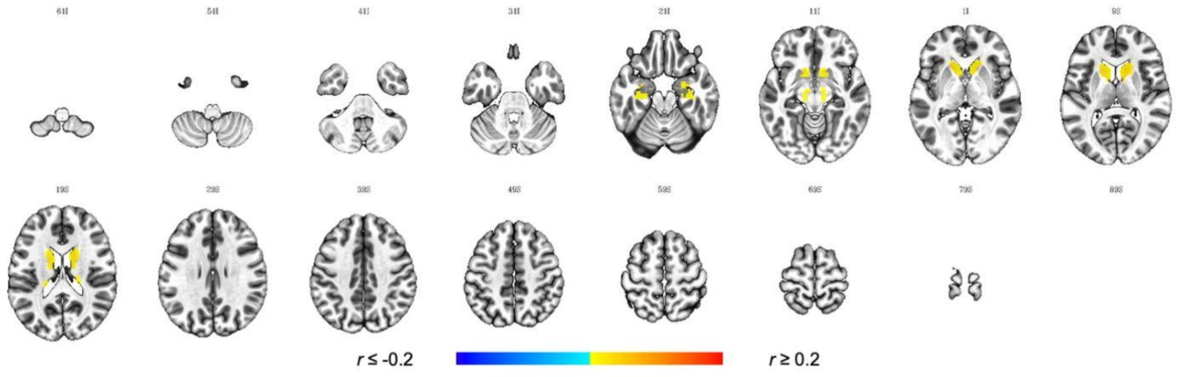

*Note*. Results are FDR-corrected at $q < 0.05$, cluster-extent corrected at $k = 5$ and plotted on the ICBM 2009c Nonlinear Asymmetric Template. Images are displayed in neurological orientation, where the left side of the brain is depicted on the left side of the image. ISC = Intersubject correlation.

**Figure S8**

*Schematic Illustration Of Intersubject Functional Connectivity (ISFC)*

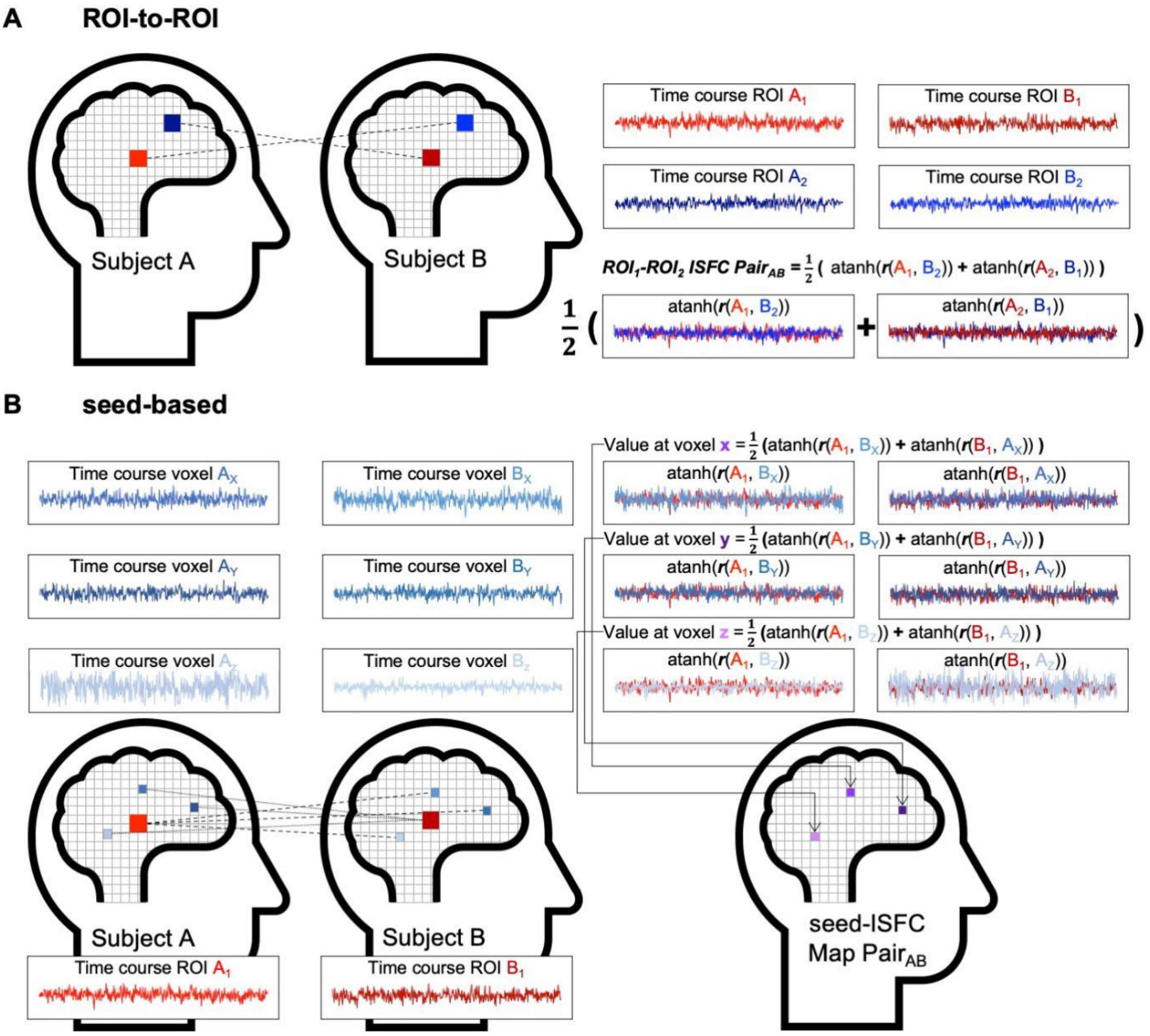

*Note.* Intersubject functional connectivity (ISFC) was computed as a measurement of consistency of responses between different regions. (A) To compute ISFC between ROIs, the averaged time course of ROI 1 was correlated with the averaged time course of ROI 2 between subjects for each pair of participants. The ROI-to-ROI ISFC for a given pair was determined as the mean of both correlations after Fisher's *z* transformation (indicated as *atanh()*). (B) To compute seed-based ISFC, the averaged ROI time course was correlated with the time course of all other voxels between subjects for each pair of subjects creating a seed-ISFC map for each subject of the pair. Both maps were averaged after Fisher's *z* transformation (indicated as *atanh()*) to determine the seed-ISFC map for the pair.

**Figure S9**

*ISFC And Incentive Effects Therein Specifying aHPC And VTA/SN As Seeds*

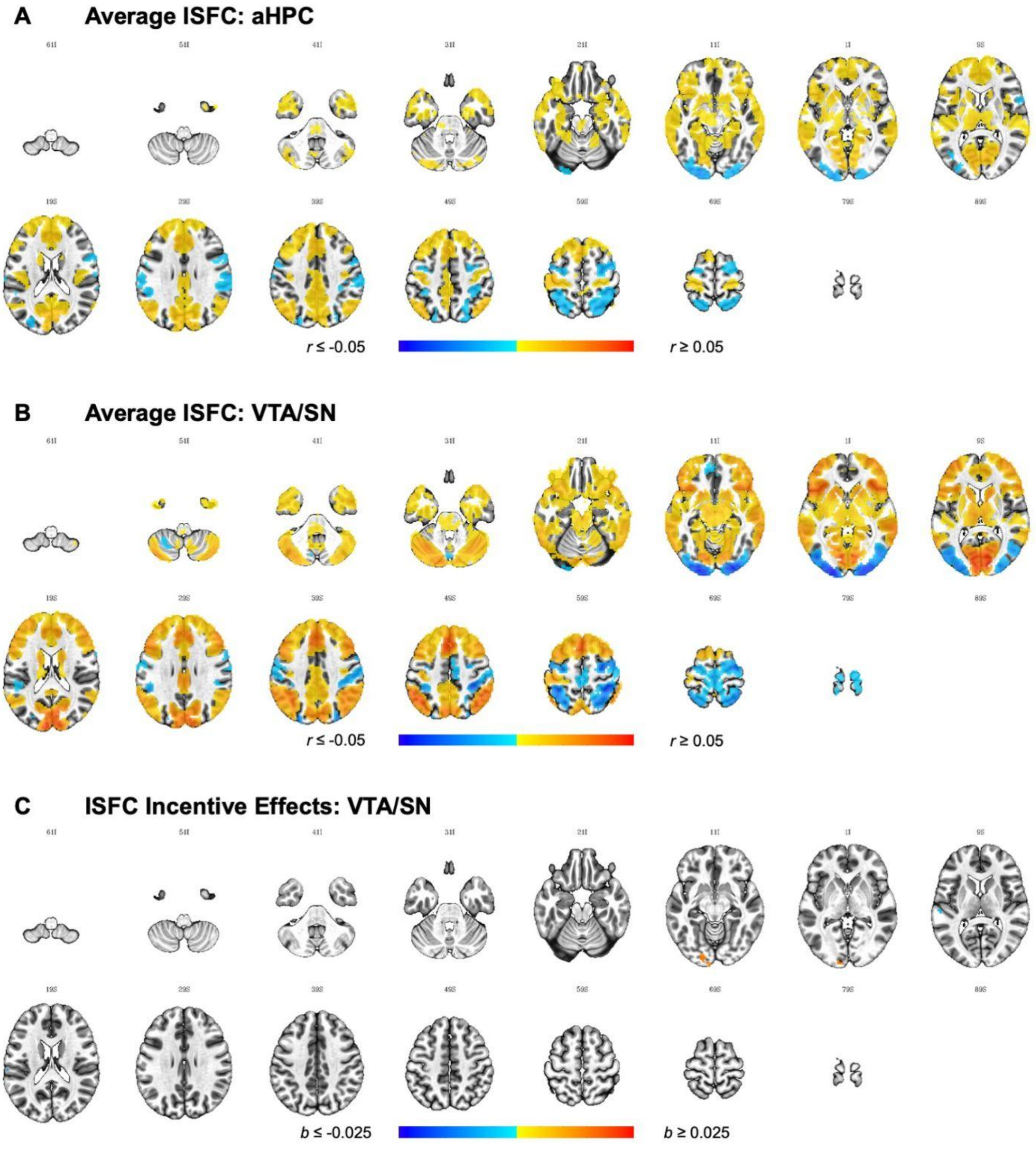

*Note*. Results are thresholded at $p < 0.001$, cluster-extent corrected at $k = 20$ (equivalent to per-cluster $\alpha = 0.05$) and plotted on the ICBM 2009c Nonlinear Asymmetric Template. Images are displayed in neurological orientation, where the left side of the brain is depicted on the left side of the image.

**Figure S10**

*ISFC-RSA For Each Behavioural Effect Of Interest Specifying aHPC And VTA/SN As Seeds*

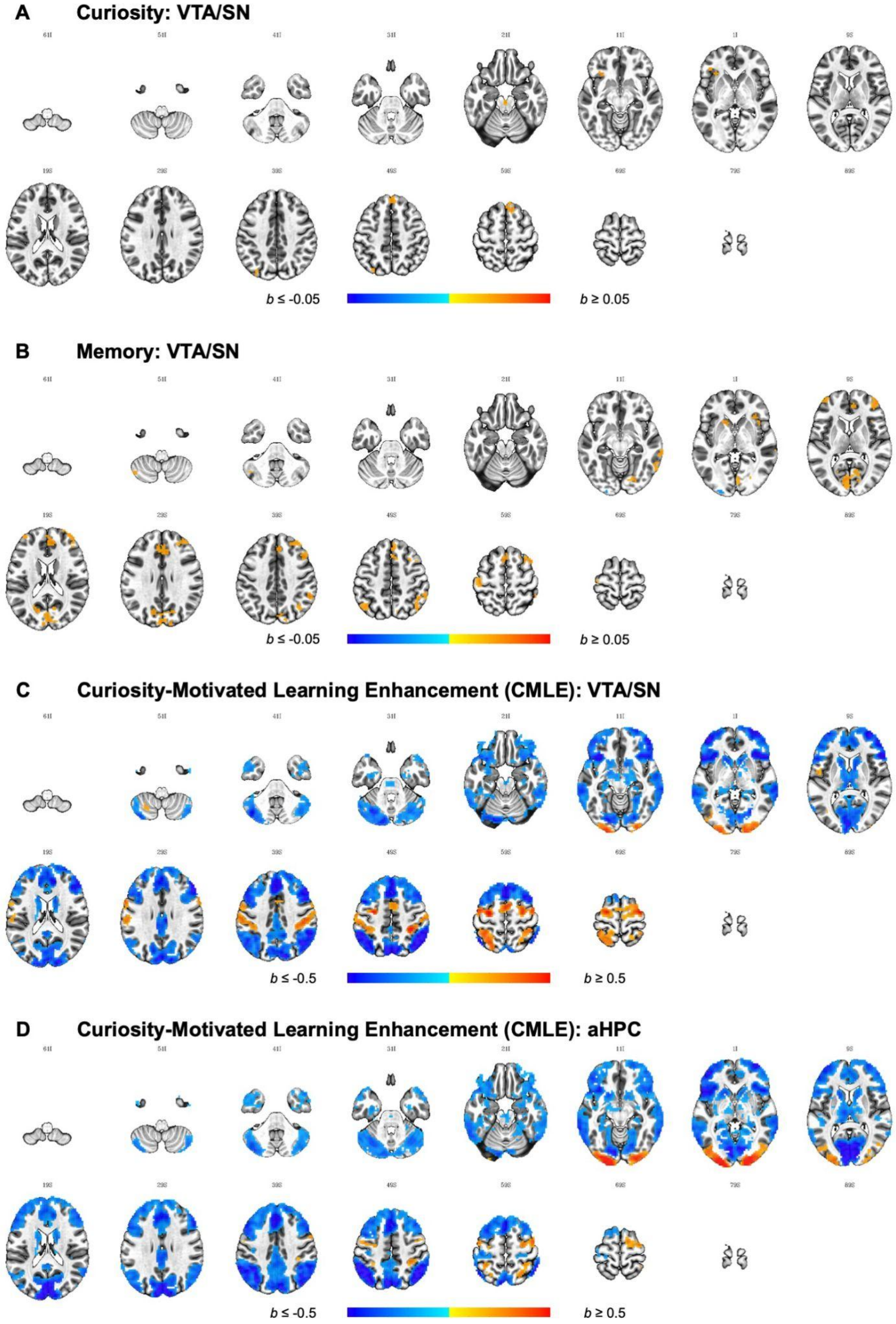

*Note*. Results are thresholded at $p < 0.001$, cluster-extent corrected at $k = 20$ (equivalent to per-cluster $\alpha = 0.05$) and plotted on the ICBM 2009c Nonlinear Asymmetric Template. Images are displayed in neurological orientation, where the left side of the brain is depicted on the left side of the image.

**Figure S11**

*ISFC-RSA For The Interaction Between The Incentive Manipulation And Each Behavioural Effect Of Interest Specifying aHPC And VTA/SN As Seeds*

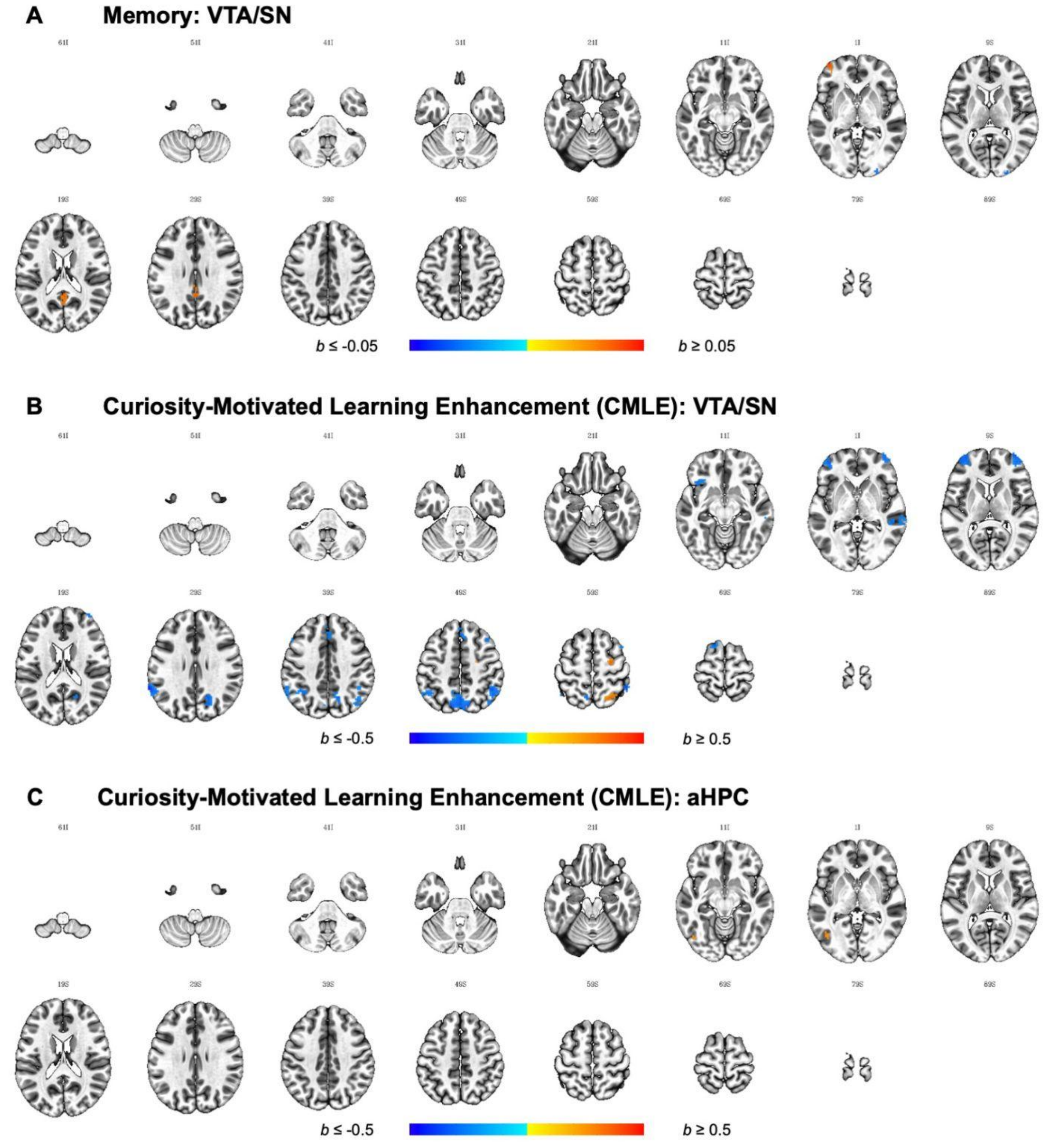

*Note*. Results are thresholded at $p < 0.001$, cluster-extent corrected at $k = 20$ (equivalent to per-cluster $\alpha = 0.05$) and plotted on the ICBM 2009c Nonlinear Asymmetric Template. Images are displayed in neurological orientation, where the left side of the brain is depicted on the left side of the image.

### Supplementary Analyses

#### Determination Of the Optimal Hemodynamic Response Function (HRF) Lag

Previous research has discussed different lags to account for the delay in the hemodynamic response function (HRF) (Hasson et al., 2004; Nastase et al., 2019; Zadbood et al., 2017), so there is no commonly applied correction to shift the time course by in the context of ISC. Here, intersubject pattern correlation (ISPC; Chen et al., 2017) - a data-driven approach was applied to determine the optimal lag (see Figure S2). In ISPC (also referred to as spatial ISC; Nastase et al., 2019), the correlation of a spatially distributed response pattern (e.g., within a searchlight or ROI) at a given time point is computed across subjects. Computing ISPC leads to a time point by time point correlation matrix where the diagonal captures the reliability of the spatial response across subjects (isolating the stimulus-driven component that is shared across subjects) at each moment in time and the off-diagonal values represent intersubject reinstatement of response patterns from time point t(i) at time point t(j) (Nastase et al., 2019). Because the time course of what was displayed on the screen is known, the reinstatement patterns of certain critical volumes can be used to validate which HRF lag is most appropriate. More specifically, after applying different HRF lags, the observed reinstatement patterns can be compared to expected reinstatement patterns: if a fixation was shown at consecutive time points t(i) and t(j), the intersubject reinstatement of t(i) at t(j) should be high. On the other hand, if a fixation volume at t(i) was not followed by another fixation volume at t(j), the intersubject reinstatement of t(i) at t(j) should be low.

To apply ISPC to our data, an initial concatenation step was carried out to reorder the volumes across subjects (Figure S2A). For each magic trick, four additional volumes before (1 TR fixation, 3 TRs mock video) and six additional volumes after the magic trick presentation (depending on jitter, 2-5 TRs fixation and 2-4TRs of rating, respectively) were selected (Figure S2B) and concatenated so that the final concatenated order of volumes remained invariant regardless of the pseudo-randomised order in which trials were presented (see Thomas et al., 2018). After the initial concatenation step, the time series consisted of 954 volumes. To determine the optimal HRF response lag, we were interested in the reliability and reinstatement of responses in the visual cortex (V2). The V2 mask was created based on the atlas by Glasser and colleagues (2016) of which the left and right second visual area were extracted before combining both and resampling them to EPI grid. The final V2 mask included 706 voxels. A leave-one-out approach was applied to compute ISPC where the response pattern in V2 in a subject at a single time point was correlated with the mean response pattern of all other subjects at all timepoints (Figure S2C, left). This procedure was repeated for all time points and subjects to create a 954 x 954 time point by time point correlation matrix for a given subject and the mean of all other subjects. The matrices were Fisher's *z*-transformed before computing the sample mean time point by time point correlation matrix (Figure S2C, right).

The $k^{th}$ upper and lower off-diagonals were extracted for $1 \leq k \leq 2$, hence values represent the intersubject reinstatement of the pattern observed at a given time point in the next volume or the after next volume. The upper off-diagonal shows the correlation between the subject response at volume t(i) and the mean response of all other subjects at volume t(i + k) whereas the lower off-diagonal shows the correlation between the mean response of all other subjects at volume t(i) and the subject response at volume t(i + k). In a next step (Figure S2D), each value extracted from the off-diagonal was labeled and categorised in correspondence to the selected volumes shown in Figure S2A. Additionally, the labels were shifted assuming different lags in the HRF response (1 volume ≤ HRF lag ≤ 6 volumes) and for each event category and HRF lag, data was averaged.

Analysis to determine the optimal lag focused on two events (outlined in black in Figure S2B): the fixation before the beginning of the video (one volume) and the first volume of fixation after the video. We predicted that the reinstatement of the fixation before the video ('pre fixation volume') should be small at k = 1 (first volume of the mock video) and even smaller at k = 2 (second volume of the mock video). For the first volume of fixation after the video ('post fixation volume'), we hypothesised a high reinstatement at k = 1 (second volume of fixation after the magic trick) decreasing at k = 2 (either third volume of fixation or first volume of rating). As shown in Figure S2sE, applying an HRF lag of 4 volumes when labelling the data best matches the predictions. Additionally, patterns in the upper and lower diagonal were largely similar ($r$ = .90).

**Intersubject Functional Connectivity (ISFC)**

Intersubject correlation (ISC) uses shared variance in the brain response between subjects to capture synchronicity of a given brain region inter-individually. Functional connectivity (FC; Biswal et al., 1995), on the other hand, computes the correlation of the seed regions (voxels or ROIs) within a subject to quantify synchronicity of different brain regions intra-individually.

Intersubject functional connectivity (ISFC; Simony et al., 2016) combines ISC and FC by using the time course of a seed region in one subject to correlate it with the time courses of all other regions or voxels in another subject's brain hence isolating shared, stimulus-evoked FC patterns (Vanderwal et al., 2019). While FC is prone to artefacts (e.g., head movement; Power et al., 2012) as well as stimulus-unrelated, spontaneous activity, ISFC isolates the stimulus-dependent inter-regional correlation between subjects because intrinsic neural dynamics and artefacts within a subject are not correlated across brains and hence provides an estimate for the temporal consistency of responses between regions (Nastase et al., 2019; Simony et al., 2016; Simony & Chang, 2020). Like FC, ISFC can be used to correlate the activity of a seed region with the activity in pre-defined ROIs (ROI-to-ROI ISFC, Figure S8A) or all other voxels in the brain (seed-based ISFC, Figure S8B).

We focused on anterior hippocampus (aHPC) and ventral tegmental area/substantia nigra (VTA/SN) as seeds because previous research showed that the two brain areas form a loop in the context of motivated learning where the encoding in the hippocampus is modulated by dopaminergic activity stemming from the midbrain (Lisman et al., 2011; Shohamy & Adcock, 2010). To compute ROI-to-ROI ISFC, i.e., the ISFC between aHPC and VTA/SN (aHPC-VTA/SN-ISFC thereafter), the averaged aHPC time course in subject A was correlated with the averaged VTA/SN time course in subject B. Likewise, the averaged VTA/SN time course of subject A was correlated with the averaged aHPC time course of subject B. To determine aHPC-VTA/SN-ISFC for participant pair A-B, the two correlation coefficients were averaged after Fisher's $z$ transformation. This procedure was repeated for all pairs of subjects and is illustrated in Figure S8A.

Firstly, we were interested in whether there was significant aHPC-VTA/SN-ISFC during the magic trick watching task, and whether this was influenced by the incentive manipulation. Data was analysed with LME-CRE models predicting aHPC-VTA/SN-ISFC with a fixed effect for group (deviation coded) accounting for the interrelatedness using crossed random intercepts for both subjects in each pair. Similar to ISC analysis, this approach cancels out any spontaneous response that is unrelated to the stimuli or any idiosyncratic patterns.

We also expanded the model to link idiosyncratic aHPC-VTA/SN-ISFC patterns to curiosity, memory, and curiosity-motivated learning enhancements (CMLE). A separate LME-CRE model was specified with crossed random intercepts for both subjects in each pair, where the incentive effect together with all three behavioural effects of interest were added as fixed effects into the same model. More specifically, the deviated coded group variable as well as grand-mean centred pairwise curiosity rating and memory encoding correlations as well as grand-mean centred pairwise mean values of the extracted individual curiosity beta values were used.

To examine the interaction between the incentive manipulation and the behavioural effects of interest, another LME-CRE model was specified to include the interaction terms between monetary incentives and each behavioural effect of interest as fixed effects in addition to the fixed effects described above. Interaction effects were computed by multiplying each main effect with the deviation coded incentive effect and all main and interaction effects were combined into one model. Like previous models, interrelatedness in the pairwise aHPC-VTA/SN-ISFC data was accounted for by defining crossed random intercepts for both subjects in each pair. All three LME-CRE models described here were applied using the *lme4* package in R (*'lmer()'*).

In addition to the ISFC between the aHPC and VTA/SN ROI, seed-based ISFC was conducted separately for both ROIs as seeds for exploratory purposes. The average time course of each seed in subject A was correlated with the (whole brain) voxel-wise time course subject B and vice versa (using *'3dTcorr1D'*, Figure S8B), creating two asymmetric matrices $r(x_A, y_B)$ and $r(x_B, y_A)$ that were averaged after Fisher's $z$ transformation (Nastase et al., 2019). The pair-averaged seed-based ISFC maps were analysed using LME-CRE models (using *'3dISC'*) akin to those used in ISC and IS-RSA predicting the Fisher's $z$-transformed pairwise ISFC maps for each seed (rather than Fisher's $z$-transformed pairwise ISC maps) to (a) determine overall seed-based ISFC of aHPC and VTA/SN, respectively, during magic trick watching, (b) identify incentive effects therein, and (c) behavioural effects of curiosity, memory, and CMLE and their interaction with incentives.

**ROI-to-ROI ISFC Between Anterior Hippocampus and VTA/SN.** Previous studies suggested that within-subject FC between aHPC and VTA/SN predicts curiosity- and reward-motivated learning (Gruber et al., 2014, 2016). Here, we were interested in between-subject FC between those two ROIs. More specifically, we investigated (a) whether the availability of monetary incentives influenced the strength of aHPC-VTA/SN-ISFC, and (b) whether the behavioural similarity matrices could predict aHPC-VTA/SN-ISFC and whether this was further affected by the incentive manipulation.

To determine whether the aHPC-VTA/SN-ISFC differed between control and incentive group, a LME-CRE model was specified using the availability of monetary incentives to predict aHPC-VTA/SN-ISFC. There are two observations from the results (Table S11). First, the intercept was positive and significant, meaning that there is, in fact, systematic stimulus-driven communication between VTA/SN and aHPC activity across subjects when watching magic tricks. Second, the incentive effect was not significant, suggesting that the availability of monetary incentives did not affect the ISFC between VTA/SN and aHPC.

In a next step, we examined whether the behavioural effects of interest could predict aHPC-VTA/SN-ISFC. All three behavioural effects of interest were entered as covariates into the same LME-CRE model; however, no statistically significant effects were found and only the main effect of curiosity reached trend level (see Table S11).

To explore whether the incentive manipulation influenced the relationship between the behavioural effects of interest and aHPC-VTA/SN-ISFC, a last LME-CRE model was specified to include the interaction effects together with the main effects of incentive and behavioural covariates. As indicated in Table S11, no effects reached statistical significance, despite the fact that the curiosity effect remained at trend level.

**Seed-Based ISFC Of Anterior Hippocampus And VTA/SN.** In addition to the ISFC between aHPC and VTA/SN, their respective whole brain ISFC was examined by specifying them as seeds creating seed-based ISFC maps where each voxel value represents the between-subject correlation of that voxel's time course with the averaged time course of the seed region. To identify inter-regional correlations of aHPC and VTA/SN when exposed to magic tricks, LME-CRE models were used. These analyses revealed large clusters where shared information about the magic trick stimuli is encoded in a similar manner as in the respective seeds (Figure S9A and S9B, Table S12), suggesting that they encode shared information about the magic trick stimuli. Overall, maps for both seeds were highly similar (correlation unthresholded effect size map = .855, correlation unthresholded statistics map = .819, dice coefficient of masked cluster-extend thresholded results = .627), but ISFC for the VTA/SN seed is spread further and values are numerically higher compared to the aHPC seed. More specifically, using the 7-Network parcellation proposed by Yeo and colleagues (2011) as reference, cortically, the seeds show positive ISFC with the medial Visual (Vis), Somatosensory, Limbic, Default Mode (DMN), Frontoparietal (FPN), and Ventral Attention (VAN) networks whereas negative clusters were located in lateral part of the Vis and the Dorsal Attention network (DAN). Positive clusters were further found in the striatum, midbrain, medial temporal lobe, and thalamus. While the incentive manipulation did not affect ISFC patterns of the aHPC seed, two clusters were found for the VTA/SN seed (Figure S9C, Table S12). More specifically, a cluster in the Vis (left occipital pole stretching into inferior occipital gyrus laterally) was found where the VTA/SN seed ISFC was more negative in the incentive compared to the control group, whereas ISFC values were larger in the incentive compared to the control group in the left VAN, more specifically in the left superior temporal gyrus.

To determine brain regions where similarity in the behavioural effects of interest covaries with similarity in the seed-based connectivity with aHPC and VTA/SN, the IS-RSA framework was extended to predict pairwise seed-based ISFC maps rather than the pairwise ISC maps. Separate LME-CRE models were run for each effect and each seed and the detailed results can be found in the supplementary material (for main effects, see Figure Figure S10 and Table S13; for interaction effects, see Figure S11 and Table S14).

For the aHPC seed, no main effects of curiosity or memory were found. When trying to relate the behavioural effects of interest to VTA/SN ISFC, on the other hand, clusters could be found for all three variables. Similarity in curiosity predicted similarity in the VTA/SN FC in the bilateral superior medial gyrus (BA 8), left AIC, IPL (BA 39), and the dorsal pons. Similarity in memory could positively predict similarity in FC with the VTA/SN in the bilateral medial visual cortex, bilateral frontal cortex (BA 9, BA 46, BA 9-46, BA 6, BA 8), anterior cingulate cortex, left somatosensory cortex, bilateral angular gyrus (BA 39), supramarginal gyrus (BA 40), right AIC, right middle (BA 21) and inferior temporal gyrus (BA 37), the left CN, and left cerebellum. Additionally, in one cluster in the left occipital pole, similarity in memory negatively predicted similarity in FC with the VTA/SN.

Looking at how effects of similarity in CMLE predict similarity in ISFC with both seeds, respectively, the resulting maps for aHPC (6 positive and 4 negative clusters) and VTA/SN (10 positive and 7 negative clusters) again looked very similar (correlation unthresholded effect size map = .890, correlation unthresholded statistics map = .888, dice coefficient of masked cluster-extend thresholded results = .754). For both seeds, positive clusters were predominantly located in the DAN and Vis (the latter laterally) whereas negative clusters were found in medial areas of the Vis, DMN, FPN, Limbic, and VAN. Intriguingly, the maps found for the CMLE ISFC-RSA effects appeared like a ‘flipped’ version of the ISFC obtained during magic trick watching (correlation unthresholded effect size maps $\leq$ -.850, correlation unthresholded statistics map $\leq$ -.811, dice coefficient of masked cluster-extend thresholded results $\geq$ .616).

When investigating whether the availability of monetary incentives influenced the association between similarity in behavioural effects of interest and VTA/SN and aHPC

ISFC, respectively, no clusters were found for curiosity in either seed. Likewise, no clusters were found for the memory incentive interaction for the aHPC seed. For at the VTA/SN seed, however, in total 3 clusters were found: one cluster in the right occipital pole was found where the predictive effect was larger in the incentive compared to the control group. For the opposite effect, two clusters located in the posterior cingulate cortex (PCC) and left dlPFC were identified. With respect to CMLE, one cluster was found for the aHPC seed where values were more positive in the control compared to the incentive group. For the incentive CMLE interaction on VTA/SN seed-based ISFC, in total 16 clusters were found. The two positive clusters (control > incentive) were located in the right superior parietal lobe and right superior frontal gyrus. The 14 negative (incentive > control) clusters were located in the superior parietal lobe, right precuneus, bilateral MFG, bilateral inferior parietal lobe, left supramarginal gyrus, right middle temporal gyrus, the superior medial gyrus, left superior frontal gyrus, and left anterior insula/frontal operculum.

### Supplementary Discussion

#### Curiosity-Motivated Learning Outside The Reward-Related Areas And The Hippocampus

In addition to the results within the *a priori* ROIs (the reward-related areas and the hippocampus), our whole brain IS-RSA showed the broad network of the brain supporting curiosity, memory, and curiosity-motivated learning enhancement (CMLE). To better understand the IS-RSA results, the resulting clusters for each effect of interest were compared with the 7-Network parcellation proposed by Yeo and colleagues (2011) dividing the brain into Visual (Vis), Somatosensory, Dorsal Attention (DAN), Ventral Attention (VAN), Limbic, Frontoparietal (FPN), and Default Mode (DMN) networks.

**Curiosity.** With respect to the effects of curiosity elicitation, we found that similarity in the curiosity ratings predicted similarity in the brain response in visual areas, the inferior frontal gyrus (IFG), the supplementary motor area (SMA), the postcentral gyrus, precuneus, anterior insula, and the supramarginal gyrus. The elicitation of curiosity has repeatedly been linked to a state of uncertainty, potentially due to a violation of expectations (Gruber & Ranganath, 2019; Murayama et al., 2019). Such violations of expectations have previously been linked to - amongst other regions - dlPFC, premotor cortex, posterior parietal cortex (PPC), and ventral visual stream (Murty & Adcock, 2014). In alignment with this proposal, our findings show that the IS-RSA effect of curiosity is located in the SMA, an area that has been implicated in the processing of uncertainty (Cheung et al., 2019; Volz et al., 2005) and in the IFG - part of the lateral PFC. The IFG has been linked to the elicitation of curiosity in the trivia question paradigm (Gruber et al., 2014; Kang et al., 2009). According to the Prediction, Appraisal, Curiosity, and Exploration (PACE) framework explaining how curiosity enhances HPC-dependent memory (Gruber & Ranganath, 2019), the IFG is involved in the appraisal processes determining whether PE and associated uncertainty elicits curiosity or anxiety. The IFG has also been linked to the violation of expectations (Danek et al., 2015) and causal relationships (Parris et al., 2009) in magic tricks. This suggests that as participants watch magic tricks, the curiosity IS-RSA effect in the SMA and the IFG could reflect that uncertainty-related signals and their appraisal processes of the experienced prediction errors share a similar signature when experienced curiosity is similar.

Curiosity IS-RSA effects were also observed in the posterior parietal cortex (PPC) - a region anteriorly responding to motoric and perceptual representations of actions and candidate for mirror neuron system areas (Chong et al., 2008; Dinstein et al., 2007; Ishida et al., 2010). This could reflect attentional processes. The dorsal part of the PPC (lateral and medial parts of BA7) has been implicated in top-down attention together with dorsal frontal regions, whereas the ventral part (corresponding to BA 39 and BA 40, often referred to as inferior parietal lobe (IPL)) together with ventral frontal regions are involved in bottom-up attentional processes (Corbetta et al., 2008; Corbetta & Shulman, 2002). Because IS-RSA curiosity effects were located in dorsal (i.e., precuneus) and ventral (i.e., right supramarginal gyrus) PPC regions, as well as in the IFG - part of the wider ventral attention system - this suggests that curiosity is associated with top-down goal-directed attention related to the judgement task or the re-direction of attention in response to the ventral system signalling the violation of expectations and causal relationships as salient events in a bottom-up manner. Indeed, the PPC and more specifically, the IPL has previously been linked to signalling the moment of expectation violation in magic tricks (Danek et al., 2015), but also to signalling surprise and model update thereafter in other tasks (O'Reilly et al., 2013). Overall, these attentional mechanisms could further be related to uncertainty underlying curiosity and exploration and information-seeking to resolve it. In fact, investigating the effect of curiosity on eye movements, Baranes and colleagues (2015) suggested that curiosity is associated with prioritisation and allocation of attention which could be supported by 'priority maps' in the parietal cortex (Bisley & Goldberg, 2010). The induction of curiosity has also been linked to increased activity in the IPL in a lottery task (van Lieshout et al., 2018) as well as within the trivia question paradigm (Duan et al., 2020; Ligneul et al., 2018), but also more broadly to knowledge uncertainty (Volz et al., 2004). Overall, this suggests that, when the state of curiosity is high, people tend to show shared activity of areas related to (re-)directing attention and processing uncertainty triggered by salient events that violate predictions — people in a curious state may have similar time courses of attention (re-)direction and experience of uncertainty.

Importantly, while initial fMRI research using the trivia question paradigm suggested that the elicitation of curiosity is supported in dopaminergic regions in the striatum and midbrain (Gruber et al., 2014; Kang et al., 2009), later studies failed to replicate this effect and instead found striatal activity at curiosity relief (Duan et al., 2020; Ligneul et al., 2018). While these studies differed in various aspects from another (e.g., intentional vs. incidental encoding; general knowledge- vs. cinema-related trivia questions), the latter studies included a jittered period between cue and target presentation whereas the initial studies did not. As such, the elicitation of curiosity is confounded with the anticipation of rewarding information. Hence, it is possible that the dopaminergic brain activity found was due to the anticipation of rewarding information rather than due to the elicitation of curiosity *per se*. Indeed, in the context of fully-predictable extrinsic rewards, it has been shown that dopaminergic neurons have signalled the reward-predicting stimuli rather than the reward itself (Knutson et al., 2001; Schultz, 1998; Tobler et al., 2005), creating blunt prediction error signals (O’Doherty et al., 2003). In a similar manner, if perceptual curiosity is relieved in only half of the trials, striatal activity is found at relief and not at elicitation (Jepma et al., 2012). As such, results by Jepma et al. (2012) and Ligneul et al. (2018) show that if curiosity is relieved in a stochastic manner, prediction error signals are observed during relief. Taken together, this suggests that when the elicitation of curiosity is not confounded with the anticipation of rewarding information, the curiosity-related effects can be found during curiosity relief and not as previously suggested merely during elicitation. Because in our paradigm, curiosity was elicited, but never relieved and no rewarding information was anticipated, this could potentially further explain the absence of curiosity effects in reward-related structures.

**Memory.** It has previously been proposed that in addition to the medial temporal lobe (MTL), cortical systems or networks support memory encoding (Bastin et al., 2019; Fuster, 1997; Kim, 2011; Ranganath & Ritchey, 2012; Spaniol et al., 2009). As such, different components in the encoding process have been proposed (Kim, 2011): storage, content processing, and attention. While the MTL is implicated in the storage function (Squire et al., 2004), other brain areas mediate the other components. Critically, while the IS-RSA effects during the incidental encoding of magic tricks were not located in the classical storage regions within the MTL, we found clusters in areas of the brain involved in other components of the memory process — content processing and attention. For example, we observed significant IS-RSA memory effects in IFG and fusiform gyrus. Evidence suggests that the lateral PFC is involved in control processes where ventrolateral regions select goal-relevant item information and dorsolateral regions support the organisation of information within working memory to form associations (Blumenfeld & Ranganath, 2007). Also, other IS-RSA memory clusters in the parietal cortex are located in more posterior regions, supporting a top-down and bottom-up attentional control (Shomstein, 2012). An attention-driven involvement of the PPC in memory has predominantly been discussed in the context of retrieval (for reviews, see Cabeza et al., 2008; Shimamura, 2011), potentially explaining why the PPC is more activated during recollection- compared to familiarity-based recognition (Kim, 2010) and is further involved in autobiographical memory retrieval (Cabeza et al., 2004).

These findings are consistent with the idea that memory is not solely supported by the MTL, but rather memory output is a consequence of complex interaction of different mental processes. For example, the hierarchy of process memory framework (Hasson et al., 2015) posits that each cortical circuit along the processing hierarchy is able to accumulate information over time, but the temporal receptive windows (time spans in which prior information can impact current processing) increase as information travels from sensory to high-order cognitive regions. This suggests that memory of recent events (for instance, the events of the magic trick that is currently viewed) are not stored in dedicated areas in e.g., the MTL, but are organised in a hierarchical manner across the cortical regions processing the information, enabling the online processing of continuous, dynamic, complex stimuli by retaining information independent of MTL involvement. The widespread cortical IS-RSA effects with the visual cortex in its centre suggest that as participants view the magic tricks, neurons process and retain the information within their respective temporal receptive windows at each level of hierarchy which in turn also supports later memory for the magic trick.

**Curiosity-Motivated Learning Enhancement (CMLE).** Significant CMLE effects were observed in broad cortical areas including large parts of the DMN (e.g., bilateral ACC, angular gyrus, middle temporal gyrus), FPN (e.g., bilateral MFG, SMA), DAN (e.g., bilateral posterior superior parietal lobe), VAN (e.g., anterior insula/frontal operculum (aI/fO)) as well as Vis. This replicates results by others showing that the effect of curiosity on memory is reflected in distributed brain regions including, e.g., the FPN (Duan et al., 2020; Murphy et al., 2021). Importantly, these effects were negative, suggesting that participants with high CMLE scores showed more individualised responses to the magic trick stimuli compared to participants with low CMLE scores.

These observations have various implications. For example, it has been shown that during naturalistic viewing the anterior (e.g., medial PFC) and lateral DMN (e.g., angular gyrus) are strongly intrinsic whereas the preceneus shows more extrinsic responses that correlate between subjects, making the DMN a prime candidate to integrate extrinsic, stimulus-driven activity with internal processes due to the interconnectivity between these regions (Ren et al., 2017). A recent perspective proposes that the DMN is highly implicated in the processing of the dynamic structure of external, naturalistic stimuli, regardless of modality, by integrating inputs across longer temporal receptive windows with idiosyncratic, internal prior dispositions (e.g., prior knowledge or beliefs) to form models of the experienced situation (Yeshurun et al., 2021). This perspective can explain the role of the DMN in curiosity-motivated learning: In the context of the trivia question paradigm, the involvement of the DMN has been discussed in light of the successful accumulation and integration of new information into prior knowledge and schemas (Murphy et al., 2021), however, the DMN has also been implicated in processing surprise in naturalistic viewing due to its ability to detect the mismatch between incoming information and internal models (Brandman et al., 2021). Any internal models shaped by prior experiences could be supported by unique firing patterns within the brain and communication across brain regions, hence explaining why curiosity-motivated learning is accompanied by endogenous responses within the DMN, further supporting the update of internal models in the context of curiosity-motivated learning.

Regarding the negative cluster in the FPN, previous work has argued that the FPN exerts cognitive control, especially in the context of multiple demands, allowing the brain to remain flexible and adaptive (Dosenbach et al., 2007). A potential mechanism to achieve cognitive control is to maintain task-relevant information and suppress irrelevant information. In naturalistic conditions, the FPN, showing high variability in FC, may be involved in shifts in the broader processing of stimuli (Vanderwal et al., 2017). Our findings that participants with high CMLE scores are less similar in their stimulus-induced brain response in the FPN compared to low scorers sheds further light on the mechanisms by which the FPN is involved in the enhancing effects of curiosity on memory reported by others (Duan et al., 2020; Murphy et al., 2021): the FPN de-synchronises across participants suggesting individualised, internal processes by which the FPN - previously described as flexible hub (Cole et al., 2013) - supports curiosity-motivated learning, potentially by integrating external sensory information with internal representations by transferring information between DMN and DAN, especially in the context of stimulus-related conflict (Vincent et al., 2008) as caused by the violation of cause and effect relationships in magic tricks.

**The Surprising Role Of The Anterior Insula**

As a surprising result, we found that IS-RSA effects of curiosity, memory, and CMLE all anchored their similarity onto similarity in the right anterior insula/frontal operculum (aI/fO) with clusters overlapping in the right anterior insula cortex (AIC). The AIC is part of the putative ‘salience network’ (Seeley et al., 2007), but also of the cingulo-opercular task maintenance network (Dosenbach et al., 2007) suggesting the AIC to be involved in both, tonic and phasic alertness supporting the detection of salient as well as homeostatically relevant signals and their integration into awareness (Craig, 2009) and has been found to be co-activated in thousands of studies (Chang et al., 2012; Craig, 2009; Droutman et al., 2015; Seeley, 2019; Yarkoni et al., 2011). As such, the AIC has been linked to urge generation and addiction (Naqvi & Bechara, 2010), as well as attention, cognitive control and executive functioning (Dosenbach et al., 2006; Mayer et al., 2007; Sridharan et al., 2008). The AIC is also associated with the processing of reward, punishment and subjective value of choice alternatives regardless of their valence (Bartra et al., 2013) and is recruited during the anticipation of rewards (Diekhof et al., 2012; Liu et al., 2011; Wilson et al., 2018) where the

AIC might play a specific role in reward-based attention (Wang et al., 2015) or the cognitive expectations expressing causal relations regarding act-outcome representations of incentive value (Berridge, 2000). Investigating the basis of motivational decision-making under uncertainty in the field of neuroeconomics (reviewed by Platt & Huettel, 2008), research has observed increased activity in the AIC associated with risky decision-making (Paulus et al., 2003), risk prediction and risk prediction errors (Preuschoff et al., 2008). Likewise, the AIC has been implicated in error processing (Ullsperger et al., 2010), information update (van Lieshout et al., 2018), prediction errors (Weilnhammer et al., 2017) and their magnitude (Pine et al., 2018), the violation of expectations (Danek et al., 2015; Schiffer & Schubotz, 2011), uncertainty (Grinband et al., 2006; Huettel et al., 2005, 2006; Volz et al., 2003), and surprise (Loued-Khenissi et al., 2020), but also the elicitation (Jepma et al., 2012) and relief of curiosity (Lee & Reeve, 2017). Moreover, the AIC not only signals the magnitude of unsigned prediction errors but is further involved in memory updates thereafter (Pine et al., 2018). Pre-stimulus activity in the AIC before the onset of movie clips has been shown to positively predict the encoding thereof and further influences memory performance by increased activity in temporal regions, but decreased activity in the posterior precuneus, cingulate, and striatum during stimulus presentation (Cohen et al., 2020). The AIC has further been implicated in the subsequent memory effects (Kim, 2011) in general, and curiosity-motivated learning in particular (Duan et al., 2020).

On a broader level, curiosity is often seen as an element or a source of intrinsic motivation (Deci & Ryan, 1985). In an attempt to identify the neural mechanisms of intrinsic motivation, the AIC has been discussed as a basis thereof (Di Domenico & Ryan, 2017; Lee, 2016). As such, the AIC has been found to be more activated when participants imagined doing a task out of intrinsic rather than extrinsic reasons (Lee et al., 2012; Lee & Reeve, 2013). Likewise, higher activity was found when participants performed intrinsically motivating tasks compared to non-intrinsically motivating tasks, and activity in the AIC not only predicted trial-level interest but also interacted with the striatum, thereby potentially forming the intrinsic motivation system by combining intrinsic reward processing in the striatum with feelings of inherent satisfaction in the AIC (Lee & Reeve, 2017).

In another account linking the AIC to intrinsic motivation, Di Domenico & Ryan (2017) discussed the AIC as the hub in the bilateral salience network (Menon & Uddin, 2010) involved in the bottom-up detection of subjectively important events to support goal-directed behaviour by providing attentional resources and flexible control. Importantly, because the overlap here is found on the right hemisphere during encoding, the activity observed here could also reflect the right-lateralized ventral attention network described by Corbetta and colleagues (2008) - a bottom-up ‘alerting system’ or ‘circuit breaker’ to shift attention in the top-down DAN. More specifically, the AIC plays a central role in switching between default mode and central executive networks (Sridharan et al., 2008) as well as the DAN (Huang et al., 2021), potentially making the AIC a candidate to represent a gateway for sensory information to enter consciousness, together with other structures like the thalamus.

Furthermore, the AIC has anatomical and functional connections with the HPC (Fanselow & Dong, 2010) as well as the dopaminergic striatum (Chikama et al., 1997; Cho et al., 2013; Flynn, 1999). Dopamine depletion is associated with reduced connectivity between the salience network and other parts of the brain (Shafiei et al., 2019), altogether suggesting that the AIC is innervated by dopamine and modulated. Importantly, dopamine does not only signal reward, but in addition to value-coding dopamine neurons, there are also salience-coding dopamine neurons (Bromberg-Martin et al., 2010). Additionally, there is also evidence for noradrenergic effects in the AIC, not only because noradrenaline is associated with salience (Totah et al., 2019), but ISC in the salience network is dependent on noradrenaline and pharmacologically blocking it during movie watching decreased ISC in the salience network including the aI/fO (Hermans et al., 2011). Noradrenaline has in fact been linked to the reward effect on memory, potentially mediated via surprise (Hauser et al., 2019). On the other hand, surprise during naturalistic viewing has also been linked to dopaminergic activity (Antony et al., 2021). Likewise, while the role of dopamine in curiosity has long been discussed, studies suggest that noradrenaline is additionally involved in curiosity (Sakaki et al., 2018). Traditionally, it was assumed that dopamine is released in the VTA/SN and noradrenaline in the locus coeruleus, however, recent evidence suggests that catecholamine pluripotent neurons in the LC also release dopamine affecting hippocampal consolidation processes (Kempadoo et al., 2016; Takeuchi et al., 2016). Overall, this suggests that both dopamine and noradrenaline could be important in the context of curiosity and curiosity-motivated learning, making the AIC a potential candidate in signalling the effects of both neurotransmitters.

How can this rich and diverse literature on the AIC be integrated theoretically? Sterzer and Kleinschmidt (2010) proposed a framework according to which the AIC plays a central role in perception: AIC activity represents the recruitment of processing resources during ‘challenging’ sensory stimulation to mediate states of sensitivity and reactivity to the environment through reciprocal connections with all sensory cortices. The challenge may arise in a bottom-up manner related to salient aspects of the stimuli or in a top-down manner due to task demands requiring cognitive effort as found in the context of uncertainty or ambiguity. Watching magic tricks whilst performing a judgement task is likely to recruit processing resources in a top-down as well as bottom-up manner. Especially in the context of ISC analysis, it has been suggested that certain brain areas - amongst them the AIC - are more likely to represent information that has been derived from the stimuli (rather than the stimuli themselves) and is hence more idiosyncratic across individuals (Ren et al., 2017). Using IS-RSA, we were able to link variations in the stimulus-induced response in the AIC to fluctuations in curiosity, memory, and CMLE. Hence, our results showing that similarities in our behavioural effects of interest are reflected by similarities in the stimulus-evoked responses in the AIC could reflect recruitment of cognitive resources during their naturalistic viewing that can potentially have an impact on all behavioural effects of interest.

Moving beyond perception and extending computational models of the AIC on risk-taking, Singer and colleagues (2009) postulated a unifying model of various fMRI evidence linking the AIC to feelings, empathy, and the processing of uncertainty in decision-making. According to their model, the AIC signals representations about predicted feeling states, current feeling states, and feeling state prediction errors that in combination support error-based learning, especially in the context of uncertainty. Information about these states is then integrated into a dominant subjective feeling state modulated by individual characteristics (e.g., sensation seeking) as well as contextual appraisal processes. Together, these mechanisms can not only motivate behaviour and guide decision-making in complex and uncertain environments, but also facilitate learning. Applying this framework to curiosity-motivated learning where curiosity can be conceptualised as a state of uncertainty (Gruber & Ranganath, 2019), the AIC might support representations of the predicted state of knowledge, the actual state of knowledge, and knowledge state prediction errors in light of that uncertainty where the latter is used to inform future predicted knowledge states to reduce uncertainty in the future, hence functioning as learning.

A recent fMRI study (Ligneul et al., 2018) further supported the idea that curiosity and curiosity-motivated learning might be better explained within the framework of predictive coding and uncertainty than using reward-maximisation. Predictive coding theories (Friston, 2010; Friston et al., 2012) propose that organisms strive towards minimising uncertainty and surprise levels associated with sensory inputs by optimising the internal generative model of the environment. However, this maxim is somewhat conflicting with curiosity-related behaviour that might lead to transient increases in uncertainty (Schwartenbeck et al., 2013). This is why second-order expectations regarding an optimal amount of uncertainty have been proposed (Clark, 2013) functioning homeostatically to maintain uncertainty around an expected value. Trying to link these concepts to curiosity and curiosity-motivated learning, Ligneul and colleagues (2018) tested the hypothesis of whether a large (or low) estimate of average surprise (and thus uncertainty) experienced in a given context will down- (or up-) regulate situational epistemic curiosity levels. Indeed, using a modified version of the trivia question paradigm, behavioural data showed an inverse relationship between average surprise triggered in previous trials and epistemic curiosity ratings. Analysis of the fMRI data indicated that average surprise was monitored and updated during answer presentation by deactivation in the rostrolateral PFC - a region previously linked to uncertainty-driven exploration - to regulate (i.e., reduce) upcoming curiosity levels. Likewise, during question presentation, high average surprise levels reduced the amplitude of responses in the salience (e.g., AIC) and executive control network (e.g., dlPFC, IPL) that might potentially signal post-processing activity or depletion of attentional resources according to the authors. Although some have argued that curiosity is a reward-seeking process where ‘information is reward’ (Marvin & Shohamy, 2016), the predictive coding framework tested by Ligneul and colleagues (2018) characterises curiosity in terms of uncertainty. As such, high states of curiosity during the question presentation would not reflect the anticipation of more pleasurable outcomes, but the experience of more uncertainty. Likewise, activity during the answer presentation would be interpreted as relief- and/or surprise-related rather than in terms of reward processing. The authors further argue that stochastic relief of curiosity might be associated with a dopaminergic response whereas average surprise levels are likely to be signalled by noradrenaline which is well suited to mediate the interplay between surprise and memory. Intriguingly, both noradrenaline

(Lawson et al., 2021) as well as dopamine (Gershman & Uchida, 2019) have not only been discussed in the context of signalling uncertainty, but also in respect to curiosity-motivated learning (Sakaki et al., 2018).

If uncertainty is a key driver behind curiosity as indicated in previous fMRI studies as well as our findings (Duan et al., 2020; Jepma et al., 2012; Ligneul et al., 2018; van Lieshout et al., 2018), could curiosity-motivated learning also be better characterised in terms of learning under uncertainty than as reward-based learning? To answer this question, more research is needed. Information reducing uncertainty is likely associated with a dopaminergic neural response (Duan et al., 2020; Jepma et al., 2012; Ligneul et al., 2018). However, as discussed elsewhere (Ligneul et al., 2018), such a dopaminergic response in and of itself cannot be interpreted as evidence favouring reward over predictive coding frameworks. Indeed, recent results (Duan et al., 2020; Murphy et al., 2021) have linked curiosity-motivated learning to brain areas associated with the update of internal models of the environment (e.g., anterior cingulate cortex and precuneus), further supporting the idea that curiosity-motivated learning can (partly) be explained using the predictive coding framework. Because the AIC is not only linked to uncertainty through lack of knowledge but also associated subjective feelings (Singer et al., 2009), it might be a promising candidate in signaling curiosity and supporting curiosity-motivated learning.

While more research is necessary to fully understand the role of the AIC in curiosity-motivated learning, our results suggest that the IS-RSA could be caused by similar brain activity related to the context-based prediction errors and surprise triggered by the violation of expectation in magic tricks as a state of uncertainty, creating a salient signal that could be mirrored in the curiosity ratings and influence encoding. Importantly, the AIC might also be a hub where dopaminergic and noradrenergic signals converge, potentially supporting curiosity-motivated learning. Future research is needed to determine whether similar processes also apply in the context of information-based prediction errors within the trivia question paradigm.

**Intersubject Functional Connectivity (ISFC)**

We conducted exploratory ISFC analysis using aHPC and VTA/SN as seed regions. There were a few observations. First, we found significant ISFC between aHPC and VTA/SN during magic trick watching. Second, both seeds showed widespread seed-based ISFC within similar cortical and subcortical areas including the striatum and thalamus, while the ISFC of the VTA/SN seed seems to descriptively span across larger regions. Overall, both seeds were positively correlated across subjects with the DMN, VAN, FPN, medial Vis, Limbic, and Somatosensory Network and negatively correlated with the lateral Vis and DAN. Given that previous studies have linked the DAN and Vis to extrinsic, exogenous systems, whereas all other networks support endogenous and exogenous processing, the pattern observed here suggests that aHPC and VTA/SN belong to the latter category.

However, our further analysis showed that this pattern was modulated by some behavioural indices. For example, similarity in curiosity and memory predicted similarity in seed-based ISFC with the dopaminergic midbrain in various regions belonging predominantly to the DMN or FPN. One interesting finding was that ISFC-RSA memory effects were also found in the left CN, overlapping with the cluster where the ISC-RSA memory effect was found in the ROI analysis, suggesting that the effect could at least partly have been driven by dopamine. In addition, similarity in CMLE predicted similarity in whole-brain ISFC for the aHPC and the VTA/SN seed, resulting in positive and negative clusters. The ISFC-RSA CMLE effects were highly similar across both seeds. Positive clusters were located in the Vis around the occipital pole partly stretching laterally as well as in the DAN (e.g., posterior superior parietal cortex and superior frontal gyrus) bordering the Somatosensory network (pre- and postcentral gyrus). Negative clusters were found in the DMN (e.g., ACC, PCC, angular gyrus), FPN (e.g., MFG, inferior temporal gyrus), VAN (e.g., aI/fO), Vis medially (e.g., cuneus) as well as in the Limbic network (e.g., temporal pole, entorhinal and perirhinal cortex) and subcortical regions including the striatum and thalamus. Importantly using the aHPC seed, voxels showing negative ISFC-RSA CMLE effects in the VTA/SN were found, and vice versa. These results indicate that, while aHPC and VTA/SN show positive correlations with endogenous systems during naturalistic viewing, in the context of curiosity-motivated learning, the communication of both seeds and other members of the endogenous systems (e.g., in the DMN, FPN, or Limbic network) becomes more variable and less stimulus-driven in participants with high compared to low CMLE. In comparison, the communication between both seeds and members of the exogenous system (DAN and Vis) becomes more similar for participants with high compared to low CMLE scores. Overall, this suggests that CMLE is supported by stimulus-driven communication between the aHPC and VTA/SN seed and predominantly exogenous regions and unique ISFC patterns with more endogenous systems.

One critical factor that helps us explain these results are prediction errors. Magic tricks violate expectations, predictions, and cause-effect relationships. Such (unsigned) prediction errors often create a sense of surprise (Antony et al., 2021; O'Reilly et al., 2013). Surprise, in turn, can trigger curiosity (Lau et al., 2020; Ligneul et al., 2018; Ozono et al., 2021; Vogl et al., 2019). Importantly, the HPC does not only represent space and context in form from cognitive maps (O'Keefe & Nadel, 1978), but also more generally predictive maps of future states, encoding expectations thereof to support subsequent learning (Stachenfeld et al., 2017). As such, the HPC plays a role in information processing by weighing expectations and novel evidence (Rigoli et al., 2019) and in the detection and encoding of unexpected events (Axmacher et al., 2010). Some also suggest that the HPC signals uncertainty or predictability of events (Harrison et al., 2006) reflecting a more generic context-sensitivity to the probabilistic structure of the environment observed events. A prediction error also signals saliency, warning that a belief update is necessary to improve the internal model and future predictions. This saliency is signalled by a dopaminergic response and research suggests that this is not only the case for reward prediction errors, but prediction errors in general (Antony et al., 2021; Horvitz, 2000; Pine et al., 2018; Ungless, 2004). Therefore, the ISFC between aHPC and VTA/SN may be explained by the common stimulus-induced responses that signal salient information in the context of uncertainty and PEs elicited by the magic tricks, replicating previous results of increased FC between both brain regions in the context of unexpected, salient single images (Murty & Adcock, 2014).

Similarly, the results of the ISFC-RSA of curiosity and memory using the VTA as seed could reflect the interplay between the prediction error related dopaminergic response of the VTA, curiosity and memory during naturalistic viewing. In both cases, parts of the VAN (more specifically, aI/fO) are activated, assumed to play a major role in processing salient events (Corbetta et al., 2008; Corbetta & Shulman, 2002; Menon & Uddin, 2010). The salience-related signal could further recruit FPN as an adaptive control mechanism to ensure cognitive flexibility in the context of prediction error processing. Indeed, the fronto-insular cortex, where the effects found here are located, is involved in switching between the FPN and the DMN (Sridharan et al., 2008). Likewise, the DMN recruitment could reflect activity associated with the update of internal models. In each case, it seems reasonable to assume that these networks hence anchor idiosyncratic patterns of curiosity and memory, respectively.

We found that curiosity-motivated learning (as the difference in similarity in ISFC between participants with a high CMLE compared to a low CMLE score) is supported by more individualised, endogenous patterns in ISFC in response to the stimuli between both seeds and the DMN, FPN, and VAN, respectively, but more exogenous responses in the

DAN. These findings can be contrasted and interpreted with the findings by Murphy, Ranganath, and Gruber (2021). Using Psychophysiological Interaction (PPI) analysis, the authors showed that coupling between HPC and subparts of the DMN during the relief, but not the elicitation, of curiosity supports curiosity-enhanced learning. Taking a more fine-grained approach, they further showed that during relief, the HPC correlates with the medial PFC and the VTA/SN with the PCC to support curiosity-motivated learning. However, no FC between the FPN and the subcortical structures (HPC and VTA/SN) was found to support curiosity-motivated learning, whereas FC between DMN and FPN during both elicitation and relief supported curiosity-motivated learning.

Intriguingly, in conjunction with the results by Murphy and colleagues (2021), we also identify ISFC between aHPC and medial PFC as well as between the VTA/SN and the PCC as neural substrates of CMLE. Our results extend their findings by showing that the stimulus-induced communication between subcortical and cortical areas in support of CMLE is individualised and endogenous. While subcortical structures and FPN did not show an interaction between curiosity and memory in the PPI analysis, our results suggest that participants with high CMLE scores show more individualised, endogenous ISFC patterns between the FPN and both seeds in response to the stimuli compared to participants with low scores. This might reflect the engagement of cognitive appraisal and control mechanisms in support of the ongoing information-seeking following the encounter of a prediction error (Gruber & Ranganath, 2019). Lastly, while Murphy et al. (2021) did not include the VAN and DAN, we found that ISFC patterns for both seeds were more exogenous and stimulus-driven in the DAN, but more endogenous and variable in the VAN in support of curiosity-motivated learning. Assuming that stimulus-induced responses in the aHPC and VTA/SN are indeed related to signalling salient information in the context of uncertainty and PEs, this suggests that salience and bottom-up attentional processes in the VAN are signalled in an endogenous, variable manner across participants in response to the stimulus, whereas top-down attentional processes to redirect attention in response to the unexpected events in DAN are more exogenous and stimulus-driven to support curiosity-motivated learning.

Taken together, the results of the ISFC analysis suggest that while the time course of aHPC and VTA/SN synchronise with the response in large cortical networks across subjects when presented with magic tricks, these patterns of FC overall flip to support curiosity-motivated learning where higher CMLE scores are associated with more endogenous, unique ISFC patterns, not only between both seeds but also to cortical networks. As unexpected events and prediction errors are appraised in a way eliciting curiosity, curiosity in turn enhances encoding by individualised ISFC patterns between subcortical and cortical areas. As such, curiosity-motivated learning is potentially related to or dependent on the high-order response to unexpected events described by Brandmann and colleagues (2021) in the context of surprise, but extends beyond that by further also recruiting ISFC connections between VTA/SN and aHPC and FPN, VAN, and DAN, respectively, partly by stimulus-induced synchronisation of ISFC patterns across participants, but predominantly partly by stimulus-induced de-synchronisation across participants.

### Supplementary References

Antony, J. W., Hartshorne, T. H., Pomeroy, K., Gureckis, T. M., Hasson, U., McDougle, S. D., & Norman, K. A. (2021). Behavioral, Physiological, and Neural Signatures of Surprise during Naturalistic Sports Viewing. *Neuron*, *109*(2), 377–390.e7.

Axmacher, N., Cohen, M. X., Fell, J., Haupt, S., Dümpelmann, M., Elger, C. E., Schlaepfer, T. E., Lenartz, D., Sturm, V., & Ranganath, C. (2010). Intracranial EEG Correlates of Expectancy and Memory Formation in the Human Hippocampus and Nucleus Accumbens. *Neuron*, *65*(4), 541–549.

Baranes, A., Oudeyer, P. Y., & Gottlieb, J. (2015). Eye movements reveal epistemic curiosity in human observers. *Vision Research*, *117*, 81–90.

Bartra, O., McGuire, J. T., & Kable, J. W. (2013). The valuation system: a coordinate-based meta-analysis of BOLD fMRI experiments examining neural correlates of subjective value. *NeuroImage*, *76*, 412–427.

Bastin, C., Besson, G., Simon, J., Delhaye, E., Geurten, M., Willems, S., & Salmon, E. (2019). An integrative memory model of recollection and familiarity to understand memory deficits. *The Behavioral and Brain Sciences*, *42*, e281.

Berridge, K. C. (2000). Reward learning: Reinforcement, incentives, and expectations. In *Psychology of Learning and Motivation* (Vol. 40, pp. 223–278). Academic Press.

Bisley, J. W., & Goldberg, M. E. (2010). Attention, intention, and priority in the parietal lobe. *Annual Review of Neuroscience*, *33*, 1–21.

Biswal, B., Yetkin, F. Z., Haughton, V. M., & Hyde, J. S. (1995). Functional connectivity in the motor cortex of resting human brain using echo-planar MRI. *Magnetic Resonance in Medicine: Official Journal of the Society of Magnetic Resonance in Medicine / Society of Magnetic Resonance in Medicine*, *34*(4), 537–541.

Blumenfeld, R. S., & Ranganath, C. (2007). Prefrontal cortex and long-term memory encoding: an integrative review of findings from neuropsychology and neuroimaging. *The Neuroscientist: A Review Journal Bringing Neurobiology, Neurology and Psychiatry*, *13*(3), 280–291.

Brandman, T., Malach, R., & Simony, E. (2021). The surprising role of the default mode network in naturalistic perception. *Communications Biology*, *4*(1), 79.

Bromberg-Martin, E. S., Matsumoto, M., & Hikosaka, O. (2010). Dopamine in motivational control: rewarding, aversive, and alerting. *Neuron*, *68*(5), 815–834.

Cabeza, R., Ciaramelli, E., Olson, I. R., & Moscovitch, M. (2008). The parietal cortex and episodic memory: an attentional account. *Nature Reviews. Neuroscience*, *9*(8), 613–625.

Cabeza, R., Prince, S. E., Daselaar, S. M., Greenberg, D. L., Budde, M., Dolcos, F., LaBar, K. S., & Rubin, D. C. (2004). Brain activity during episodic retrieval of autobiographical and laboratory events: an fMRI study using a novel photo paradigm. *Journal of Cognitive Neuroscience*, *16*(9), 1583–1594.

Chang, L. J., Yarkoni, T., Khaw, M. W., & Sanfey, A. G. (2012). Decoding the Role of the Insula in Human Cognition: Functional Parcellation and Large-Scale Reverse Inference. *Cerebral Cortex*, *23*(3), 739–749.

Chen, J., Leong, Y. C., Honey, C. J., Yong, C. H., Norman, K. A., & Hasson, U. (2017). Shared memories reveal shared structure in neural activity across individuals. *Nature Neuroscience*, *20*(1), 115–125.

Cheung, V. K. M., Harrison, P. M. C., Meyer, L., Pearce, M. T., Haynes, J.-D., & Koelsch, S. (2019). Uncertainty and Surprise Jointly Predict Musical Pleasure and Amygdala, Hippocampus, and Auditory Cortex Activity. *Current Biology: CB*, *29*(23), 4084–4092.e4.

Chikama, M., McFarland, N. R., Amaral, D. G., & Haber, S. N. (1997). Insular cortical projections to functional regions of the striatum correlate with cortical cytoarchitectonic organization in the primate. *The Journal of Neuroscience: The Official Journal of the Society for Neuroscience*, *17*(24), 9686–9705.

Chong, T. T.-J., Cunnington, R., Williams, M. A., Kanwisher, N., & Mattingley, J. B. (2008). fMRI adaptation reveals mirror neurons in human inferior parietal cortex. *Current Biology: CB*, *18*(20), 1576–1580.

Cho, Y. T., Fromm, S., Guyer, A. E., Detloff, A., Pine, D. S., Fudge, J. L., & Ernst, M. (2013). Nucleus accumbens, thalamus and insula connectivity during incentive anticipation in typical adults and adolescents. *NeuroImage*, *66*, 508–521.

Clark, A. (2013). Whatever next? Predictive brains, situated agents, and the future of cognitive science. *The Behavioral and Brain Sciences*, *36*(3), 181–204.

Cohen, N., Ben-Yakov, A., Weber, J., Edelson, M. G., Paz, R., & Dudai, Y. (2020). Prestimulus Activity in the Cingulo-Opercular Network Predicts Memory for Naturalistic Episodic Experience. *Cerebral Cortex*, *30*(3), 1902–1913.

Cole, M. W., Reynolds, J. R., Power, J. D., Repovs, G., Anticevic, A., & Braver, T. S. (2013). Multi-task connectivity reveals flexible hubs for adaptive task control. *Nature Neuroscience*, *16*(9), 1348–1355.

Corbetta, M., Patel, G., & Shulman, G. L. (2008). The Reorienting System of the Human Brain: From Environment to Theory of Mind. *Neuron*, *58*(3), 306–324.

Corbetta, M., & Shulman, G. L. (2002). Control of goal-directed and stimulus-driven attention in the brain. *Nature Reviews. Neuroscience*, *3*(3), 201–215.

Craig, A. D. B. (2009). How do you feel--now? The anterior insula and human awareness. *Nature Reviews. Neuroscience*, *10*(1), 59–70.

Danek, A. H., Öllinger, M., Fraps, T., Grothe, B., & Flanagin, V. L. (2015). An fMRI investigation of expectation violation in magic tricks. *Frontiers in Psychology*, *6*, 84.

Deci, E. L., & Ryan, R. M. (1985). *Intrinsic Motivation and Self-Determination in Human Behavior*. Springer, Boston, MA.

Di Domenico, S. I., & Ryan, R. M. (2017). The Emerging Neuroscience of Intrinsic Motivation: A New Frontier in Self-Determination Research. *Frontiers in Human Neuroscience*, *11*, 145.

Diekhof, E. K., Kaps, L., Falkai, P., & Gruber, O. (2012). The role of the human ventral striatum and the medial orbitofrontal cortex in the representation of reward magnitude - An activation likelihood estimation meta-analysis of neuroimaging studies of passive reward expectancy and outcome processing. *Neuropsychologia*, *50*(7), 1252–1266.

Dinstein, I., Hasson, U., Rubin, N., & Heeger, D. J. (2007). Brain areas selective for both observed and executed movements. *Journal of Neurophysiology*, *98*(3), 1415–1427.

Dosenbach, N. U. F., Fair, D. A., Miezin, F. M., Cohen, A. L., Wenger, K. K., Dosenbach, R. A. T., Fox, M. D., Snyder, A. Z., Vincent, J. L., Raichle, M. E., Schlaggar, B. L., & Petersen, S. E. (2007). Distinct brain networks for adaptive and stable task control in humans. *Proceedings of the National Academy of Sciences of the United States of America*, *104*(26), 11073–11078.

Dosenbach, N. U. F., Visscher, K. M., Palmer, E. D., Miezin, F. M., Wenger, K. K., Kang, H. C., Burgund, E. D., Grimes, A. L., Schlaggar, B. L., & Petersen, S. E. (2006). A Core System for the Implementation of Task Sets. *Neuron*, *50*(5), 799–812.

Droutman, V., Bechara, A., & Read, S. J. (2015). Roles of the Different Sub-Regions of the Insular Cortex in Various Phases of the Decision-Making Process. *Frontiers in Behavioral Neuroscience*, *9*, 309.

Duan, H., Fernández, G., van Dongen, E., & Kohn, N. (2020). The effect of intrinsic and extrinsic motivation on memory formation: insight from behavioral and imaging study. *Brain Structure & Function*, *225*(5), 1561–1574.

Fanselow, M. S., & Dong, H.-W. (2010). Are the dorsal and ventral hippocampus functionally distinct structures? *Neuron*, *65*(1), 7–19.

Flynn, F. G. (1999). Anatomy of the insula functional and clinical correlates. *Aphasiology*, *13*(1), 55–78.

Friston, K. (2010). The free-energy principle: a unified brain theory? *Nature Reviews. Neuroscience*, *11*(2), 127–138.

Friston, K., Thornton, C., & Clark, A. (2012). Free-energy minimization and the dark-room problem. *Frontiers in Psychology*, *3*, 130.

Fuster, J. M. (1997). Network memory. *Trends in Neurosciences*, *20*(10), 451–459.

Gershman, S. J., & Uchida, N. (2019). Believing in dopamine. *Nature Reviews. Neuroscience*, *20*(11), 703–714.

Glasser, M. F., Coalson, T. S., Robinson, E. C., Hacker, C. D., Harwell, J., Yacoub, E., Ugurbil, K., Andersson, J., Beckmann, C. F., Jenkinson, M., Smith, S. M., & Van Essen, D. C. (2016). A multi-modal parcellation of human cerebral cortex. *Nature*, *536*(7615), 171–178.

Grinband, J., Hirsch, J., & Ferrera, V. P. (2006). A neural representation of categorization uncertainty in the human brain. *Neuron*, *49*(5), 757–763.

Gruber, M. J., Gelman, B. D., & Ranganath, C. (2014). States of curiosity modulate hippocampus-dependent learning via the dopaminergic circuit. *Neuron*, *84*(2), 486–496.

Gruber, M. J., & Ranganath, C. (2019). How Curiosity Enhances Hippocampus-Dependent Memory: The Prediction, Appraisal, Curiosity, and Exploration (PACE) Framework. *Trends in Cognitive Sciences*, *23*(12), 1014–1025.

Gruber, M. J., Ritchey, M., Wang, S. F., Doss, M. K., & Ranganath, C. (2016). Post-learning Hippocampal Dynamics Promote Preferential Retention of Rewarding Events. *Neuron*, *89*(5), 1110–1120.

Harrison, L. M., Duggins, A., & Friston, K. J. (2006). Encoding uncertainty in the hippocampus. *Neural Networks: The Official Journal of the International Neural Network Society*, *19*(5), 535–546.

Hasson, U., Chen, J., & Honey, C. J. (2015). Hierarchical process memory: Memory as an integral component of information processing. *Trends in Cognitive Sciences*, *19*(6), 304–313.

Hasson, U., Nir, Y., Levy, I., Fuhrmann, G., & Malach, R. (2004). Intersubject Synchronisation of Cortical Activity During Natural Vision. *Science*, *303*(5664), 1634–1640.

Hauser, T. U., Eldar, E., Purg, N., Moutoussis, M., & Dolan, R. J. (2019). Distinct Roles of Dopamine and Noradrenaline in Incidental Memory. *The Journal of Neuroscience: The Official Journal of the Society for Neuroscience*, *39*(39), 7715–7721.

Hermans, E. J., Van Marle, H. J. F., Ossewaarde, L., Henckens, M. J. A. G., Qin, S., Van Kesteren, M. T. R., Schoots, V. C., Cousijn, H., Rijpkema, M., Oostenveld, R., & Fernández, G. (2011). Stress-related noradrenergic activity prompts large-scale neural network reconfiguration. *Science*, *334*(6059), 1151–1153.

Horvitz, J. C. (2000). Mesolimbocortical and nigrostriatal dopamine responses to salient non-reward events. *Neuroscience*, *96*(4), 651–656.

Huang, Z., Tarnal, V., Vlisides, P. E., Janke, E. L., McKinney, A. M., Picton, P., Mashour, G. A., & Hudetz, A. G. (2021). Anterior insula regulates brain network transitions that gate conscious access. *Cell Reports*, *35*(5), 109081.

Huettel, S. A., Song, A. W., & McCarthy, G. (2005). Decisions under uncertainty: probabilistic context influences activation of prefrontal and parietal cortices. *The Journal of Neuroscience: The Official Journal of the Society for Neuroscience*, *25*(13), 3304–3311.

Huettel, S. A., Stowe, C. J., Gordon, E. M., Warner, B. T., & Platt, M. L. (2006). Neural signatures of economic preferences for risk and ambiguity. *Neuron*, *49*(5), 765–775.

Ishida, H., Nakajima, K., Inase, M., & Murata, A. (2010). Shared mapping of own and others' bodies in visuotactile bimodal area of monkey parietal cortex. *Journal of Cognitive Neuroscience*, *22*(1), 83–96.

Jepma, M., Verdonschot, R. G., van Steenbergen, H., Rombouts, S. A. R. B., & Nieuwenhuis, S. (2012). Neural mechanisms underlying the induction and relief of perceptual curiosity. *Frontiers in Behavioral Neuroscience*, *6*, 5.

Kang, M. J., Hsu, M., Krajbich, I. M., Loewenstein, G., McClure, S. M., Wang, J. T.-Y., & Camerer, C. F. (2009). The wick in the candle of learning: epistemic curiosity activates reward circuitry and enhances memory. *Psychological Science*, *20*(8), 963–973.

Kempadoo, K. A., Mosharov, E. V., Choi, S. J., Sulzer, D., & Kandel, E. R. (2016). Dopamine release from the locus coeruleus to the dorsal hippocampus promotes spatial learning and memory. *Proceedings of the National Academy of Sciences of the United States of America*, *113*(51), 14835–14840.

Kim, H. (2010). Dissociating the roles of the default-mode, dorsal, and ventral networks in episodic memory retrieval. *NeuroImage*, *50*(4), 1648–1657.

Kim, H. (2011). Neural activity that predicts subsequent memory and forgetting: a meta-analysis of 74 fMRI studies. *NeuroImage*, *54*(3), 2446–2461.

Knutson, B., Adams, C. M., Fong, G. W., & Hommer, D. (2001). Anticipation of increasing monetary reward selectively recruits nucleus accumbens. *The Journal of Neuroscience: The Official Journal of the Society for Neuroscience*, *21*(16), RC159.

Lau, J. K. L., Ozono, H., Kuratomi, K., Komiya, A., & Murayama, K. (2020). Shared striatal activity in decisions to satisfy curiosity and hunger at the risk of electric shocks. *Nature Human Behaviour*, *4*(5), 531–543.

Lawson, R. P., Bisby, J., Nord, C. L., Burgess, N., & Rees, G. (2021). The Computational, Pharmacological, and Physiological Determinants of Sensory Learning under Uncertainty. *Current Biology: CB*, *31*(1), 163–172.e4.

Lee, W. (2016). Insular Cortex Activity as the Neural Base of Intrinsic Motivation. In *Recent Developments in Neuroscience Research on Human Motivation* (Vol. 19, pp. 127–148). Emerald Group Publishing Limited.

Lee, W., & Reeve, J. (2013). Self-determined, but not non-self-determined, motivation predicts activations in the anterior insular cortex: an fMRI study of personal agency. *Social Cognitive and Affective Neuroscience*, *8*(5), 538–545.

Lee, W., & Reeve, J. (2017). Identifying the neural substrates of intrinsic motivation during task performance. *Cognitive, Affective & Behavioral Neuroscience*, *17*(5), 939–953.

Lee, W., Reeve, J., Xue, Y., & Xiong, J. (2012). Neural differences between intrinsic reasons for doing versus extrinsic reasons for doing: an fMRI study. *Neuroscience Research*, *73*(1), 68–72.

Ligneul, R., Mermillod, M., & Morisseau, T. (2018). From relief to surprise: Dual control of epistemic curiosity in the human brain. *NeuroImage*, *181*, 490–500.

Lisman, J. E., Grace, A. A., & Duzel, E. (2011). A neoHebbian framework for episodic memory; role of dopamine-dependent late LTP. *Trends in Neurosciences*, *34*(10), 536–547.

Liu, X., Hairston, J., Schrier, M., & Fan, J. (2011). Common and distinct networks underlying reward valence and processing stages: A meta-analysis of functional neuroimaging studies. *Neuroscience and Biobehavioral Reviews*, *35*(5), 1219–1236.

Loued-Khenissi, L., Pfeuffer, A., Einhäuser, W., & Preuschoff, K. (2020). Anterior insula reflects surprise in value-based decision-making and perception. *NeuroImage*, *210*, 116549.

Marvin, C. B., & Shohamy, D. (2016). Curiosity and reward: Valence predicts choice and information prediction errors enhance learning. *Journal of Experimental Psychology. General*, *145*(3), 266–272.

Mayer, J. S., Bittner, R. A., Nikolić, D., Bledowski, C., Goebel, R., & Linden, D. E. J. (2007). Common neural substrates for visual working memory and attention. *NeuroImage*, *36*(2), 441–453.

Meliss, S., Pascua, C., Skipper, J. I., & Murayama, K. (2022). *The Magic, Memory, and Curiosity fMRI Dataset of People Viewing Magic Tricks*. https://doi.org/10.31234/osf.io/zq7gv

Menon, V., & Uddin, L. Q. (2010). Saliency, switching, attention and control: a network model of insula function. *Brain Structure & Function*, *214*(5-6), 655–667.

Murayama, K., FitzGibbon, L., & Sakaki, M. (2019). Process Account of Curiosity and Interest: A Reward-Learning Perspective. *Educational Psychology Review*, *31*(4), 875–895.

Murphy, C., Ranganath, C., & Gruber, M. J. (2021). Connectivity between the hippocampus and default mode network during the relief – but not elicitation – of curiosity supports curiosity-enhanced memory enhancements. In *bioRxiv* (p. 2021.07.26.453739). https://doi.org/10.1101/2021.07.26.453739

Murty, V. P., & Adcock, R. A. (2014). Enriched encoding: reward motivation organizes cortical networks for hippocampal detection of unexpected events. *Cerebral Cortex*, *24*(8), 2160–2168.

Naqvi, N. H., & Bechara, A. (2010). The insula and drug addiction: an interoceptive view of pleasure, urges, and decision-making. *Brain Structure & Function*, *214*(5-6), 435–450.

Nastase, S. A., Gazzola, V., Hasson, U., & Keysers, C. (2019). Measuring shared responses across subjects using intersubject correlation. *Social Cognitive and Affective*

*Neuroscience*, *14*(6), 669–687.

O'Doherty, J. P., Dayan, P., Friston, K., Critchley, H., & Dolan, R. J. (2003). Temporal difference models and reward-related learning in the human brain. *Neuron*, *38*(2), 329–337.

O'Keefe, J., & Nadel, L. (1978). *The hippocampus as a cognitive map*. Oxford University Press.

O'Reilly, J. X., Schüffelgen, U., Cuell, S. F., Behrens, T. E. J., Mars, R. B., & Rushworth, M. F. S. (2013). Dissociable effects of surprise and model update in parietal and anterior cingulate cortex. *Proceedings of the National Academy of Sciences of the United States of America*, *110*(38), E3660–E3669.

Ozono, H., Komiya, A., Kuratomi, K., Hatano, A., Fastrich, G., Raw, J. A. L., Haffey, A., Meliss, S., Lau, J. K. L., & Murayama, K. (2021). Magic Curiosity Arousing Tricks (MagicCATs): A novel stimulus collection to induce epistemic emotions. *Behavior Research Methods*, *53*(1), 188–215.

Parris, B. A., Kuhn, G., Mizon, G. A., Benattayallah, A., & Hodgson, T. L. (2009). Imaging the impossible: an fMRI study of impossible causal relationships in magic tricks. *NeuroImage*, *45*(3), 1033–1039.

Paulus, M. P., Rogalsky, C., Simmons, A., Feinstein, J. S., & Stein, M. B. (2003). Increased activation in the right insula during risk-taking decision making is related to harm avoidance and neuroticism. *NeuroImage*, *19*(4), 1439–1448.

Pine, A., Sadeh, N., Ben-Yakov, A., Dudai, Y., & Mendelsohn, A. (2018). Knowledge acquisition is governed by striatal prediction errors. *Nature Communications*, *9*(1), 1–14.

Platt, M. L., & Huettel, S. A. (2008). Risky business: the neuroeconomics of decision making under uncertainty. *Nature Neuroscience*, *11*(4), 398–403.

Power, J. D., Barnes, K. A., Snyder, A. Z., Schlaggar, B. L., & Petersen, S. E. (2012).

Spurious but systematic correlations in functional connectivity MRI networks arise from subject motion. *NeuroImage*, *59*(3), 2142–2154.

Preuschoff, K., Quartz, S. R., & Bossaerts, P. (2008). Human Insula Activation Reflects Risk Prediction Errors As Well As Risk. *The Journal of Neuroscience: The Official Journal of the Society for Neuroscience*, *28*(11), 2745–2752.

Ranganath, C., & Ritchey, M. (2012). Two cortical systems for memory-guided behaviour. *Nature Reviews. Neuroscience*, *13*(10), 713–726.

Ren, Y., Nguyen, V. T., Guo, L., & Guo, C. C. (2017). Inter-subject Functional Correlation Reveal a Hierarchical Organization of Extrinsic and Intrinsic Systems in the Brain. *Scientific Reports*, *7*(1), 1–12.

Rigoli, F., Michely, J., Friston, K. J., & Dolan, R. J. (2019). The role of the hippocampus in weighting expectations during inference under uncertainty. *Cortex; a Journal Devoted to the Study of the Nervous System and Behavior*, *115*, 1–14.

Sakaki, M., Yagi, A., & Murayama, K. (2018). Curiosity in old age: A possible key to achieving adaptive aging. *Neuroscience and Biobehavioral Reviews*, *88*, 106–116.

Schiffer, A.-M., & Schubotz, R. I. (2011). Caudate nucleus signals for breaches of expectation in a movement observation paradigm. *Frontiers in Human Neuroscience*, *5*, 38.

Schultz, W. (1998). Predictive Reward Signal of Dopamine Neurons. *Journal of Neurophysiology*, *80*(1), 1–27.

Schwartenbeck, P., Fitzgerald, T., Dolan, R. J., & Friston, K. (2013). Exploration, novelty, surprise, and free energy minimization. *Frontiers in Psychology*, *4*, 710.

Seeley, W. W. (2019). The Salience Network: A Neural System for Perceiving and Responding to Homeostatic Demands. *The Journal of Neuroscience: The Official Journal of the Society for Neuroscience*, *39*(50), 9878–9882.

Seeley, W. W., Menon, V., Schatzberg, A. F., Keller, J., Glover, G. H., Kenna, H., Reiss, A. L., & Greicius, M. D. (2007). Dissociable Intrinsic Connectivity Networks for Salience Processing and Executive Control. *The Journal of Neuroscience: The Official Journal of the Society for Neuroscience*, *27*(9), 2349–2356.

Shafiei, G., Zeighami, Y., Clark, C. A., Coull, J. T., Nagano-Saito, A., Leyton, M., Dagher, A., & Mišic, B. (2019). Dopamine Signaling Modulates the Stability and Integration of Intrinsic Brain Networks. *Cerebral Cortex*, *29*(1), 397–409.

Shimamura, A. P. (2011). Episodic retrieval and the cortical binding of relational activity. *Cognitive, Affective & Behavioral Neuroscience*, *11*(3), 277–291.

Shohamy, D., & Adcock, R. A. (2010). Dopamine and adaptive memory. *Trends in Cognitive Sciences*, *14*(10), 464–472.

Shomstein, S. (2012). Cognitive functions of the posterior parietal cortex: top-down and bottom-up attentional control. *Frontiers in Integrative Neuroscience*, *6*, 38.

Simony, E., & Chang, C. (2020). Analysis of stimulus-induced brain dynamics during naturalistic paradigms. *NeuroImage*, *216*, 116461.

Simony, E., Honey, C. J., Chen, J., Lositsky, O., Yeshurun, Y., Wiesel, A., & Hasson, U. (2016). Dynamic reconfiguration of the default mode network during narrative comprehension. *Nature Communications*, *May 2015*. https://doi.org/10.1038/ncomms12141

Singer, T., Critchley, H. D., & Preuschoff, K. (2009). A common role of insula in feelings, empathy and uncertainty. *Trends in Cognitive Sciences*, *13*(8), 334–340.

Spaniol, J., Davidson, P. S. R., Kim, A. S. N., Han, H., Moscovitch, M., & Grady, C. L. (2009). Event-related fMRI studies of episodic encoding and retrieval: meta-analyses using activation likelihood estimation. *Neuropsychologia*, *47*(8-9), 1765–1779.

Squire, L. R., Stark, C. E. L., & Clark, R. E. (2004). The medial temporal lobe. *Annual*

*Review of Neuroscience*, *27*, 279–306.

Sridharan, D., Levitin, D. J., & Menon, V. (2008). A critical role for the right fronto-insular cortex in switching between central-executive and default-mode networks. *Proceedings of the National Academy of Sciences of the United States of America*, *105*(34), 12569–12574.

Stachenfeld, K. L., Botvinick, M. M., & Gershman, S. J. (2017). The hippocampus as a predictive map. *Nature Neuroscience*, *20*(11), 1643–1653.

Sterzer, P., & Kleinschmidt, A. (2010). Anterior insula activations in perceptual paradigms: often observed but barely understood. *Brain Structure & Function*, *214*(5-6), 611–622.

Takeuchi, T., Duszkiewicz, A. J., Sonneborn, A., Spooner, P. A., Yamasaki, M., Watanabe, M., Smith, C. C., Fernández, G., Deisseroth, K., Greene, R. W., & Morris, R. G. M. (2016). Locus coeruleus and dopaminergic consolidation of everyday memory. *Nature*, *537*(7620), 357–362.

Thomas, R. M., De Sanctis, T., Gazzola, V., & Keysers, C. (2018). Where and how our brain represents the temporal structure of observed action. *NeuroImage*, *183*, 677–697.

Tobler, P. N., Fiorillo, C. D., & Schultz, W. (2005). Adaptive Coding of Reward Value by Dopamine Neurons. *Science*, *307*(5715), 1642–1645.

Totah, N. K. B., Logothetis, N. K., & Eschenko, O. (2019). Noradrenergic ensemble-based modulation of cognition over multiple timescales. *Brain Research*, *1709*, 50–66.

Ullsperger, M., Harsay, H. A., Wessel, J. R., & Ridderinkhof, K. R. (2010). Conscious perception of errors and its relation to the anterior insula. *Brain Structure & Function*, *214*(5-6), 629–643.

Ungless, M. A. (2004). Dopamine: the salient issue. *Trends in Neurosciences*, *27*(12), 702–706.

Vanderwal, T., Eilbott, J., & Castellanos, F. X. (2019). Movies in the magnet: Naturalistic paradigms in developmental functional neuroimaging. *Developmental Cognitive Neuroscience*, *36*, 100600.

Vanderwal, T., Eilbott, J., Finn, E. S., Craddock, R. C., Turnbull, A., & Castellanos, F. X. (2017). Individual differences in functional connectivity during naturalistic viewing conditions. *NeuroImage*, *157*, 521–530.

van Lieshout, L. L. F., Vandenbroucke, A. R. E., Müller, N. C. J., Cools, R., & de Lange, F. P. (2018). Induction and relief of curiosity elicit parietal and frontal activity. *The Journal of Neuroscience: The Official Journal of the Society for Neuroscience*, *38*(10), 2816–2817.

Vincent, J. L., Kahn, I., Snyder, A. Z., Raichle, M. E., & Buckner, R. L. (2008). Evidence for a frontoparietal control system revealed by intrinsic functional connectivity. *Journal of Neurophysiology*, *100*(6), 3328–3342.

Vogl, E., Pekrun, R., Murayama, K., Loderer, K., & Schubert, S. (2019). Surprise, Curiosity, and Confusion Promote Knowledge Exploration: Evidence for Robust Effects of Epistemic Emotions. *Frontiers in Psychology*, *10*, 2474.

Volz, K. G., Schubotz, R. I., & von Cramon, D. Y. (2003). Predicting events of varying probability: uncertainty investigated by fMRI. *NeuroImage*, *19*(2 Pt 1), 271–280.

Volz, K. G., Schubotz, R. I., & von Cramon, D. Y. (2004). Why am I unsure? Internal and external attributions of uncertainty dissociated by fMRI. *NeuroImage*, *21*(3), 848–857.

Volz, K. G., Schubotz, R. I., & von Cramon, D. Y. (2005). Variants of uncertainty in decision-making and their neural correlates. *Brain Research Bulletin*, *67*(5), 403–412.

Wang, L., Yu, H., Hu, J., Theeuwes, J., Gong, X., Xiang, Y., Jiang, C., & Zhou, X. (2015). Reward breaks through center-surround inhibition via anterior insula. *Human Brain Mapping*, *36*(12), 5233–5251.

Weilnhammer, V., Stuke, H., Hesselmann, G., Sterzer, P., & Schmack, K. (2017). A predictive coding account of bistable perception - a model-based fMRI study. *PLoS Computational Biology*, *13*(5), e1005536.

Wilson, R. P., Colizzi, M., Bossong, M. G., Allen, P., Kempton, M., MTAC, & Bhattacharyya, S. (2018). The Neural Substrate of Reward Anticipation in Health: A Meta-Analysis of fMRI Findings in the Monetary Incentive Delay Task. *Neuropsychology Review*, *28*(4), 496–506.

Yarkoni, T., Poldrack, R. A., Nichols, T. E., Van Essen, D. C., & Wager, T. D. (2011). Large-scale automated synthesis of human functional neuroimaging data. *Nature Methods*, *8*(8), 665–670.

Yeo, B. T. T., Krienen, F. M., Sepulcre, J., Sabuncu, M. R., Lashkari, D., Hollinshead, M., Roffman, J. L., Smoller, J. W., Zöllei, L., Polimeni, J. R., Fischl, B., Liu, H., & Buckner, R. L. (2011). The organization of the human cerebral cortex estimated by intrinsic functional connectivity. *Journal of Neurophysiology*, *106*(3), 1125–1165.

Yeshurun, Y., Nguyen, M., & Hasson, U. (2021). The default mode network: where the idiosyncratic self meets the shared social world. *Nature Reviews. Neuroscience*, *22*(3), 181–192.

Zadbood, A., Chen, J., Leong, Y. C., Norman, K. A., & Hasson, U. (2017). How We Transmit Memories to Other Brains: Constructing Shared Neural Representations Via Communication. *Cerebral Cortex*, *27*(10), 4988–5000.
